# Connectome architecture shapes large-scale cortical alterations in schizophrenia: a worldwide ENIGMA study

**DOI:** 10.1101/2023.02.12.527904

**Authors:** Foivos Georgiadis, Sara Larivière, David Glahn, L. Elliot Hong, Peter Kochunov, Bryan Mowry, Carmel Loughland, Christos Pantelis, Frans A. Henskens, Melissa J. Green, Murray J. Cairns, Patricia T Michie, Paul E. Rasser, Paul Tooney, Rodney J. Scott, Stanley Catts, Ulrich Schall, Vaughan Carr, Yann Quidé, Axel Krug, Frederike Stein, Igor Nenadić, Katharina Brosch, Tilo Kircher, Raquel Gur, Ruben Gur, Theodore D. Satterthwaite, Andriana Karuk, Edith Pomarol- Clotet, Joaquim Radua, Paola Fuentes-Claramonte, Raymond Salvador, Gianfranco Spalletta, Aristotle Voineskos, Kang Sim, Benedicto Crespo-Facorro, Diana Tordesillas Gutiérrez, Stefan Ehrlich, Nicolas Crossley, Dominik Grotegerd, Jonathan Repple, Rebekka Lencer, Udo Dannlowski, Vince Calhoun, Caroline Demro, Ian S. Ramsay, Scott R. Sponheim, Andre Schmidt, Stefan Borgwardt, Alexander S. Tomyshev, Irina Lebedeva, Cyril Hoschl, Filip Spaniel, Adrian Preda, Dana Nguyen, Anne Uhlmann, Dan J Stein, Fleur M Howells, Henk S. Temmingh, Ana M. Diaz Zuluaga, Carlos López Jaramillo, Felice Iasevoli, Ellen Ji, Stephanie Homan, Wolfgang Omlor, Philipp Homan, Stefan Kaiser, Erich Seifritz, Bratislav Misic, Sofie L. Valk, Paul Thompson, Theo G.M. van Erp, Jessica Turner, Boris Bernhardt, Matthias Kirschner

**Affiliations:** Department of Psychiatry, Psychotherapy and Psychosomatics, Psychiatric Hospital University of Zurich, Switzerland; McGill University, Montreal Neurological Institute; Department of Psychiatry, Harvard Medical School, Boston, MA USA; Maryland Psychiatric Research Center, Department of Psychiatry, University of Maryland School of Medicine; Queensland Brain Institute, The University of Queensland, QLD Australia; School of Medicine and Public Health, University of Newcastle; Melbourne Neuropsychiatry Centre, Department of Psychiatry, The University of Melbourne, Carlton South, Victoria, Australia; School of Medicine and Public Health, University of Newcastle, Australia; School of Clinical Medicine, Discipline of Psychiatry and Mental Health, UNSW Sydney, Sydney, NSW, Australia; School of Biomedical Science and Pharmacy, University of Newcastle, NSW Australia; School of Psychological Sciences, University of Newcastle, NSW Australia; School of Medicine and Public Health, College of Health, Medicine, and Wellbeing, The University of Newcastle, Callaghan, NSW, Australia; Faculty of Medicine, University of Queensland, QLD Australia; Hunter Medical Research Institute, Newcastle, Australia; School of Clinical Medicine, Discipline of Psychiatry, UNSW Sydney, Sydney, NSW, Australia; University Hospital Bonn, Department of Psychiatry and Psychotherapy, Venusberg-Campus 1, 53127 Bonn, Germany; Department of Psychiatry, University of Marburg, Rudolf Bultmann Str. 8, 35039 Marburg, Germany; Dept. of Psychiatry and Psychotherapy, Philipps-University Marburg, Marburg, Germany; University of Pennsylvania Perelman School of Medicine, Philadelphia, Pennsylvania, USA; FIDMAG Germanes Hospitalàries Research Foundation, Barcelona, Spain; Institut d’Investigacions Biomèdiques August Pi i Sunyer (IDIBAPS), Barcelona, Spain; Santa Lucia Foundation IRCCS, Rome; West Region, Institute of Mental Health, Singapore; Hospital Universitario Virgen del Rocío, IBiS-CSIC, Universidad de Sevilla, Seville, Spain; Department of Radiology, Marqués de Valdecilla University Hospital, Valdecilla Biomedical Research Institute IDIVAL, Santander, Spain; Division of Psychological & Social Medicine and Developmental Neurosciences, Technischen Universität Dresden, Faculty of Medicine, University Hospital C.G. Carus, Germany; Department of Psychiatry, Pontificia Universidad Católica de Chile; Institute for Translational Psychiatry, University of Münster, Germany; Tri-Institutional Center for Translational Research in Neuroimaging and Data Science (TReNDS), Georgia State, Georgia Tech, Emory, Atlanta, GA, USA; University of Minnesota Department of Psychology; University of Minnesota Department of Psychiatry & Behavioral Sciences; Minneapolis VA Health Care System; University of Basel, Department of Psychiatry, Basel, Switzerland; Department of Psychiatry, University of Lübeck. Germany; Mental Health Research Center, Moscow, Russian Federation; National Institute of Mental Health, Topolova 748, 250 67 Klecany, Czech Republic; Department of Psychiatry and Human Behavior, University of California Irvine, CA, USA; Department of Pediatric Neurology University of California Irvine, CA, USA; Department of child and adolescent psychiatry, TU Dresden, Germany; Department of Psychiatry and Mental Health, University of Cape Town, South Africa; Research Group in Psychiatry, Department of Psychiatry, School of Medicine, Universidad de Antioquia, Medellin, Colombia; University of Naples, Department of Neuroscience; Division of Adult Psychiatry, Department of Psychiatry, Geneva University Hospitals, Geneva, Switzerland; Forschungszentrum Jülich, Jülich, Germany; Max Planck Institute for Cognitive and Brain Sciences, Leipzig, Germany; Imaging Genetics Center, Stevens Institute for Neuroimaging and Informatics, Keck School of Medicine, University of Southern California, Los Angeles, CA, USA

## Abstract

While schizophrenia is considered a prototypical network disorder characterized by widespread brain-morphological alterations, it still remains unclear whether distributed structural alterations robustly reflect underlying network layout. Here, we tested whether large-scale structural alterations in schizophrenia relate to normative structural and functional connectome architecture, and systematically evaluated robustness and generalizability of these network-level alterations. Leveraging anatomical MRI scans from 2,439 adults with schizophrenia and 2,867 healthy controls from 26 ENIGMA sites and normative data from the Human Connectome Project (n=207), we evaluated structural alterations of schizophrenia against two network susceptibility models: i) hub vulnerability, which examines associations between regional network centrality and magnitude of disease-related alterations; ii) epicenter mapping, which identify regions whose typical connectivity profile most closely resembles the disease-related morphological alterations. To assess generalizability and specificity, we contextualized the influence of site, disease stages, and individual clinical factors and compared network associations of schizophrenia with that found in affective disorders. Schizophrenia-related structural alterations co-localized with interconnected functional and structural hubs and harbored temporo-paralimbic and frontal epicenters. Findings were robust across sites and related to individual symptom profiles. We observed localized unique epicenters for first-episode psychosis and early stages, and transmodal epicenters that were shared across first-episode to chronic stages. Moreover, transdiagnostic comparisons revealed overlapping epicenters in schizophrenia and bipolar, but not major depressive disorder, yielding insights in pathophysiological continuity within the schizophrenia-bipolar-spectrum. In sum, cortical alterations over the course of schizophrenia robustly follow brain network architecture, emphasizing marked hub susceptibility and temporo-frontal epicenters at both the level of the group and the individual. Subtle variations of epicenters across disease stages suggest interacting pathological processes, while associations with patient-specific symptoms support additional inter-individual variability of hub vulnerability and epicenters in schizophrenia. Our work contributes to recognizing potentially common pathways to better understand macroscale structural alterations, and inter-individual variability in schizophrenia.

## Introduction

Neuropsychiatric disorders including schizophrenia (SCZ), bipolar (BD) and major depressive disorders (MDD), are increasingly recognized as network disorders (1–4) characterized by widespread cortical and subcortical alterations (5–10). Among these, SCZ is associated with the most severe and distributed structural alterations (6, 11, 12) appearing in all disease stages (13), with temporal-paralimbic and frontal regions being the most affected (5, 13–15). However, the pattern is not uniformly distributed across the entire brain (15) raising the question how the brain’s connectome architecture guides the spatial distribution of subcortical and cortical alterations in SCZ (5, 6). The staggering interconnectedness of regions in the brain, which gives rise to a multi-level network architecture, suggests that local aberrant functioning may affect synaptically-connected neuronal populations (16, 17). Distributed structural alterations could, thus, arise from, or be influenced by, network architecture and manifest as a diverse set of cognitive and affective symptoms. Recent studies have found that patterns of cortical alterations in SCZ, when compared to controls, reflect distinct functional and cytoarchitectural systems (14, 18), as well as cortical connectivity (10). This suggests that SCZ-related cortical alterations are not randomly distributed, but shaped by the underlying network architecture of the cortex.

Notably, there are important inter-regional variations in connectivity, with hub regions showing denser connectivity patterns to other regions (19, 20). These regions may serve as central information relays, promoting processing efficiency (1). High centrality of a region, however, may render it more susceptible to metabolic demands, potentially predisposing it to pronounced structural alterations— termed the nodal stress or hub vulnerability hypothesis. (1, 21, 22). Consistent with this, higher vulnerability of hubs to structural damage has been reported across multiple neurological and neurodegenerative disorders (20, 22–26) and more recently in SCZ (2, 14, 15, 27). In addition, regional pathological processes might propagate from so-called “disease epicenters” to connected brain regions leading to network-spreading patterns of morphological alterations. Emerging evidence from neuropsychiatric and cross-disorder studies suggest that such disease epicenters can be revealed by identifying those brain regions whose connectivity profile most closely resembles a disease-related morphological alteration pattern (14, 22, 26, 28). In schizophrenia, paralimbic and frontal regions have been recently identified as the most likely epicenters, by comparing structural alteration pattern in chronic stages to neurotypical brain connectivity information (9). An important next step is to test whether associations between normative network features and the structural alteration in SCZ are robust and replicable using multisite imaging data and standardized methods (29, 30) and how they relate to different disease stages, and individual clinical profiles. In addition, it remains an open question how patterns of hub vulnerability and epicenters differ between major psychiatric disorders along the schizophrenia-bipolar-spectrum.

The Enhancing Neuro Imaging Genetics through Meta Analysis (ENIGMA) consortium, has aggregated neuroimaging data of disease phenotypes (31) in thousands of patients and controls, enabling robust inquiries into morphological alterations across sites, disease stages and individual clinical factors. Here, we used two network-based models combining MRI-based morphology data of 2,439 adults with SCZ and 2,867 healthy controls (HC) from the ENIGMA-Schizophrenia consortium and high definition structural and functional normative connectivity data from the HCP (n=207) (32). We tested the hypothesis that network-central brain regions (*i.e.,* hubs) are more vulnerable to morphometric alterations in SCZ and that individual brain regions’ connectivity profiles shape the spatial distribution of cortical alterations in SCZ. First, we applied hub vulnerability models to spatially correlate regional network centrality with the degree of SCZ-related cortical thickness and subcortical volume alterations derived from case-control comparisons. Second, we leveraged epicenter models to systematically investigate the influence of every brain region’s connectivity profile to the spatial distribution of SCZ-related cortical thickness alterations. To assess generalizability and specificity of these network associations, we contextualized the influence of site, disease stage of SCZ, and individual clinical profiles. Finally, we examined whether our network models would reflect SCZ-specific features or rather represent a shared signature across other major psychiatric conditions (33) such as has been shown for subcortical volume, cortical thickness, surface area measures (34–36), structural covariance (37), and connectome properties (3). Therefore, we extended our analysis to meta-analytic case-control alterations in BD and MDD. Both disorders show various degrees of clinical, genetic, and neuroanatomic proximity to SCZ, with BD being more proximal (33, 38–42) and MDD being more distant with respect to the genetic, clinical, and morphometric patterns of SCZ (34, 35, 39).

## Methods

The overall workflow including the multi-source data integration and the different levels of analysis are illustrated in Figure 1.

**Figure 1.**
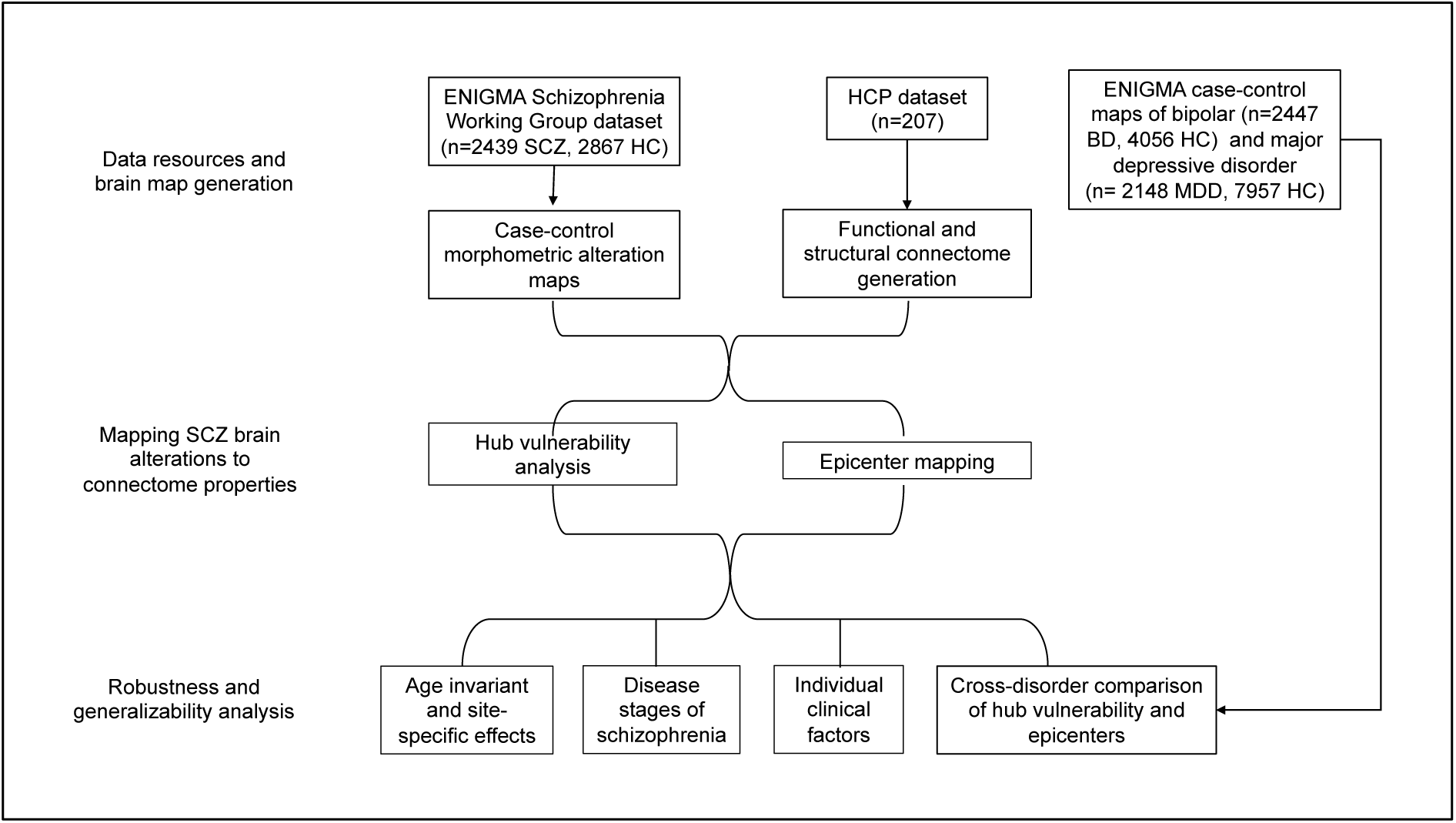
Flowchart of multi-source data integration and analysis steps.

### ENIGMA participants

We analyzed subject-level data from 26 international centers of the ENIGMA-SCZ Working Group including 2,439 adults with SCZ (786 females, mean age ± SD = 35.5±12.4 years) and 2,867 healthy controls (1,408 females, mean age ± SD = 35.0±13.4 years). Diagnosis of SCZ or schizoaffective disorder was confirmed at each center using the international diagnostic criteria of the ICD-10 (or DSM-IV). Details on subject inclusion criteria and site-specific demographic information are provided in Tables S1-2. Local institutional review boards and ethics committees approved each cohort study included, and written informed consent was provided according to local requirements.

### Image acquisition and processing

Following published ENIGMA pipelines (5, 6), all 26 participating sites processed 3D T1-weighted structural brain MRI scans using FreeSurfer (43, 44) and extracted cortical thickness (CT) from 68 Desikan-Killiany (DK) atlas regions (45) as well as subcortical volume (SV) data from 16 brain structures. Number of scanners, vendor, strength, sequence, acquisition parameters, and FreeSurfer versions are provided in Table S3. Quality control followed standard ENIGMA protocols at each site before subsequent mega-analysis (http://enigma.usc.edu/protocols/imaging-protocols).

### Statistical mega-analysis

We analyzed CT measures from 68 cortical regions and SV measures from 12 subcortical gray matter regions (bilateral amygdala, caudate, nucleus accumbens, pallidum, putamen, and thalamus) and bilateral hippocampi. Data were first harmonized across scanners and sites using ComBat, a batch effect correction tool that uses a Bayesian framework to improve the stability of the parameter estimates (46, 47). Linear models were used to compare morphometric difference profiles of CT and SV in patients relative to controls, correcting for age and sex, using SurfStat, available at (https://mica-mni.github.io/surfstat/). Findings were corrected for multiple comparisons using the false discovery rate (FDR) procedure (48). To further confirm that our batch-effect correction with ComBat on the structural MRI data was successful, we performed the original linear regression with ComBat adjusted, as well as unadjusted data, and added site as an independent variable. We then compared the partial R^2^ of site as independent variable in both models. Table S4 showed that in the ComBat adjusted data any variance explained by site was removed, thereby confirming successful batch-effect correction in our mega-analysis.

### Functional and connectivity matrix generation from HCP data

Resting-state fMRI, and diffusion MRI of unrelated healthy adults (n = 207, 83 males, mean age ± SD = 28.73 ± 3.73 years, range = 22 to 36 years) from the HCP were used and underwent the initiative’s minimal preprocessing (49, 50). High-resolution functional and structural data were parcellated according to the Desikan-Killiany atlas to align with the ENIGMA-Schizophrenia dataset. Functional and structural 68×68 connectivity matrices were generated following previous work and are described in detail in the Supplement (See supplementary methods). Within these connectivity matrices, each region’s row or column corresponds to a 68×1 or 1×68 vector respectively that reflects the connectivity profile of a given region with all other brain regions.

### Hub vulnerability model

To test the hub vulnerability hypothesis, *i.e.,* that nodes with higher normative network centrality (established based on an independent sample of healthy individuals of the HCP) would display higher levels of disease-specific morphometric alterations in SCZ, we performed spatial correlations between the cortical and subcortical SCZ-related morphometric profile and normative weighted degree centrality maps (Fig.2 A-B). Weighted degree centrality was defined by the sum of all weighted cortico-cortical and subcortico-cortical connections for every region and used to identify structural and functional hub regions. Regions with a higher weighted degree centrality are denoted as hubs, compared to regions with relatively lower weighted degree centrality. To mitigate potential bias from selecting an arbitrary threshold and inadvertently excluding valuable information, the analyses were conducted on unthresholded connectivity matrices. Since correlations between brain maps are known to be inflated due to spatial autocorrelation (51, 52) we generated a null distribution using 10,000 rotated maps based on the spin permutation test (30). In the case of the subcortical map we used a similar approach, with the exception that subcortical labels were randomly shuffled as opposed to being projected onto spheres, such as in the spin test procedure (26).

**Figure 2.**
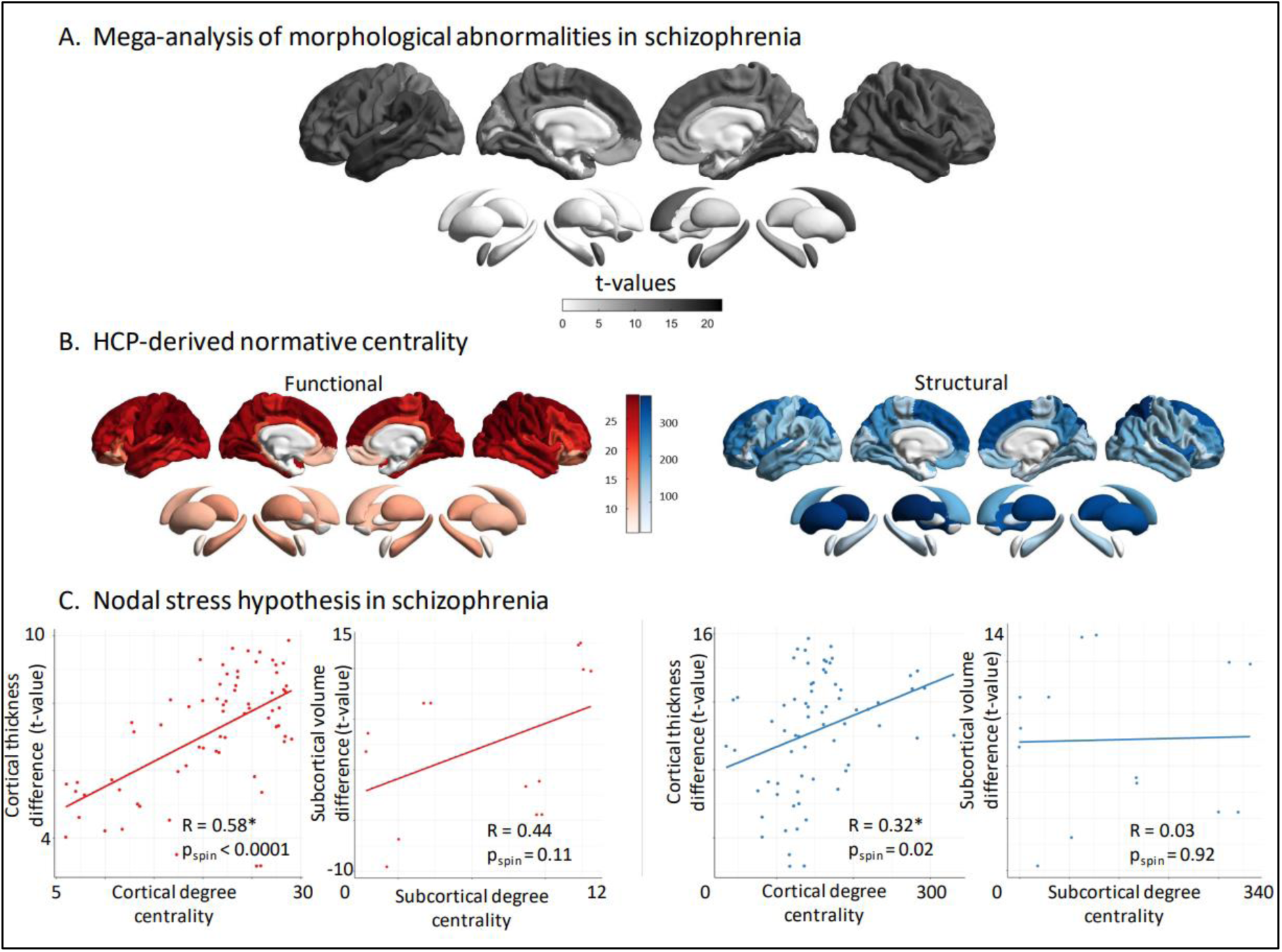
Hub vulnerability mapping. **(A)** Unthresholded *t-*maps of cortical thickness and subcortical volume deficiencies in SCZ (n = 2,439), compared to HCP (n = 2,867). **(B)** Normative functional and structural network organization, derived from the HCP dataset, was used to identify hubs (*i.e.,* regions with greater degree centrality). **(C)** Correlation of gray matter morphological alterations with node-level functional (*left*) and structural (*right*) maps of degree centrality. In SCZ, regions with high functional or structural centrality are significantly more likely to display morphological alterations in the cortex. Subcortical functional subcortico-cortical degree centrality showed a moderate non-significant correlation with morphometric alterations, while no correlation was observed for structural subcortico-cortical degree centrality. HCP: Healthy control participants; SCZ: Schizophrenia.

### Mapping disease epicenters

To identify disorder-specific cortical and subcortical epicenters, we spatially correlated every region’s healthy functional and structural cortical connectivity profile to the whole-brain patterns of cortical alterations in SCZ (Fig. 3A) (26, 30, 53). We repeated this approach systematically for each parcellated region and for functional and structural cortical connectivity separately. Statistical significance of spatial similarity between an individual brain region’s functional and structural connectivity profile and schizophrenia-related cortical alterations was determined through correlation analyses correcting for spatial autocorrelation with permutation testing as described above. Regions were ranked in descending order based on the strength of their correlation coefficients, with the highest-ranked regions being considered the most significant disease epicenters. A region can be considered an epicenter regardless of its cortical abnormality level (effect-size of case-control differences), as long as it is strongly connected to other regions with high abnormality and weakly connected to regions with low abnormality. Additionally, it is important to note that disease epicenters do not necessarily have to be hub regions but may rather be connected to them as feeder nodes that directly connect peripheral nodes to hubs. To account for multiple testing (n=68 for cortical regions and n=14 for subcortical regions), epicenters were only considered significant if spin-test p-values (p_spin_) survived Bonferroni correction (p<0.05).

**Figure 3.**
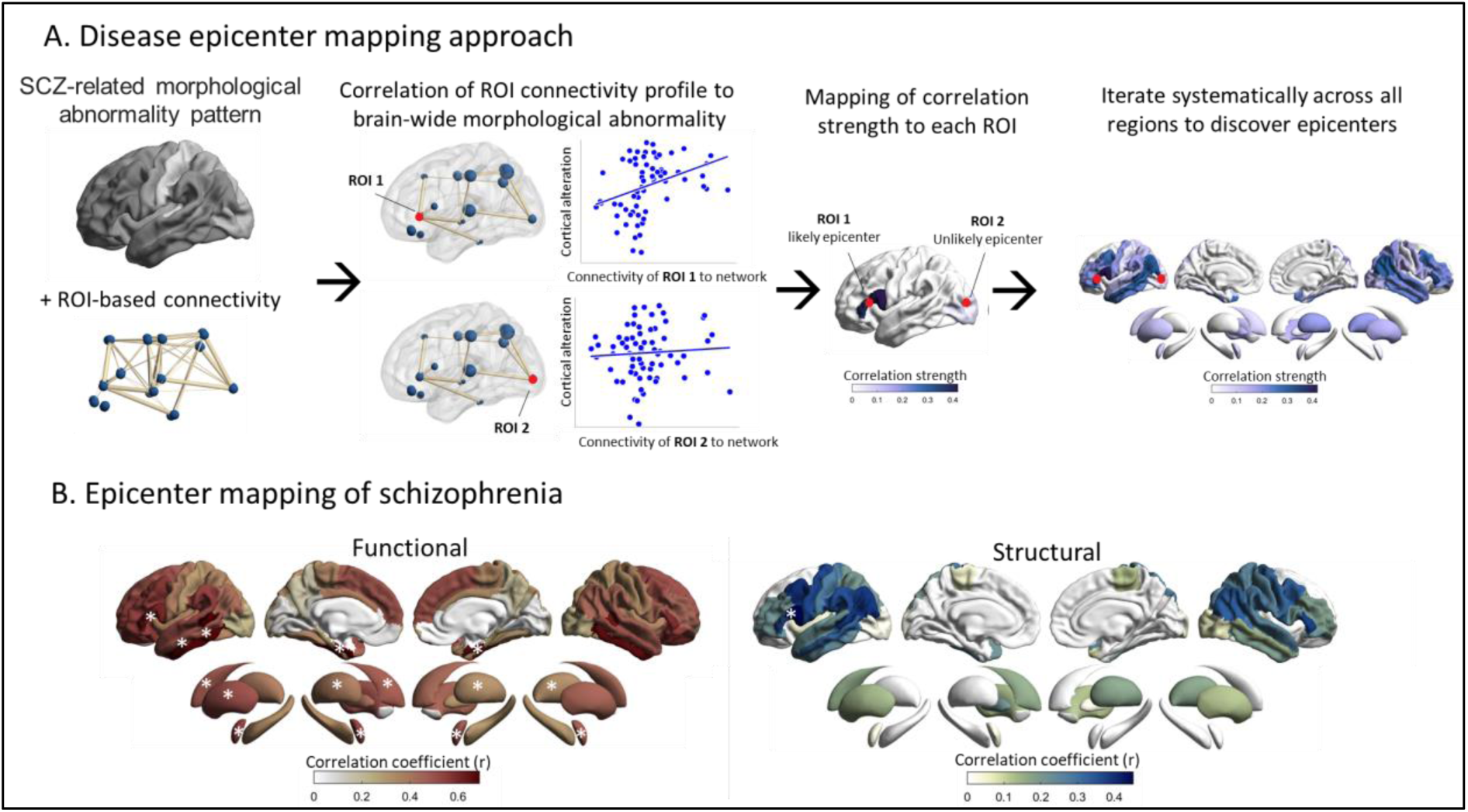
Disease epicenters. **(A)** Disease epicenter mapping schematic: Combining morphological abnormality patterns with ROI-based connectivity, we systematically computed the degree of correlation of each region to the disease-specific morphological alterations. Epicenters are regions whose connectivity profiles significantly correlated with the syndrome-specific morphological alterations; significance was assessed using spin permutation tests. This procedure was repeated systematically to assess the epicenter value of every cortical and subcortical region, as well as for both functional and structural connectivity matrices. **(B)** Correlation coefficient maps depicting the strength of association between the normative region-based functional (*left*) and structural (*right*) connectivity and the SCZ-specific morphological abnormality maps. Disease epicenters are regions more strongly connected with morphologically altered regions - and, inversely, more weakly connected with regions with less pronounced alterations. Asterisks denote the top five significant epicenters. Top-5 functional epicenters, cortical: entorhinal cortex (L+R), banks of superior temporal sulcus (L,), inferior temporal gyrus (L); subcortical: amygdala (L+R), putamen (L+R), caudate (L). Top-5 structural epicenters, cortical: pars opercularis of inferior frontal gyrus (L); subcortical: none.

### Robustness and generalizability analyses

To assess the robustness and generalizability of our mega-analytic findings, we applied a series of sensitivity analyses contextualizing the effects of different model parameters, age discrepancy, sites, disease stages, and individual clinical factors. Stability of the hub vulnerability model was tested using several other graph-based nodal metrices including betweenness, eigenvector and closeness centrality. Next, to account for age discrepancy in the ENIGMA SCZ and the HCP dataset, all analyses were repeated in age-matched subgroups of ENIGMA SCZ samples. To evaluate robustness and compare our multisite mega-analytic findings with potential site-specific effects, we repeated the hub vulnerability and disease epicenter analyses across each site. For more details, see supplementary methods and results.

#### Hub vulnerability and epicenters across different stages of SCZ

In a next step, we examined whether the associations between network features and structural alterations can be generalized or vary between different disease stages of SCZ. To this end, we investigated extreme ends of early and late disease stages that differ from the mean duration of illness (DURILL) of 11.6 (SD=7.0) years of the total sample (n=2,439) included in the main analysis. The following subgroups were defined based on available data time of scanning: first-episode psychosis (<=1 month of DURILL, n=214, mean age=25 (15–50)), early course (>1 month but <2 years DURILL, n=503, mean age=27 (15–72) and late chronic disease stage (>20 years DURILL, n=457, mean age=40 (35–77). The hub vulnerability and epicenter models were re-computed for each disease stage and compared accordingly. We investigated divergent (unique) epicenters, defined as those significant in only one of the three disease stages, and convergent (shared) epicenters, defined as those significant in all three disease stages (spin-test adjusted p-value, p_spin_ <0.05).

#### Subject-level cortical abnormality modeling

We next sought to examine whether our network-based models can be translated to individual schizophrenia patients’ data and how they are influenced by individual clinical factors. Batch-corrected cortical thickness data in patients were first adjusted for age and sex and subsequently z-scored relative to healthy controls to generate individualized cortical abnormality z-score maps. To test the hub vulnerability hypothesis, we correlated the derived patient-specific cortical abnormality z-score maps with the normative functional and structural centrality metrics (derived from HCP; Fig. 2B). The derived patient-specific correlation coefficients describe the individual scores of hub vulnerability. Subsequently, we determined each patient’s structural and functional disease epicenters by identifying brain regions whose healthy connectivity profiles were significantly correlated with the patient’s brain abnormality map. Having established both network-based models in subject-level data (see results below), we next examined the association between clinical factors and patient’s hub vulnerability and epicenters respectively. To this end, we correlated the subject-level hub vulnerability scores and the most relevant epicenters with antipsychotic medication, duration of illness and symptom severity (measured with the PANSS). For details, see supplementary methods.

### Cross-disorder comparison with ENIGMA BD and MDD

Finally, we tested whether the associations between network modeling features and brain morphometric alterations are specific for SCZ, or reflect a shared signature across other psychiatric disorders. Summary statistics from the latest cortical morphometry meta-analyses of case-control differences in BD and MDD including, in total, 4,595 patients and 12,013 controls were derived from the ENIGMA Toolbox (30). The extracted regional Cohen’s *d* values were then used to analyze the BD and MDD data using the hub vulnerability model and epicenter mapping approaches described above. To analyze the overlap between the disease-specific epicenter maps of SCZ, BD, and MDD, we compared the magnitude to which a region serves as an epicenter for a specific disease, as captured by the Pearson’s correlation coefficient *r*, to the corresponding *r* magnitude for the other two diseases. This was achieved by pairwise comparisons of the correlation coefficient of the normative connectivity profile with pairs of disorder morphological alteration patterns. The differences between correlation coefficients were compared for statistical significance with an *r*-to-*z* transform, using R package *cocor* and Zou’s CIs. Since low absolute differences would confirm the null hypothesis, we report the lowest absolute difference and corresponding CIs in the 3-way comparison (SCZ-BD-MDD).

## Results

### Hub vulnerability and epicenters of structural alteration in SCZ

Using a mega-analytic approach, we observed widespread cortico-subcortical morphological alteration patterns in people with SCZ relative to HC, closely mirroring the cortical and subcortical meta-analytic case-control maps previously reported (5, 6). Specifically, patients with SCZ showed strongest cortical thickness reduction in fronto-temporal regions including the bilateral supramarginal, temporal, and fusiform gyri, the bilateral insula, and frontal regions including the bilateral superior frontal as well as the bilateral caudal middle frontal gyri (all *t*-values > 11, FDR p< 2.2e-16, Fig. 2A, Table S5). In addition, people with SCZ showed strongest volume reductions in the bilateral hippocampus followed by bilateral thalamus, and bilateral amygdala(all *t*-values >6, FDR p<1e-11; Fig. 2A, Table S6).

#### Functional and structural degree centrality predict regional susceptibility

To test the hub vulnerability hypothesis in SCZ, we compared the spatial patterns of normative nodal degree centrality (Fig 2B) and SCZ-related morphological alterations (Fig 2A). We found that cortical thickness reductions in SCZ were more strongly pronounced in functional (*r* = 0.58, p_spin_ <0.0001), as well as structural (*r* = 0.32, p_spin_= 0.01) cortico-cortical hubs compared to non-hub regions (Fig. 1C). In other words, functional and structural degree centrality predicted susceptibility to SCZ-related cortical thickness alterations. Lower subcortical volume showed non-significant associations with functional subcortico-cortical hubs (*r* = 0.44, p_spin_= 0.11), while no relationship was observed between subcortical volume abnormalities and structural subcortico-cortical hubs (*r* = 0.03, p_spin_= 0.92; Fig. 2C). Robustness of the observed hub vulnerability in SCZ was confirmed using different centrality metrics including eigenvector, betweenness and closeness centralities (Table S10-11).

#### Disease epicenters of schizophrenia

Having identified that cortico-cortical hubs are more susceptible to cortical alterations in SCZ, we next examined whether SCZ-related cortical thickness alterations are reflected in the connectivity profile of one or more brain regions. This would indicate that these regions likely serve as network-level epicenters of the disease. We systematically compared the functional and structural connectivity profile of each cortical region to whole-brain patterns of cortical alterations in SCZ (Fig. 3A). This approach identified several temporo-paralimbic regions - including the entorhinal cortices, banks of superior temporal sulci, inferior temporal, and additionally frontal gyri (bilateral *pars triangularis*, left *pars opercularis*) - as the strongest functional and structural epicenters (all p_spin_ < 0.05; Table S7, Fig. 2B). In the subcortex, the bilateral amygdala, caudate, putamen and the left pallidum emerged as functional epicenters (all p_spin_ < 0.05; Table S8, Fig. 3B), whereby no structural subcortical epicenters were identified. Overall, highest-ranked disease epicenters were not themselves hub regions, but were rather feeder nodes, ranking on average (median) at the 40^th^ percentile of nodal centrality distribution.

### Robustness and generalizability of hub and epicenter analyses

#### Age-matched and site-specific confirmation analyses

To show the robustness of our findings to the mean age group discrepancy of the multisite ENIGMA SCZ and HC sample, and the HCP sample, we repeated the analysis with subsamples of the ENIGMA SCZ matched to the mean age of the HCP sample. Hub vulnerability and disease epicenter models revealed virtually the same results compared to the non-age-matched total ENIGMA SCZ sample (see supplementary results). These findings suggest that the network effects remain stable across age-matched and age-divergent morphological alterations patterns of the ENIGMA SCZ sample. We next examined the reproducibility of our multisite mega-analytic analyses across each participating site. Overall, the SCZ-related morphological alteration patterns were consistent across sites and showed high agreement with that obtained from the multisite mega-analysis (Fig. S1A, Table S13, supplementary results). In line with the mega-analytic findings, site-specific hub vulnerability analysis, revealed highest stability for the association between SCZ-related cortical alteration and functional cortico-cortical degree centrality (Table S14, supplementary results). As observed in the multisite findings, site-specific epicenters were most often identified in temporo-paralimbic extending to frontal brain regions (Fig S1B), with a high degree of overlap of the site-specific epicenters maps with the ones derived from the multisite aggregation (Table S15, supplementary results).

### Hub vulnerability and epicenter mapping across different disease stages

Building on the significant associations of functional and structural degree centrality and cortical alterations across the complete sample, we next asked how this association might relate to first-episode (<1 month), early (< 2 years) and late (> 20 years) stages of SCZ. We found that both functional and structural degree centrality correlated with thinner cortex in individuals with first-episode (*r*_func_ = 0.35, p_spin_ = 0.01, *r*_struc_ = 0.28, p_spin_ = 0.04), early disease stage (*r*_func_ = 0.37, p_spin_ = 0.04, *r*_struc_ = 0.24, p_spin_ = 0.09), and late chronic disease stages (*r*_func_ = 0.44, p_spin_ =0.003, *r*_struc_ = 0.28, p_spin_ = 0.04) (Fig. 4A). In other words, across all disease stages, cortical alterations were significantly greater in cortico-cortical hubs compared to non-hub regions. Epicenter mapping of cortical alterations across disease stages revealed unique occipital-parietal epicenters in first-episode psychosis and additional unique temporo-frontal epicenters in early stages, while no unique epicenters were found in chronic stages (Fig. 4B, Table S9). Across all disease stages, convergent functional and structural epicenters were circumscribed across transmodal areas including parietal, temporal and frontal regions (Fig 3B). Overall, these findings suggest localized unique epicenters for first-episode psychosis and early stages and transmodal epicenters that are shared across first-episode to chronic disease course.

**Figure 4.**
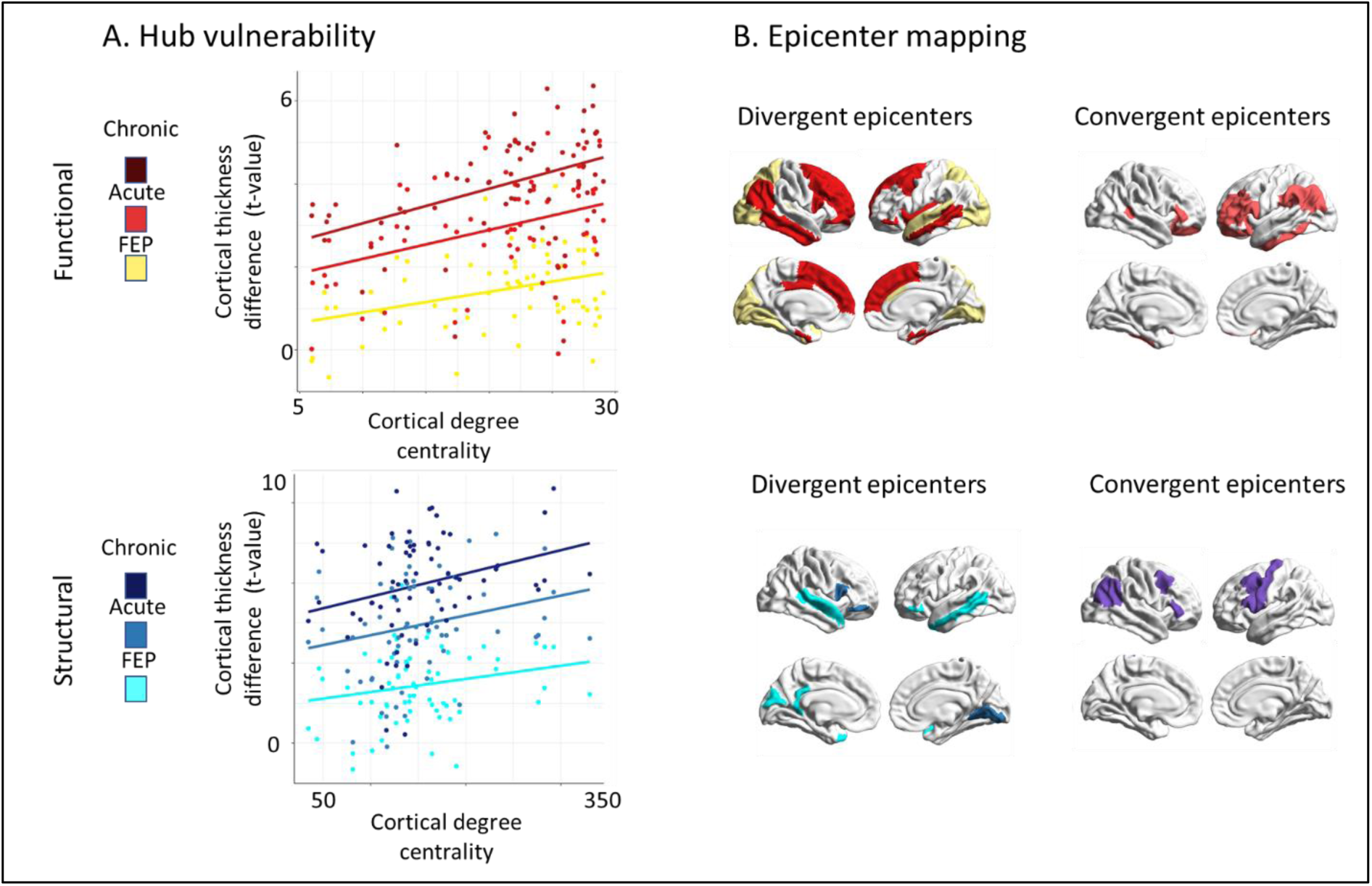
Disease stage comparison of network modeling. **(A)** Correlation of cortical thickness abnormality in SCZ to node-level functional (*upper*) and structural (*lower*) maps of degree centrality (all r>0.23, pspin<0.05). Color code indicates first-episode, early disease stage, and chronic disease stage from brighter to darker colors. **(B)** SCZ epicenter mapping (for details, see Fig. 2). Divergent (unique) epicenters are only significant in one of three disease stages with color indicating the corresponding disease stage. Convergent epicenters are significant in all three disease stages (p_spin_ <0.05). Subgroups of disease stages are defined by duration of illness: first-episode psychosis (<1 month); early disease stage (>1 month but < 2 years); chronic disease stage (>20 years). FEP: First episode psychosis.

### Subject-level cortical abnormality modeling

We next assessed whether our network-based susceptibility models also relate to brain abnormality patterns in individual patients. Although the individual subject-level data displayed overall lower sensitivity, due to the increased heterogeneity in cortical abnormality patterns, the results closely mirrored our multisite hub vulnerability and epicenter analyses (Fig. S2). We observed similar associations between patient-specific cortical abnormality maps and functional as well as structural cortico-cortical hubs (Fig. S2A). Similarly, significant epicenters were consistently observed in individual patients with temporo-paralimbic regions emerging as the most frequent epicenters (Fig. S2B). Individual level of hub vulnerability indicated by subject-level higher positive correlation coefficients was consistently associated with PANSS general symptom score (*r*_func_=0.21, p_Bonf_<0.0001, *r*_struc_=0.13, p_Bonf_=0.01, df = 383) and PANSS total score (*r*_func_=0.10, p_Bonf_=0.001, *r*_struc_=0.09, p_Bonf_=0.004, df = 1039) (Fig. 5A-B, Table S16). No significant association was observed with either positive symptoms, negative symptoms or antipsychotic medication. Correlating the individual epicenters with clinical scores we found significant positive correlations between PANSS general symptom scores and higher epicenter likelihood of sensory-motor areas extending to the cingulate gyrus and insula (all r>.18,, p_Bonf_<0.05, Fig. 5C, Table S17). For the PANSS total score, significant associations were observed for bilateral somatosensory and motor cortices extending to supramarginal gyrus (all r>.11, p_Bonf_<0.05, Fig. 5D, Table S18). None of the other epicenters were significantly associated with the clinical scores tested (Table S17-18). Robustness of these results were confirmed by running 100 permutations with 80% of the sample each time without resampling. For details, see supplementary results, Fig. S3-S8.

**Figure 5.**
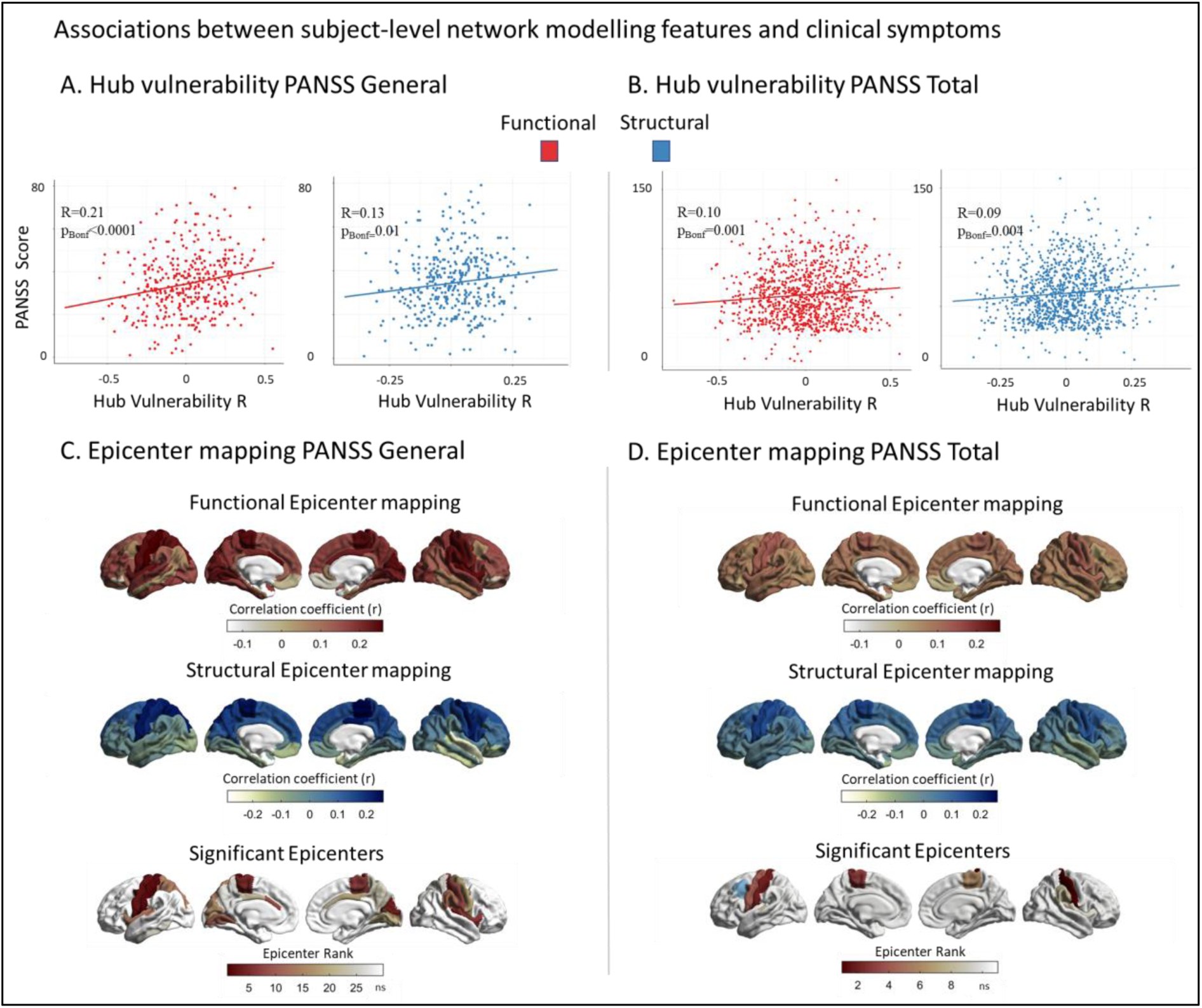
Associations between subject-level network modeling and individual symptoms. **(A)** Correlation between subject-level functional (red) and structural (blue) hub vulnerability of cortical alterations and PANSS general scores (r_func_=0.21, p_Bonf_<0.0001, r_struc_=0.13, p_Bonf_=0.01). **(B)** Correlation between subject-level functional (red) and structural (blue) hub vulnerability of cortical alterations and PANSS total scores (r_func_=0.10, p_Bonf_=0.001, r_struc_=0.09, p_Bonf_=0.004). **(C)** Correlations between subject-level functional (red) and structural (blue) epicenter likelihood and PANSS general scores. Significant associations were only found for functional epicenters spanning from sensory-motor areas to cingulate gyrus and insula (all r>.18, p_Bonf_<0.05). **(D)** Correlations between subject-level functional (red) and structural (blue) epicenter likelihood and PANSS total scores. Significant associations were found for functional epicenters in the bilateral somatosensory and motor cortices extending to supramarginal gyrus (in blue) as the only structural epicenter ((all r>.11, p_Bonf_<0.05).

### Cross-disorder comparison of hub vulnerability and epicenter mapping

Last, we wished to explore whether our network models are specific to SCZ or rather reflect transdiagnostic features along the schizophrenia-affective disorder spectrum, we applied both network-based susceptibility models to meta-analytic cortical maps of BD and MDD. The hub vulnerability model revealed that cortical thickness alterations in BD were correlated with cortico-cortical hubs (*r*_struc_ = 0.35, p_spin_ = 0.01; *r*_func_ = 0.32, p_spin_ = 0.07) as well as marginally with functional subcortico-cortical hubs (*r*_func_ = 0.52, p_spin_ = 0.06). No associations were found in MDD (all p_spin_ > 0.05, Fig. 6A). In BD, epicenter modeling identified functional and structural disease epicenters predominantly in the frontal cortices specifically, the bilateral orbitofrontal cortex, the middle frontal gyrus, and the left inferior frontal gyrus (all p_spin_ < 0.05, Fig. 6B). By contrast, no significant cortical epicenters were observed in MDD (all p_spin_ >0.05, Fig. 6B). In the subcortex, the caudate nucleus emerged as the only functional and structural epicenter in BD, and the nucleus accumbens as sole structural epicenter in MDD (all p_spin_ <0.05, Fig. 6b). Correlating the epicenter maps of SCZ and BD revealed a significant overlap both cortically (r_func_ = 0.88, p_spin_ < 0.05, *r*_struc_= 0.74, p_spin_ < 0.05) and to a lesser extent subcortically (*r*_func_ = 0.63, p= 0.007, *r*_struc_= 0.41, p= 0.07, Fig. 6C). In the cerebral cortex the following regions emerged as unique epicenters for SCZ; banks of the superior temporal sulcus (diff_R_= 0.29, CI_R_: 0.15 – 0.47; diff_L_= 0.28, CI_L_: 0.16 – 0.45), the superior temporal gyrus (diff_R_= 0.33, CI_R_: 0.18 – 0.49; diff_L_= 0.33, CI_L_: 0.19 – 0.49), the supramarginal gyrus (diff_R_= 0.24, CI_R_: 0.09 – 0.40; diff_L_= 0.21, CI_L_: 0.07 – 0.37), the right entorhinal cortex (diff = 0.24, CI: 0.11 – 0.40), the right pars opercularis of inferior frontal gyrus (diff = 0.16, CI: 0.01 – 0.31). In the subcortex the bilateral amygdala (diff_R_= 0.35, CI_R_: 0.20 – 0.50; diff_L_= 0.29, CI_L_: 0.20 – 0.50), and putamen (diff_R_= 0.16, CIR: 0.01 – 0.32; diff_L_= 0.14, CIL: 0.01 – 0.30), emerged as unique epicenters (Fig. 6C). In BD the left caudate nucleus (diff = 0.25, CI: 0.10, 0.40) and in MDD the nucleus accumbens bilaterally (diff_R_= 0.45, CI_R_: 0.20 – 0.68; diff_L_= 0.35, CI_L_: 0.10 – 0.58), were found to be disease-specific epicenters (Fig. 6C). Taken together, these findings indicate some convergence of the hub vulnerability and epicenter models in SCZ and BD, but not in MDD. For SCZ, specific temporo-limbic epicenters were identified, while distinct regions within the striatum emerged as unique epicenters for SCZ, BD, and MDD.

**Figure 6.**
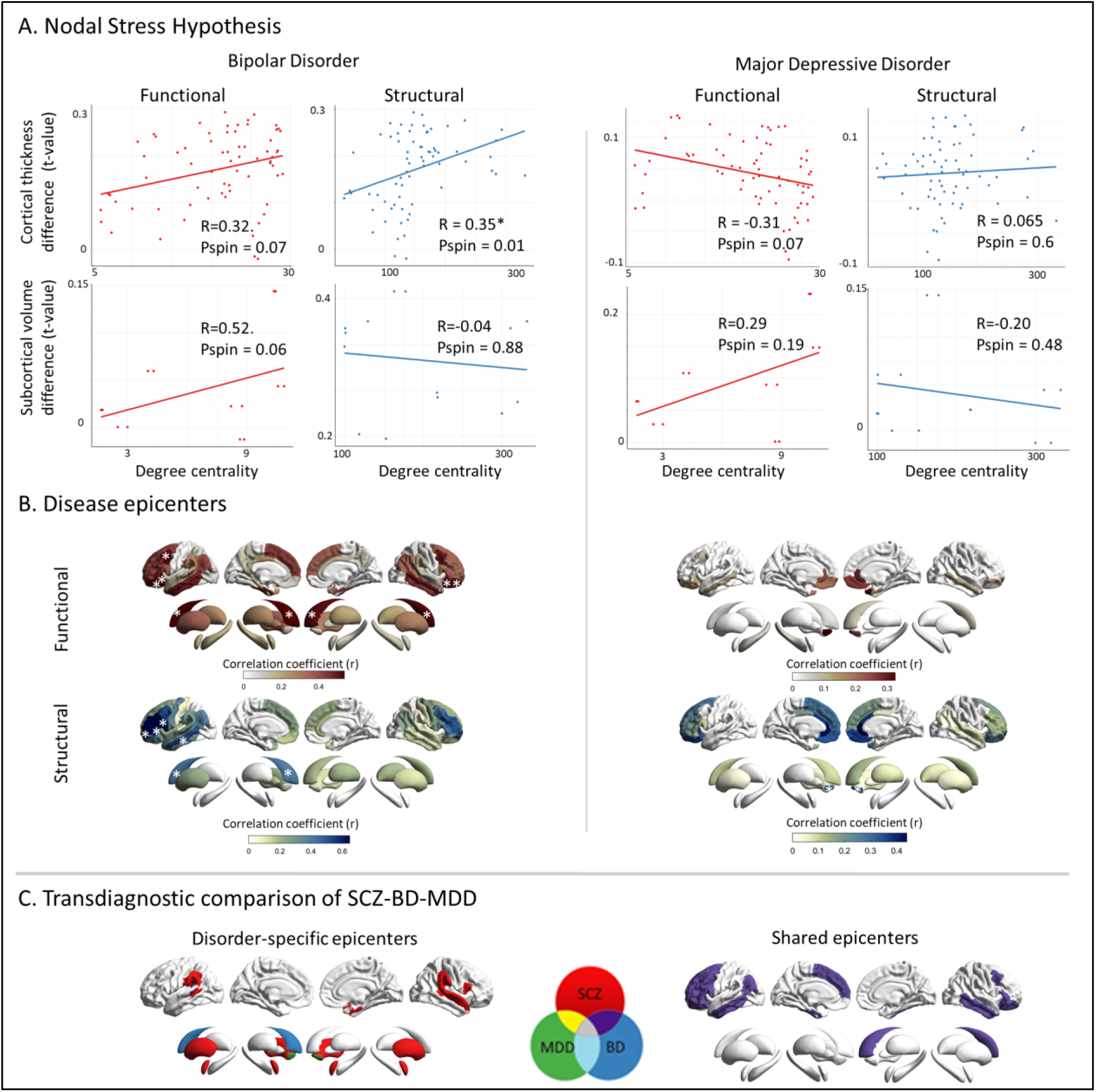
Cross-disorder comparison of network modeling. (**A**) Correlation of disorder-related gray matter alterations to node-level functional (left) and structural (right) maps of degree centrality in BD and MDD. In BD, cortical regions with high structural centrality are significantly more likely to display higher morphological alterations; this trend is also observed for functional centrality. No such relationship is observed in MDD. (**B**) Correlation coefficient maps depicting strength of association between the normative region-based functional (top) and structural (bottom) connectivity and the BD-(left) and MDD-specific (right) morphological alteration maps. Asterisks denote the significant epicenters. Functional BD epicenters: pars orbitalis of inferior frontal gyrus (L), lateral orbitofrontal cortex (L), caudal middle frontal gyrus (L), inferior and middle temporal gyrus (L) cortically and the caudate nucleus (L) in the subcortex (func: bilateral, struc: left). Structural BD epicenters: pars opercularis (L) and pars triangularis (L) of inferior frontal gyrus, rostral middle frontal gyrus (L), middle temporal gyrus (L) cortically and the caudate nucleus (L), subcortically. In MDD, no functional epicenters were detected. Structural MDD epicenters: nucleus accumbens (R). (**C**) Transdiagnostic comparison of epicenters between SCZ, BD, and MDD. Disorder-specific epicenters which are only significant in one disorder are displayed on the left side. Shared epicenters between at least two disorders are shown on the right side. Please note, that shared epicenters were only found for SCZ and BD but not MDD: Bipolar disorder; MDD: Major depressive disorder; SCZ: Schizophrenia

## Discussion

Our study *(i)* explored spatial associations between normative brain network architecture with morphometric alterations of SCZ and *(ii)* systematically assessed the robustness of these associations across different sites and inter-individual factors, leveraging the largest multisite curated neuroimaging dataset to date. We found that cortical regions with higher cortical network centrality were, on average, more vulnerable to SCZ-related cortical alterations in all stages of the disease, while subcortical regions with high subcortico-cortical centrality did not share this property. Examining the degree to which each brain region’s network connections match the morphological alteration patterns of SCZ, we identified temporo-paralimbic regions extending to the inferior frontal gyrus as the most probable disease epicenters. Robustness and generalizability of findings was shown using different centrality measures, ages, individual sites and subject-level data. Across disease stages, we observed unique epicenters for first-episode and early stages (<2years), and shared transmodal epicenters across first-episode to chronic (>20years) stages. Translating our network models to subject-level data, we found that individual hub vulnerability and sensory-motor as well as paralimbic epicenters were associated with overall symptom severity. Finally, *(iii)* our cross-disorder comparison identified similar associations of cortical node centrality and cortical alterations in BD and SCZ, but not in MDD and SCZ. Additionally, we observed shared epicenters in SCZ and BD as well as unique epicenters for SCZ, BD, and MDD. Overall, our work contributes to recognizing potential network mechanisms that underlie macroscale structural alterations and inter-individual variability in SCZ.

By combining our findings of SCZ-related cortical thickness alterations with normative brain connectivity information, we provide evidence for the hub vulnerability hypothesis in SCZ, i.e., the centrality of a cortical region in terms of its position in both the functional and the structural connectome is well correlated with the degree of its deformation. Wannan *et al.* (15) previously found that cortical deformation tends to cluster in regions with high normative structural covariance, and Shafiei *et al.* (14) have shown that the deformation of a cortical region correlates with the collective deformation of its network-neighboring regions, weighted by their respective connection strength. Prior studies further observed that network-central cortical nodes (54, 55) and the neuroanatomical tracts connecting them to other regions (56) are disproportionately affected in SCZ. Our findings extend these observations by identifying robust network organizations of SCZ-related alterations leveraging multiple level of investigation including large-scale multisite, site-specific and subject-level analyses. We found that relationships between regional cortical alterations and regional node centrality were present in first-episode, early, and late disease stages. Individual hub vulnerability further correlated with clinical profiles of higher general symptom severity. These findings suggest that network organization may be guiding cortical alterations throughout the disease process of SCZ. The vulnerability of cortical hub regions to deficiencies in cortical thickness was originally examined in neurological disorders, including Parkinson’s disease (23), Alzheimer’s disease (20, 24), and other neurodegenerative conditions (25), as well as epilepsy (26). Several biological mechanisms could lead to a marked susceptibility of hub regions. According to the stress-threshold model, centrally located neuronal populations are especially vulnerable to pathophysiological perturbations due to their higher neuronal activity and concomitant metabolic demands (55, 57). In addition, hub regions are more likely to be part of the association cortices which, partly due to their prolonged window of developmental plasticity, have been shown to be especially vulnerable to developmental insults (58) - tying into the known neurodevelopmental aspects of SCZ pathogenesis (59).

Having identified the relevance of brain connectivity as a factor potentially contributing to morphological alteration in SCZ, we tested if the underlying connectivity of specific regions – or epicenters – constrains the SCZ-related cortical abnormality pattern. We identified temporo-paralimbic brain regions extending to the frontal gyrus as the most probable SCZ epicenters. Epicenters were not restricted to cortical regions with caudate/putamen and amygdala suggesting that cortical alterations might also be guided by subcortico-cortical connectivity. Across all regions, the bilateral entorhinal cortex was most consistently the top-ranked epicenter for SCZ. This is in line with recent reports in first-episode psychosis (60) and chronic SCZ (19), while also mirroring epicenters of cortical atrophy in neurodegenerative disorders (61). Our epicenter mapping across disease stages revealed unique occipital-parietal epicenters in first-episode psychosis and shared transmodal areas across disease course, confirming recent observations (60). In addition, higher general symptom severity was associated with sensory-motor and paralimbic (cingulate, insula) epicenters suggesting that state-dependent clinical facets contribute to the development of individual epicenter profiles. The consistent involvement of temporo-paralimbic and frontal cortical regions, as potential epicenters across all disease stages, aligns with a series of studies implicating these regions in SCZ pathophysiology. Cortical thickness deficits in these regions emerge early in the disease (13, 62), are predictors for conversion (62) and are most strongly affected in chronic SCZ (5). More generally, aggregation of epicenters in the temporo-paralimbic cortex could be explained by regional cellular and molecular features that promote brain plasticity, higher metabolic activity, making the paralimbic cortex more vulnerable to developmental disruptions (63, 64). In terms of their positioning in the network, the most significant epicenters were not themselves hubs, but rather regions with centrality slightly above the median. If they are indeed active in the pathophysiological process, epicenters could represent regions affected earlier in the disease process, and serve as a gateway through which hub nodes are affected downstream due to their connections to these nodes.

Our cross-disorder comparison showed that the hub vulnerability models were also implicated in cortical alterations in BD, while cortical findings in MDD did not share this property. Epicenter mapping revealed some convergence of SCZ and BD along a temporo-limbic to frontal gradient, with an accentuation of unique epicenters in temporo-paralimbic regions in SCZ. In the subcortex disease-specific epicenters were the nucleus accumbens for MDD, the left caudate for BD, and bilateral putamen and amygdala for SCZ, while right caudate reflected a shared epicenter of SCZ and BD. This suggest that subcortico-cortical connectivity contribute to disease-specific cortical alterations that partly differentiates affective from psychotic processes along a ventral to dorsal striatal axis. Overall, the magnitude of observed cortical and subcortical disease epicenters was most pronounced in SCZ, intermediate in BD and relatively lacking in MDD, supporting evidence of overlapping neurobiological substrates across the SCZ-BD spectrum. Both, the hub susceptibility and epicenter models further revealed differential aspects of functional and structural connectivity to SCZ and BD-specific pathophysiology. Specifically, hubs and epicenters of SCZ and BD were more sensitive to detection with the functional and structural connectome, respectively. This finding could reflect the inherent differential sensitivity of diffusion MRI to long-range fiber bundles and that of resting-state functional MRI to short-range intracortical spatially distributed polysynaptic cortical systems (65). Disease epicenters in SCZ preferentially occurred in polysynaptic temporo-limbic regions, making them more amenable to detection through the functionally-derived connectome. Conversely, disease epicenters in BD appeared more stable across modalities, with an accentuation in frontal regions, thus lending structural connectivity an increased sensitivity to their detection. Collectively, our findings suggest that shared network architecture features across SCZ and BD reflect the high biological proximity of these two disorders (5–8, 34, 40), while the unique epicenters could reflect biological differences driving their differential clinical manifestations. By contrast, cortical abnormalities in MDD seem to lack a straightforward network influence - consistent with its more different structural brain signature compared to SCZ. However, given that we used meta-analytic data, the lack of findings in MDD could reflect heterogeneity in the study samples included, whereby specific cohorts with psychotic depression could potentially display more similarities to SCZ and BD.

Several implications could arise from the associations between hub vulnerability, epicenter mapping, and morphometric signatures of SCZ. According to our network models, epicenters emerging early in the disease process might be further characterized by distinct alterations in genetic activity, macro- or micromorphology and/or activity. Aberrant signaling of cortical development involving myelination, branching, and synaptic pruning (34, 66, 67) might propagate from focal epicenters to connected regions in a network-spreading pattern leading to the characteristic pattern of cortical alteration. Applying multi-omic approaches longitudinally across neurodevelopment and early SCZ stages are of interest to test this hypothesis and examine the role of the derived epicenters in aberrant developmental processes. In SCZ such network-spreading phenomenon might not be limited to neurodevelopmental processes but additionally involves neurodegeneration later in life. Trans-neuronal spread of misfolded proteins reflecting brain network organization have been documented in several neurodegenerative disorders (22, 23, 68, 69). In this regard, hub regions would be expected to be preferentially affected during the disease process of SCZ. Insights from neurological disorders with a similar pattern could be informative, in terms of recognizing potentially common upstream pathways to better understand potential neurodegenerative processes of cortical thickness deficiencies in SCZ.

The present study is limited by its cross-sectional nature and cannot provide definitive answers pertaining to the temporal unfolding of the pathophysiological processes. With the increasing availability of longitudinal studies, future studies could more mechanistically test a “spreading” model, whereby the pathophysiology could originate from disease-specific epicenter regions, and then, through these, propagate to the vulnerable hub regions. Finally, although HCP is a current benchmark dataset to obtain normative structural and functional brain connectivity information, the use of individualized structural and functional connectivity information in patients might further improve the accuracy of network modeling of schizophrenia.

## Conclusion

Leveraging a large-scale, multisite, neuroimaging dataset from the ENIGMA-Schizophrenia working group, this study provides robust evidence that brain network architecture is closely and robustly associated to cortical alterations in SCZ. Nodal centrality and temporo-paralimbic to frontal epicenters emerge as fundamental components of this relationship. We further demonstrated that network organization may guide cortical alteration patterns throughout the disease stages of SCZ and contribute to individual patient-specific alteration profiles. SCZ showed some convergence in hub vulnerability and epicenters with BD, suggesting a partial overlap in network mechanism. Yet, the unique epicenters observed in SCZ indicate additional network-spreading processes specific to SCZ. Overall, our work contributes to recognizing potentially common pathways to better understand macroscale structural alterations, and inter-individual variability in SCZ.

## Acknowledgments

See Supplement

