## Supplement for "Connectome architecture shapes large-scale cortical alterations in schizophrenia: a worldwide ENIGMA study"

Georgiadis et al.

### Supplementary Methods and Results

#### Contents


- Table S1. Site-specific inclusion criteria
- Table S2. Site demographics
- Table S3. Sample image acquisition and image pre-processing details by cohort
- HCP MRI data and preprocessing
- Functional and structural connectivity matrix generation from HCP participants
- ComBat batch effect adjustment
- Table S4 ComBat batch effect adjustment confirmation
- Table S5. Cortical thickness differences between schizophrenia patients and healthy controls
- Table S6. Subcortical volume differences between schizophrenia patients and healthy controls
- Table S7. Schizophrenia cortical epicenter ranking
- Table S8. Schizophrenia subcortical epicenter ranking
- Tables S9. Divergent and convergent regions of different stages of schizophrenia
- Mega-analysis robustness and sensitivity analyses
  – Reproducibility across different centrality metrics
  – Table S10. Correlation matrix of centrality measures
  – Table S11. Hub vulnerability across centrality metrics
  – Reproducibility across HCP age-matched and age-divergent ENIGMA schizophrenia samples
  – Robustness and site-specific confirmation analysis of hub vulnerability and disease epicenter models in schizophrenia
  – Table S13. Correlation between site-specific and mega-analytic cortical alteration maps of schizophrenia
  – Figure S1. Site-specific epicenter mapping
  – Table S14. Site-specific hub vulnerability analysis
  – Table S15. Site-specific epicenter map agreement with mega-analytical epicenter map
- Subject-level cortical abnormality modeling
- Figure S2. Individual-level network modeling analysis in schizophrenia
- Subject-level correlation of clinical variables to hub vulnerability and epicenters

- Table S16. Correlations between subject-level functional and structural hub vulnerability and clinical scores
- Table S17. Correlations between individual subject-level functional epicenters and clinical scores
- Table S18. Correlations between individual subject-level structural epicenters and clinical scores
- Subject-level robustness and sensitivity analysis
– Figure S3. Stability of correlation of hub vulnerability at the individual level to all clinical variables
– Figure S4 Stability of correlation of epicenter values at the individual level to PANSS positive symptoms
– Figure S5. Stability of correlation of epicenter values at the individual level to PANSS negative symptoms
– Figure S6. Stability of correlation of epicenter values at the individual level to PANSS general symptoms
– Figure S7. Stability of correlation of epicenter values at the individual level to PANSS total score
– Figure S8. Stability of correlation of epicenter values at the individual level to chlorpromazine analogues
– Figure S9. Stability of correlation of epicenter values at the individual level to duration of illness
- Acknowledgements per dataset
- References

#### Table S1. Site-specific inclusion criteria

| Cohort | Country | Diagnosis measurement | Sample characteristics/inclusion criteria | Exclusion criteria |
|---|---|---|---|---|
| **ASRB** | Australia | Diagnosis was confirmed using the OPCRIT algorithm applied to interviewer ratings on the DIP, acc. to ICD-10 criteria | All participants were fluent English speakers and aged 18-65 years old | No history of an organic brain disorder, brain injury accompanied by > 24 h of amnesia, mental retardation defined as an IQ < 70, movement disorder, current substance dependence, or electro-convulsive therapy in the preceding 6 months. The control participants additionally had no personal history of psychotic disorder or family history of psychotic disorder in their first-degree biological relatives. |
| **CAMH** | Canada | SCID DSM-IV-TR Axis I | Schizophrenia outpatients who were clinically stable as determined by no medication change within the past month | Exclusion criteria were intelligence quotient < 70 as estimated by the Wechsler Test for Adult Reading (WTAR), substance dependence or abuse reported or indicated by a urine toxicology screen, head trauma with loss of consciousness, neurological disorders, and any magnetic resonance imaging contraindications. A first degree relative with a primary psychotic disorder was also an exclusion criterion for controls. |
| **CIAM** | South Africa | SCID DSM-IV-TR Axis I | Stable outpatients between ages 19-40 years with a diagnosis of schizophrenia or bipolar type I disorder with psychotic features or methamphetamine induced psychotic disorder or controls without a history or family history of psychotic symptoms. | Patients were excluded if they had a psychotic disorder other than schizophrenia or bipolar type I with psychotic features or methamphetamine induced psychotic disorder (i.e., schizophreniform disorder). Patients or controls were excluded if they had a physical condition requiring medication, prior head trauma or neurosurgery, any history of a cardiovascular event, a history or family history of epilepsy, a learning disability, if they were pregnant or lactating, or had any metal brain implants. Patients with bipolar type II disorder were excluded. Patients in the methamphetamine psychosis group were excluded if there was any evidence of symptoms persisting longer than 1 month after the cessation of methamphetamine use or if there was evidence of prior psychotic symptoms not related to the use of methamphetamine. |
| **COBRE** | USA | SCID DSM-IV Axis I Disorders | All participants were in the 18-65 age range and had a diagnosis of schizophrenia. Healthy individuals were included if they did not have a personal or family history of psychiatric disorders. | History of neurological disorder, history of mental retardation, history of severe head trauma with more than 5 minutes loss of consciousness, history of substance abuse or dependence within the last 12 months and MRI contraindications. |
| **ESO** | Czech Republic | ICD-10 (F20.x, F23, F25) | Early or first-episode psychosis, Czech language as a mother tongue, 18-60 years old | Neurocognitive disorders (organic mental disorder), mental disorders caused by addiction, mental retardation (IQ<80), severe neurological disorder, head injury, hypertension, cerebrovascular disease, epilepsy, migraine, endocrine disorders. |
| **FOR210 Marburg** | Germany | Semi-structured interview using SCID DSM-IV-TR Axis-I Disorders | All participants were aged between 18-65 and were fluent German speakers. Patients ( in-and out-patients) had a | Exclusion criteria were any history of neurological (head trauma or unconsciousness) andmedical condition (severe somatic disorders), magnetic resonance imaging |

| | | | | |
|---|---|---|---|---|
| | | | lifetime diagnosis of schizophrenia. Healthy individuals were included if they did not have any lifetime history of psychiatric disorders. | contraindications, verbal IQ < 80 (assessed using MWT-B), current substance dependence or benzodiazepine treatment. |
| **FOR210 Muenster** | Germany | Semi-structured interview using SCID DSM-IV-TR Axis-I Disorders | All participants were aged between 18- 65 and were fluent German speakers. Patients (in-and out-patients) had a lifetime diagnosis of schizophrenia. Healthy individuals were included if they did not have any lifetime history of psychiatric disorders. | Exclusion criteria were any history of neurological (head trauma or unconsciousness) and medical condition (severe somatic disorders), magnetic resonance imaging contraindications, verbal IQ < 80 (assessed using MWT-B), current substance dependence or benzodiazepine treatment. |
| **FIDMAG** | Spain | DSM-IV criteria based on interview and review of clinical history | Patients had a diagnosis of schizophrenia. All participants were in the 18-65 age range. | Controls were excluded if they reported a history of mental illness and/or treatment with psychotropic medication. Patients were excluded if have had a history of brain trauma or neurological disease or had shown alcohol/ substance abuse within 12 months before participation |
| **FSLRome** | Italy | SCID DSM-IV Axis I Disorders (SCID-I) andSCID DSM-IV Axis II Personality Disorders (SCID-II) | Inclusion criteria were (i) age between 18 and 65years; (ii) at least five years of education; and (iii) suitability for MRI scanning. | Exclusion criteria were (i) history of alcohol or drug abuse in the two years before the assessment; (ii) lifetime drug dependence; (iii) traumatic head injury with loss of consciousness; (iv) past or present major medical illness or neurological disorders; (v) any (for HC) or additional (for patients) psychiatric disorder or mental retardation; (vi) dementia or cognitive deterioration according to DSM-IV-TR criteria, and Mini-Mental State Examination (MMSE) score<25, consistent with normative data in the Italian population; (vii) not able and willing to give written informed consent. |
| **GIPSI** | Colombia | DSM-IV-TR diagnosis criteria using the Diagnostic Interview for Genetic Studies (DIGS) | Subjects with diagnosis of Schizophrenia, between the ages of18 and 60 years old. | History of traumatic brain injury, personality disorders or autism spectrum disorders. |
| **IGP** | Australia | Diagnosis was confirmed using the OPCRIT algorithm applied to interviewer ratings on the DIP, acc. to ICD-10 criteria | All participants were fluent English speakers and aged 18-65 years old | General exclusion criteria included an inability to communicate sufficiently in English, a current neurological disorder, a diagnosis of substance abuse or dependence in the pastsix months; and/or having been treated with electroconvulsive therapy in the previous six months. |
| **MCIC** | USA | SCID DSM-IV (SCID-NP for controls) or CASH were used to diagnose primary and co-morbid psychiatricdisorders in controls and patients | All subjects were between the ages of 18 and 60 and spoke English as their native language. To be included in the schizophrenia cohort, patients had to meet diagnostic criteria for schizophrenia, schizoaffective disorder, or schizophreniform disorder. Concerted effort was made to recruit | Control subjects who met criteria for current or past history of substance abuse or dependence were excluded from the study. Patients, however, were not excluded from the study unless criteria were met for current (i.e., within the past month) abuse or dependence (except for 6 patients who were found to meet criteria for current abuse after the study data was collected). Both patients and controls were excluded if they had (1) an IQ less than 70 based on a standardized IQ test, (2) history of a head injury resulting in prolonged loss of consciousness, neurosurgical procedure, neurological disease, history of skull fracture, severe or disabling medical conditions, or (3) a contraindication for MRI scanning such as |

| | | | | |
|---|---|---|---|---|
| | | | patients early in the course of their illness and especially those who were antipsychotic drug naïve. The healthy control subjects with no current or past history of psychiatric illness including substance abuse or dependence were matched within site to the patient cohort for age, sex, and parental education. Control subjects who had not been diagnosed with any psychiatric disorders, but had been medicated with antidepressants, anti-anxiety medication or medication for sleep disturbance were included in the study provided that the duration of their medication did not exceed 2 months of lifetime use and no medication was used within the 6 months preceding the baseline MRI scan. | pregnancy, metal in body or head including implanted pacemaker, medication pump, vagal stimulator, deep brain stimulator, implanted TENS unit, or ventriculo -peritoneal shunt |
| **MPRC** | USA | SCID DSM-IV combined with a review of medical records | Individuals diagnosed with schizophrenia or schizoaffective disorder; and healthy controls without current DSM-IV Axis I psychiatric illnesses. | The exclusion criteria included diagnosis with hypertension, hyperlipidemia, type 2 diabetes, heart disorders, major neurologic event such as stroke or transient ischemic attack, and recent substance use disorder (except tobacco and marijuana use). |
| **OLIN** | USA | SCID DSM-IV | **AA:** Participants (ages 18-70) were healthy controls and individuals with a DSM-IV diagnosis of schizophrenia or schizoaffective disorder. Healthy controls were allowed to have common psychiatric disorders (exceptfor any type of psychosis).<br>**BSNIP:** Participants (ages 15-65) were recruited from 5 sites (Hartford, Baltimore, Chicago, Dallas, Boston) and included healthy controls, individuals with a DSM-IV diagnosisof schizophrenia, schizoaffective disorder. Healthy controls had no | **AA:** Exclusion criteria for all subjects included a history of major medical disorders, severe head injury, MRI contraindication, IQ < 70, dementia, traces of drugs (excluding THC) in urine, or drug intoxication during cognitive or MRI assessment.<br>**BSNIP:** Exclusion criteria for all subjects included history of seizures or head injury with loss of consciousness >10 minutes; positive urine drug screen for common drugs of abuse on the day of testing; diagnosis of substance abuse in the past 30 days or substance dependence in the past 6 months; history of systemic medical or neurological disorder likely to affect cognitive abilities; age-corrected Wide-Range Achievement Test, 4th edition, reading test standard score <65; and < 6th grade English reading level.<br>**BPP:** Exclusion criteria for all subjects included alcohol or drug abuse or dependence within the past 6 months, a history of major medical or neurological disorders, or IQ <70 as assessed by the WAIS. |

| | | | | |
|---|---|---|---|---|
| | | | personal or family history (first degree) of psychotic disorders; no personal history of recurrent mood disorder; no lifetime history of substance dependence; and no history of any significant cluster A Axis II personality features defined bymeeting full criteria or within 1 criterion of a cluster A diagnosis usingthe Structured Interview for DSM-IV Personality. We provided healthy control and SZ data only from theHartford site.<br>**BPP:** Participants (ages 18-70) were healthy controls. Healthy control subjects were included if they had nolifetime history of axis I psychiatric disorders as assessed by the SCIDand no family history of mood or psychotic disorders. | |
| **PAFIP** | Spain | SCID DSM-IV for patients confirmed by an independent psychiatrist 6months after the initial contact. CASH for controls. | Patients had to meet the following criteria: (1) age 15–60 years; (2) living in the catchment area; (3) experiencing a first episode of psychosis; (4) no prior treatment with antipsychotic medication or, if previously treated, a total lifetime of adequate antipsychotic treatment of less than 6 weeks; and (5) meeting DSM-IV criteria for schizophrenia, schizophreniform disorder, briefpsychotic disorder, or schizoaffective disorder. | Patients were excluded when meeting DSM-IV criteria for (1) drug dependence (except nicotine dependence), (2) mental retardation, and when having a history of neurological disease or head injury. Controls exclusion criteria were current or past history of psychiatric, neurological or general medical illnesses, including substance dependence and significant loss of consciousness. HCs were selected to have a similar distribution in age, sex, laterality index, drug history and years of education as the patient population. The absence of psychosis in first-degree relatives was also confirmed by clinical records and family interview. After a detailed description of the study, each subject gave written informed consent to participate. |
| **PENS** | USA | SCID for DSM-IV and DIGS Psychosis module performed by a trained clinical interviewer. Then all clinical materials | The sample included individuals with a schizophrenia spectrum disorder (n = 35) or bipolar disorder, first degree biological relatives of persons with a schizophrenia spectrum or bipolar | Participants were native English speakers, 18 to 60 years old, with normal or corrected hearing and vision, and IQ of at least 70. Participantswith a history of intellectual disability were excluded. Patients and controls were additionally excluded for substance abuse or dependence. within the past 6 months; history of electroconvulsive therapy, epilepsy, |

| | | | | |
|---|---|---|---|---|
| | | reviewed by doctoral and graduate level psychologists to achieve a consensus diagnosis. | disorder, and healthy controls. Participants were recruited from the Minneapolis Veterans Affairs Health Care System (VAHCS) and mental health centers in the Minneapolis community as part of a larger research protocol that included neurocognitive, MRI, and additional electroencephalography procedures. | diagnosed seizure disorder, stroke, or neurological condition; uncontrolled medical condition likely to substantially affect brain functioning (e.g., untreated thyroid condition); and head injury resulting in fractured skull or more than 30 minutes unconsciousness. Healthy controlswere also excluded for history of primary psychotic disorder or hypomania, antipsychotic medication use, current or past depressive episodes, attention-deficit/hyperactivity disorder or other learning disability, and family history of bipolar or psychotic disorder. |
| **PHCP** | USA | SCID for DSM-IV and DIGS Psychosis module performed by a trained clinical interviewer. Then all clinical materials reviewed by doctoral and graduate level psychologists to achieve a consensus diagnosis. | People with Psychosis (PwP) were between the ages of 18 and 65 years old with a diagnosis of schizophrenia, schizoaffective disorder, or bipolar I disorder with a history of psychotic symptomatology (i.e., delusions or hallucinations) with no indication that symptoms were caused by substance use or a general medical condition. While PwP were screened and excluded for current substance use issues, a history of such issues as well as current/lifetime comorbidities of any kind were permitted for enrollment in the study in order to have a sample representative of patients with psychosis in the general population while simultaneously limiting nuisance. effects. | To be eligible for enrollment, all participants spoke English as their primary language and did not have: a legal guardian (or otherwise lack capacity to provide informed consent), alcohol/drug abuse in the past month or alcohol/drug dependence in the last 6 months, a diagnosed Learning Disability or estimated IQ lower than 70 (if either condition was diagnosed based on testing by a trained professional or the latter by research staff), a current or past central nervous system disease (including: seizures, epilepsy, encephalitis, MS, Parkinson's, stroke), his- tory of head injury with skull fracture or loss of consciousness greater than 30 min, history of electro-convulsive therapy (ECT) in the last year, tardive dyskinesia (as evidenced by medical record), obstructed or com- promised vision (e.g., lazy eye that is uncorrected or was corrected after age 17 / strabismus / cross eyes / permanent eye injury / abnormality in visual field / cataract), hearing problems (e.g., cannot hear with- out hearing aid / severe tinnitus), or a condition likely making it impossibleto perform tasks (e.g., paralysis, severe arthritis). |
| **RSCZ** | Russian Federation | ICD-10 (F20.x) | Early or first-episode psychosis in-patients (no later than 5 years since the first episode) who were clinically stable and received antipsychotic medication therapy. Mentally healthy controls were recruited from acquaintances of the researchers and clinical staff. All participants were fluent Russian speakers, right-handed males. | Common exclusion criteria for patients and controls were: organic brain disorders, neurological or severe somatic disorders, mental retardation, alcohol or substance abuse, history of head trauma with loss of consciousness for more than 5 min. In addition, controls were excluded if they had a family history of psychiatric illness. |

| | | | | |
|---|---|---|---|---|
| **SCORE** | Switzerland | ICD-10 or DSM-IV criteria | First-episode psychosis patients who fulfilled criteria for brief psychotic disorder. All patients were between 18 and 42 years of age. | History of previous psychotic disorder, psychotic symptoms secondary to an organic disorder, substance abuse (except nicotine), psychotic symptoms associated with an affective psychosis or a borderline personality disorder, age younger than 18 years, inadequate knowledge of the German language, and IQ less than 70 as measured by the Mehrfachwahl Wortschatz Test Form B. |
| **Singapore** | Singapore | SCID DSM-IV | Inclusion criteria include: - 1) DSM IV diagnosis of SZ (Patients) 2) Age: 21- 65 3) English speaking 4) Provision ofinformed written consent | Exclusion criteria include: - 1) History of significant head injury 2) Significant Neurological diseases (such as epilepsy, cerebrovascular accident) or Medical Illnesses 4) Significant DSM IV alcohol or substance use or dependence 6) Contraindications to MRI (e.g., pacemaker, orbital foreign body, recent surgery/procedure with metallic devices/implants deployed) 7) Pregnant women 8) Claustrophobia |
| **SWIFT** | Switzerland | SCID DSM-IV-TR | Age between 18-65 y/o, good German language skills that allowed them to understand the consent procedure and to undergo the clinical assessment, right-handedness according to the Edinburgh Inventory. For patients: Diagnosis with a schizophrenia spectrum disorder. | Participants were excluded if they were left-handed, pregnant, showed any contraindications for MRI (e.g. metal-containing implants such as pacemaker or cochlear implants, claustrophobia), had a history of serious neurological issues, or reported current abuse of alcohol and/or psychoactive substances (apart from nicotine). Additionally, controls had no current major psychiatric DSM-IV Axis I diagnoses, as assessed with the screening questionnaire of the Structured Clinical Interview for DSM-IV Axis I Disorders. |
| **UCISZ** | USA | SCID DSM-IV-TR criteria | All subjects diagnosed with schizophrenia were clinically stable outpatients whose antipsychotic medications and doses had not changed within the last two months. | Schizophrenia and healthy volunteers with a history of major medical illness, drug dependence in the last five years (except for nicotine), current substance abuse disorder, or MRI contraindications, were excluded. Individuals with schizophrenia who had significant tardive dyskinesia and healthy volunteers with a current or past history of major neurological or psychiatric illness or with a first-degree relative with an Axis-I psychotic disorder diagnosis were also excluded. |
| **UPenn** | USA | SCID DSM-IV-TR criteria | A DSM diagnosis of schizophrenia or schizoaffective disorder | Participants were excluded if they had a history of major medical illness that could impact brain function, active substance misuse, or a contraindication to MRI. |
| **Zurich** | Switzerland | MINI DSM IV | A diagnosis of schizophrenia | We excluded patients with any other DSM-IV Axis I disorder (in particular, current substance use disorder and major depressive disorder), those medicated with lorazepamat a dose higher than 1 mg, those with florid psychotic symptoms (i.e., any positive subscale item scores higher than 4 on the PANSS scale and those with extrapyramidal side effects (i.e., a total score higher than 2 on the MSAS). Healthy controls were screened for any neuropsychiatric disorders using the structured Mini-International Neuropsychiatric Interview to ensure that they had no previous or present psychiatric illness. Both patients and healthy controls were required to have a normal physical and neurologic status and no history of major head injury or neurologic disorder. |
| Cohort | **Country** | **Diagnosis measurement** | **Sample characteristics/inclusion criteria** | **Exclusion criteria** |

SCID: Structured Clinical Interview for DSM Disorders; CAMH: Comprehensive Assessment of Symptoms and History; MINI DSM IV: Mini-International Neuropsychiatric Interview for DSM IV; OPCRIT: Operational Criteria Checklist for Psychotic Illness and Affective Illness; DIP: Diagnostic Interview for Psychosis; ICD: international classification of disease; PANSS: Positive and Negative Syndrome Scale; MSAS: Modified Simpson–Angus Scale.

**Table S2: Site demographics**

| Site | N total | N SCZ | N HC | Age SZ | Age HC | %M/%F SCZ | %M/% F HC | Mean Duration Illness SZ | PANSS Positive | PANSS Negative | SAPS Total | SANS Total |
|---|---|---|---|---|---|---|---|---|---|---|---|---|
| ASRB | 429 | 263 | 166 | 38.6 | 39.3 | 67.3/32.7 | 47.6/52.4 | 15 | NA | NA | NA | 18.5 |
| CAMH | 264 | 118 | 146 | 43.9 | 43.6 | 59.3/40.7 | 52.7/47.3 | 19.2 | 13.9 | 14 | NA | NA |
| CIAM | 51 | 21 | 30 | 31 | 26.6 | 61.9/38.1 | 53.3/46.7 | 8.3 | 13.6 | 15.2 | NA | NA |
| COBRE | 143 | 73 | 70 | 37.4 | 35.7 | 82.2/17.8 | 71.4/28.6 | 15.8 | 15.2 | 14.8 | NA | NA |
| ESO | 80 | 40 | 40 | 29.4 | 29.1 | 50/50 | 50/50 | 0.6 | 14.2 | 16.1 | NA | NA |
| fidmag | 283 | 160 | 123 | 39.6 | 37.5 | 77.5/22.5 | 43.9/56.1 | 15.5 | 16.8 | 22.6 | NA | 37.2 |
| FOR210Marburg | 403 | 37 | 366 | 37.2 | 34 | 62.2/37.8 | 39.1/60.9 | 15.9 | NA | NA | 13.2 | 18.8 |
| FOR210Muenster | 163 | 8 | 155 | 33.4 | 27 | 50/50 | 38.7/61.3 | 11.1 | NA | NA | 6.4 | 8.1 |
| FSL_Rome | 280 | 164 | 116 | 39.4 | 37.5 | 67.1/32.9 | 62.9/37.1 | 14.9 | 20.9 | 21 | 31.5 | 28.8 |
| GIPSI | 43 | 43 | 0 | 33.5 | NA | 81.4/18.6 | NA/NA | 14.1 | NA | NA | 9.3 | 32.2 |
| IGP | 138 | 68 | 70 | 41.7 | 36 | 58.8/41.2 | 54.3/45.7 | 18.8 | 13.8 | 14.5 | 13 | 6.9 |
| MCIC | 213 | 117 | 96 | 33.9 | 32.7 | 74.4/25.6 | 67.7/32.3 | 11.1 | NA | NA | NA | NA |
| MPRC | 500 | 230 | 270 | 36.4 | 37.1 | 61.3/38.7 | 43.7/56.3 | NA | NA | NA | NA | NA |
| OLIN | 523 | 135 | 388 | 36.6 | 37.8 | 66.7/33.3 | 47.7/52.3 | 14.5 | 7.7 | 7.9 | NA | NA |
| PAFIP1.5T | 222 | 142 | 80 | 29.7 | 27.7 | 62/38 | 62.5/37.5 | 1 | NA | NA | 13.6 | 6.4 |

| | | | | | | | | | | | | |
|---|---|---|---|---|---|---|---|---|---|---|---|---|
| PAFIP3T | 217 | 114 | 103 | 29.7 | 30.1 | 56.1/43.9 | 60.2/39.8 | 0.7 | NA | NA | 14.1 | 5.5 |
| PENS | 51 | 17 | 34 | 48.4 | 46.8 | 70.6/29.4 | 44.1/55.9 | 24.7 | NA | NA | 12.2 | 17.8 |
| PHCP | 129 | 47 | 82 | 43.9 | 43.8 | 70.2/29.8 | 43.9/56.1 | 18.4 | NA | NA | 13.9 | 23.4 |
| RSCZ_data | 98 | 46 | 52 | 22.2 | 22.3 | 100/0 | 100/0 | 1.1 | 11.2 | 18.5 | NA | NA |
| SCORE | 127 | 72 | 55 | 26.9 | 26 | 72.2/27.8 | 45.5/54.5 | NA | NA | NA | NA | 8.15 |
| Singapore | 227 | 151 | 76 | 33.1 | 31.8 | 69.5/30.5 | 61.8/38.2 | 6.5 | 10.6 | 9 | NA | NA |
| STGO | 170 | 85 | 85 | 19.8 | 23.1 | 82.4/17.6 | 68.2/31.8 | 0.1 | 16.1 | 21.3 | NA | NA |
| SWIFT | 37 | 24 | 13 | 34.2 | 29.3 | 70.8/29.2 | 38.5/61.5 | 9.5 | 16.4 | 12.8 | NA | NA |
| UCISZ | 57 | 27 | 30 | 42.9 | 41.4 | 81.5/18.5 | 76.7/23.3 | 17.5 | 15.6 | 16 | 13.4 | 22.8 |
| UPenn | 370 | 177 | 193 | 38.9 | 36.4 | 59.3/40.7 | 46.6/53.4 | 17.3 | NA | NA | 18.3 | 23.7 |
| Zurich | 88 | 60 | 28 | 30.5 | 32.5 | 75/25 | 64.3/35.7 | 8.4 | 10.7 | 14.5 | NA | 24.9 |

#### Table S3. Sample image acquisition and image pre-processing details by cohort

| Cohort | Number of scanners | Scanner Vendor & Type | Imaging Protocols | Slice orientation | FreeSurfer Version | Operating System | Number of subjects removed from analysis due to QC failure with reasons |
|---|---|---|---|---|---|---|---|
| ASRB | 5 | Siemens<br>Avanto<br>1.5T | High-resolution T1-weighted structural magnetic resonance imaging (sMRI) brain scans (MPRAGE) were acquired using an optimized magnetization prepared rapid acquisition gradient echo on 1.5 T Siemens Avanto scanners (Siemens, Erlangen, Germany) across five Australian research sites (Loughland and al., 2010). Image parameters were set to 176 slices of 1mm thickness, no gap with field-of view 250 x 250 mm2, repetition time 1980 ms, echo time 4.3 ms, data acquisition matrix 256 x 256, with a flip matrix of 15°, resulting in a voxel size of 0.98×0.98×1.0 mm3 | Sagittal | v5.1.0 | Mac OSX | 0 |
| CAMH | 1 | GE 1.5T | SPGR, TR/TE/TI=12.3/5.3/300ms, flip angle=20°, 256x256x128 matrix, FOV=240x240mm, slice thickness=1.5mm | Axial | v5.3.0 | ubuntu x86_64-Linux | 0 |
| CASSI | 1 | 3-Tesla Philips Achieva with an 8-channel head coil | T1 weighted high-resolution anatomical scans were obtained (1mm slice thickness, no gap, 180 slices, TR = 5.4ms, TE = 2.4ms, field of view 256mm) | Sagittal | v.5.1.0 | Mac OSX 10.8 | Scans from three participants were excluded for motion artifact (1 control and 2 patients), and two participants due to neuroanatomical abnormalities (one healthy control showing a meningioma and one patient with agenesis of the corpus callosum) |

| | | | | | | | |
|---|---|---|---|---|---|---|---|
| COBRE | 1 | 3T Siemens TIM Trio | T1-weighted images were acquired with a 5-echo multi-echo MPRAGE sequence [TE (echo times) = 1.64, 3.5, 5.36, 7.22, 9.08 ms, TR (repetition time) = 2.53 s, TI (inversion time) = 1.2 s, 7○ flip angle, number of excitations (NEX) = 1, slice thickness = 1 mm, FOV (field of view) = 256 mm, resolution = 256x256] | Sagittal | v5.3.0 | Linux RedHat | A total of 9 participants were excluded following ENIGMA QA protocol.<br><br>9 total: 6 SCZ, 3 HC; 8 Males; average age: 37.89 (SD = 10.30). age range: 25-52 y.o. |
| EONCKS | 1 | 3T siemens allegra | MPRAGE sequence, 2080 ms repetition time; 4.88 ms echo-time, Field of view: 230 mm, 176 slices, 0.9 × 0.9 × 1 mm3 voxel size | Sagittal | v 6.0 | Linux | Nineteen patients and 11 controls were excluded due to motion artefacts or a technical scanner error.<br>Fourteen patients and sixteen controls were excluded due to motion artefacts or a technical scanner error. Of the N=30 excluded participants, n=22 were males, n=8 were controls, Mean age [range] across all excluded participants were: 26.18 years [16-39]. |
| ESO | 1 | 3T Siemens Tim Trio | MP-RAGE 3D, 1mm thickness, acquisition matrix 256 x 256, TR=2300ms, TE=4.63ms, TI=900ms | Sagittal | v5.3.0 | Linux | Only subjects without significant motion artifacts (assessed by visual inspection) were included.<br>Apart from ENIGMA QA protocol, visual inspection of all slices and edits to the skullstrip, white matter segmentation and control points insertion for correction of signal intensity normalization were done where needed. No subjects were excluded. |

| | | | | | | | |
|---|---|---|---|---|---|---|---|
| FBIRN | 7 | 3T Siemens Tim Trio; 3T GE | High-resolution structural imaging scans were acquired on six 3T Siemens Tim® Trio System and one 3T General Electric Discovery MR750 scanner. MP-RAGE scan parameters for the Siemens scanner were scan plane=sagittal, TR/TE/TI=2300/2.94/1100ms, GRAPPA acceleration factor=2, flip angle=9°, resolution=256×256x160, FOV=220mm2, voxel size=0.86x0.86x1.2mm, and NEX=1. IR-SPGR scan parameters for the General Electric scanner were scan plane=sagittal, TR/TE/TI=5.95/1.99/450ms, ASSET acceleration factor=2, a flip angle=12°, resolution=256×256x166, FOV=220mm2, voxel size=0.86x0.86x1.2mm, and NEX=1. All scans covered the entire brain. | Sagittal | v5.3.0 | Centos 64bit | 0 |
| FIDMAG | 1 | 1.5T GE Signa | 180 axial slices; 1mm slice thickness, no gap, matrix size 512x512; 0.5x0.5x1mm3 voxel resolution; TE 4ms, TR 2000ms, flip angle 15 | Axial | v5.3.0 | Linux Ubuntu | 0 |
| FSLRome | 1 | Siemens 3T Allegra | 3D MPRAGE: TE/TR=2.4/7.92 ms, flip angle=15°, voxel size 1×1×1 mm | Sagittal | 6.0dev | Linux | 0 |
| IGP | 1 | Philips 3T Achieva TX | TR 8.9ms, TE 4.1ms, field of view 240mm, matrix 268 x 268, 200 sagittal slices, slice thickness 0.9mm, no gap | Sagittal | v5.3.0 | Mac OSX | 0 |
| IMH Singapore | 1 | 3T Philips Achieva | T1 scans: 180 axial slices of 0.9mm thickness with no gap, FOV = 230x230 mm2, matrix 256x204, voxel size =0.89x0.89x0.9 mm3, TR=7.2 s, TE=3.3 ms, FA=8°, | Axial | v5.3.0 | Mac OSX | 2 due to excessive motion artifacts, SZ=1 (male, age 33 years), HC=1 (male, 31 years) |
| Madrid | 1 | Philips Intera 1.5T | T1 scans; TR 25ms; TE 9.18ms; voxel size 1x0.94x0.94mm$^3$; FOV 240x240m$^2$; 175 slices; flip angle 30; Gradient Echo (FFE 3d) | Sagittal | v5.3.0 | Linux | 0 |

| | | | | | | | |
|---|---|---|---|---|---|---|---|
| MCIC | 3 | 1.5, 3T Siemens and GE | T1 scans: TR = 2530 ms for 3 T, TR = 12 ms for 1.5 T; TE = 3.79 ms for 3 T, TE = 4.76 ms for 1.5 T; FA = 7 for 3 T, FA = 20 for 1.5 T; TI = 1100 for 3 T; Bandwidth = 181 for 3 T, Bandwidth = 110 for 1.5 T; 0.625×0.625 mm voxel size; slice thickness 1.5 mm; FOV 256×256×128 cm matrix; FOV = 16 cm (could be increased to 18 cm when needed for full brain coverage). | Coronal | v4.0.1 | Linux of various flavors | 5 subjects failed automated segmentation procedure due to excessive motion artifacts 2 participants' MRI data failed the manual inspection |
| MPRC | 2 | MPRC 1: 3T Siemens Allegro<br><br>MPRC 2: 3T Siemens Trio | MPRC 1: T1-weighted, 3D MPRAGE, 1x1x1mm, TE/TR/TI=4.3/2500/1000ms, flip angle=8 degrees.<br><br>MPRC 2**:** T1-weighted, 3D MPRAGE, 1x1x1mm, TE/TR/TI=2.9/2300/900ms, flip angle=9 degrees. | Sagittal | v5.3.0 | Linux | 0 |
| PAFIP | 2 | GE 1.5T | Three-dimensional T1-weighted images, using a spoiled grass (SPGR) sequence acquired in the coronal plane with: echo time (TE)=5 ms, repetition time (TR)=24 ms, numbers of excitations (NEX)=2, rotation angle=45°, field of view (FOV)=26×19.5 cm, slice thickness=1.5mm and a matrix of 256×192. | Coronal | v5.0.0 | Ubuntu 11,04 (x86_64) | 1 subject was excluded because motion artifacts resulted in very poor segmentation |
| RSCZ | 1 | 3T Philips Achieva | A turbo field echo sequence covering the whole brain. TR = 8.2 ms, TE = 3.7 ms, flip angle = 8, FOV = 240 mm, voxel size of 0.83 × 0.83 mm with a slice thickness of 1 mm, no gap. | Sagittal | v5.3.0 | Centos 6.6 | 0. Only subjects without significant motion and other artifacts (assessed by visual inspection) were included for carrying out ENIGMA QA protocol. |
| SCORE | 1 | 3T Philips Achieva | MPRAGE: acquisition matrix: 256×256×176, isotropic spatial resolution: 1x1x1mm3, TI=1000ms, TR=2s, TE=3.4 ms, flip angle: 8° and bandwidth of 200 Hz/pixel | Sagittal | 6.0dev | ubuntu 18.04 LTS | From the initial data set, 11 subjects had to be excluded based on erroneous brain segmentation. Among them, 4 subjects revealed statistical outliers. |
| SNUH | 1 | 3T Siemens Trio | high-resolution T1-weighted, three-dimensional Magnetization Prepared Rapid Gradient Echo (TR = 670ms; TE=1.89ms; FOV=250mm; FA=9°; voxel size=1x1x1mm3) | Sagittal | v5.3.0 | OSX 10.9 | 0 |

| | | | | | | | |
|---|---|---|---|---|---|---|---|
| SWIFT | 1 | 3T Siemens Verio | MPRAGE: 160 sagittal slices, 1mm slice thickness, 256x256 matrix size, 1x1x1 mm3 voxel size. TR = 2.3ms, TE = 2.98ms, TI = 900ms. | Sagittal | v 7.1.1 | Linux | 0 |
| UCISZ | 1 | 3T Philips Achieva | T1TFE:200 sagittal slices, 320x274 matrix size, .75mm isotropic, TR = 11ms, TE=4.562ms, flip angle = 18°, | Sagittal | v6.0dev | Centos 3.10.72-1.el6.elrepo.x86_64 | 0 |
| UNIBA | 1 | 3T GE | T1-weighted 3D FFE, TR/TE 9.86/4.6ms, 0.875x0.875x1 voxels, flip angle 8, FOV 224x160x168, 160 slices | Axial | v5.3.0 | Ubuntu 10.04, Kernel Linux 2.6.32-25-generic, GNOME 2.30.2 | 3 due to motion artefacts |
| Zurich | 1 | 3T Philips | 3D T1-weighted images were acquired with an ultra-fast gradient echo T1-weighted sequence (TR=8.4ms, TE=3.8ms, flip angle=8°) in 160 sagittal plan slices (1mm slice thickness, no slice gap) of 240×240mm2 resulting in 1x1x1mm3voxels. | Sagittal | v6.0.0 | Linux | 0 |

**HCP MRI data and preprocessing**

HCP data were acquired on a Siemens Skyra 3T and included (i) T1-weighted images [magnetization-prepared rapid gradient echo sequence, repetition time (TR) = 2400 ms, echo time (TE) = 2.14 ms, field of view (FOV) = 224 × 224 mm2, voxel size = 0.7 mm3, 256 slices], (ii) resting-state fMRI [gradient-echo echo-planar imaging (EPI) sequence, TR = 720 ms, TE = 33.1 ms, FOV = 208 × 180 $mm^2$, voxel size = 2 mm3, 72 slices], and (iii) diffusion MRI (spin-echo EPI sequence, TR = 5520 ms, TE = 89.5 ms, FOV = 210 × 180$mm^2$, voxel size = 1.25 mm3, b-value = 1000/2000/3000 s/mm2, 270 diffusion directions, 18 b0 images). HCP data underwent the initiative's minimal preprocessing (1, 2). Resting-state fMRI data underwent distortion and head motion corrections, magnetic field bias correction, skull removal, intensity normalization, and were mapped to MNI152 space. Noise components attributed to head movement, white matter, cardiac pulsation, arterial, and large vein–related contributions were automatically removed using ICA-FIX (3). Preprocessed time series were mapped to standard gray ordinate space using a cortical ribbon-constrained volume-to-surface mapping algorithm and subsequently concatenated to form a single time series. Diffusion MRI data underwent b0 intensity normalization and correction for susceptibility distortion, eddy currents, and head motion. High-resolution functional and structural data were parcellated according to the Desikan-Killiany atlas to align with the ENIGMA-Schizophrenia dataset (3).

**Functional and structural connectivity matrix generation from HCP participants**

Functional connectivity matrices were generated by computing pairwise correlations between the time series of all 68 cortical regions and between all subcortical and cortical regions; negative connections were set to zero. Subject-specific connectivity matrices were then *z*-transformed and aggregated across participants to construct a group-average functional connectome. To generate structural connectivity matrices, constrained tractography was performed using different tissue types derived from the T1w image, including cortical and subcortical gray matter, white matter, and cerebrospinal fluid (4). Multishell and multi-tissue response functions were estimated (5) and constrained spherical deconvolution and intensity normalization were performed (6, 7). The initial tractogram was generated with 40 million streamlines, with a maximum tract length of 250 and a fractional anisotropy cutoff of 0.06. Spherical-deconvolution informed filtering of tractograms (SIFT2) was applied to reconstruct whole-brain streamlines weighted by the cross-sectional multipliers (6). To produce normative subject-specific connectivity matrices according to the Desikan-Killiany atlas, we mapped reconstructed streamlines onto the 68 cortical and 14 subcortical (including hippocampus) regions (8). Normative structural connectivity matrices were generated from preprocessed diffusion MRI data using MRtrix3 (9). The group-average normative structural connectome was defined using a distance-dependent thresholding, which preserved the edge length distribution in individual participants (10), and was log transformed to reduce connectivity strength variance. Hence, structural connectivity was defined by the number of streamlines between two regions (*i.e.*, fiber density). Our network centrality findings of healthy individuals reflect previously published results (11, 12) with centrality peaking in medial prefrontal, superior parietal and angular regions.

**ComBat batch effect adjustment**

In our original mega-analytic model, regional cortical thickness and subcortical volumes were the dependent variables whilst group (SCZ, HC), age and sex were the independent variables. To show that our batch-effect correction with ComBat on the structural MRI data was successful, we performed our mega-analytical linear regression with the ComBat adjusted and unadjusted data similar with the original model, hereby adding site as an independent variable. For each region, we then computed the variance explained by site as independent variable quantified by partial $R^2$. Partial $R^2$ was computed using the *rsq.partial* function of the *rsq* library in R. It can be seen on Table S4 that data adjustment with ComBat eliminates any variance explained by site as an independent variable, i.e., variation of imaging data due to site differences.

**Table S4 ComBat batch effect adjustment confirmation**

| Brain region | partial $R^2$ of site on ComBat-adjusted data | | partial $R^2$ of site on ComBat unadjusted data | |
|---|---|---|---|---|
| | **Left** | **Right** | **Left** | **Right** |
| bankssts_thickavg | 0 | 0 | 0.29 | 0.30 |
| caudalanteriorcingulate_thick | 0 | 0 | 0.25 | 0.26 |
| caudalmiddlefrontal_thickavg | 0 | 0 | 0.31 | 0.30 |
| cuneus_thickavg | 0 | 0 | 0.28 | 0.26 |
| entorhinal_thickavg | 0 | 0 | 0.21 | 0.20 |
| fusiform_thickavg | 0 | 0 | 0.42 | 0.42 |
| inferiorparietal_thickavg | 0 | 0 | 0.29 | 0.36 |
| inferiortemporal_thickavg | 0 | 0 | 0.38 | 0.42 |
| isthmuscingulate_thickavg | 0 | 0 | 0.21 | 0.15 |
| lateraloccipital_thickavg | 0 | 0.001 | 0.32 | 0.35 |
| lateralorbitofrontal_thickavg | 0 | 0 | 0.24 | 0.24 |
| lingual_thickavg | 0 | 0 | 0.28 | 0.21 |
| medialorbitofrontal_thickavg | 0 | 0 | 0.30 | 0.29 |
| middletemporal_thickavg | 0 | 0 | 0.31 | 0.37 |
| parahippocampal_thickavg | 0 | 0 | 0.18 | 0.24 |
| paracentral_thickavg | 0 | 0 | 0.18 | 0.20 |
| parsopercularis_thickavg | 0 | 0 | 0.26 | 0.27 |
| parsorbitalis_thickavg | 0 | 0 | 0.13 | 0.16 |
| parstriangularis_thickavg | 0 | 0 | 0.21 | 0.26 |
| pericalcarine_thickavg | 0 | 0 | 0.42 | 0.36 |
| postcentral_thickavg | 0 | 0 | 0.30 | 0.30 |
| posteriorcingulate_thickavg | 0 | 0 | 0.23 | 0.21 |
| precentral_thickavg | 0 | 0 | 0.29 | 0.23 |
| precuneus_thickavg | 0 | 0 | 0.27 | 0.25 |
| rostralanteriorcingulate_thic | 0 | 0 | 0.28 | 0.29 |

| | | | | |
|---|---|---|---|---|
| rostralmiddlefrontal_thickavg | 0 | 0 | 0.34 | 0.39 |
| superiorfrontal_thickavg | 0 | 0 | 0.31 | 0.28 |
| superiorparietal_thickavg | 0 | 0 | 0.34 | 0.35 |
| superiortemporal_thickavg | 0 | 0 | 0.34 | 0.36 |
| supramarginal_thickavg | 0 | 0 | 0.28 | 0.33 |
| frontalpole_thickavg | 0 | 0 | 0.17 | 0.17 |
| temporalpole_thickavg | 0 | 0 | 0.20 | 0.26 |
| transversetemporal_thickavg | 0 | 0 | 0.15 | 0.17 |
| insula_thickavg | 0 | 0 | 0.34 | 0.35 |

**Table S5. Cortical thickness differences between schizophrenia patients and healthy controls**

| Regions | T-values | p values (Bonferroni-corrected) | Cohen's D | Cohen's D 95%CI |
|---|---|---|---|---|
| Left banks of superior temporal sulcus | 12.26 | $1.44 \times 10^{-34}$ | 0.34 | [0.28, 0.39] |
| Left caudal anterior cingulate cortex | 3.01 | $8.95 \times 10^{-4}$ | 0.08 | [0.03, 0.14] |
| Left caudal middle frontal gyrus | 13.06 | $7.69 \times 10^{-39}$ | 0.36 | [0.30, 0.41] |
| Left cuneus | 6.71 | $7.21 \times 10^{-12}$ | 0.18 | [0.13, 0.24] |
| Left entorhinal cortex | 5.22 | $6.38 \times 10^{-8}$ | 0.14 | [0.09, 0.20] |
| Left fusiform gyrus | 14.46 | $5.42 \times 10^{-47}$ | 0.40 | [0.34, 0.45] |
| Left inferior parietal cortex | 14.38 | $1.64 \times 10^{-46}$ | 0.40 | [0.34, 0.45] |
| Left inferior temporal gyrus | 14.29 | $5.61 \times 10^{-46}$ | 0.39 | [0.34, 0.45] |
| Left isthmus cingulate cortex | 8.29 | $4.86 \times 10^{-17}$ | 0.23 | [0.17, 0.28] |
| Left lateral occipital cortex | 11.7 | $1.00 \times 10^{-31}$ | 0.32 | [0.27, 0.38] |
| Left lateral orbitofrontal cortex | 12.2 | $2.99 \times 10^{-34}$ | 0.34 | [0.28, 0.39] |
| Left lingual gyrus | 9.72 | $1.29 \times 10^{-22}$ | 0.27 | [0.21, 0.32] |
| Left medial orbitofrontal cortex | 6.86 | $2.64 \times 10^{-12}$ | 0.19 | [0.13, 0.24] |
| Left middle temporal gyrus | 15.26 | $5.52 \times 10^{-52}$ | 0.42 | [0.36, 0.47] |
| Left parahippocampal gyrus | 5.88 | $1.52 \times 10^{-9}$ | 0.16 | [0.11, 0.22] |
| Left paracentral lobule | 9.97 | $1.09 \times 10^{-23}$ | 0.27 | [0.22, 0.33] |
| Left pars opercularis of inferior frontal gyrus | 13.13 | $3.04 \times 10^{-39}$ | 0.36 | [0.31, 0.41] |
| Left pars orbitalis of inferior frontal gyrus | 10.85 | $1.26 \times 10^{-27}$ | 0.30 | [0.24, 0.35] |
| Left pars triangularis of inferior frontal gyrus | 11.81 | $3.01 \times 10^{-32}$ | 0.32 | [0.27, 0.38] |
| Left pericalcarine cortex | 2.34 | $6.55 \times 10^{-3}$ | 0.06 | [0.01, 0.12] |
| Left postcentral gyrus | 12.63 | $1.63 \times 10^{-36}$ | 0.35 | [0.29, 0.40] |
| Left posterior cingulate cortex | 10.02 | $6.90 \times 10^{-24}$ | 0.28 | [0.22, 0.33] |
| Left precentral gyrus | 12.71 | $5.88 \times 10^{-37}$ | 0.35 | [0.29, 0.40] |
| Left precuneus | 11.57 | $4.82 \times 10^{-31}$ | 0.32 | [0.26, 0.37] |
| Left rostral anterior cingulate cortex | 4.51 | $2.27 \times 10^{-6}$ | 0.12 | [0.07, 0.18] |
| Left rostral middle frontal gyrus | 11.95 | $5.59 \times 10^{-33}$ | 0.33 | [0.27, 0.38] |
| Left superior frontal gyrus | 13.83 | $3.22 \times 10^{-43}$ | 0.38 | [0.33, 0.43] |
| Left superior parietal cortex | 9.87 | $3.02 \times 10^{-23}$ | 0.27 | [0.22, 0.33] |
| Left superior temporal gyrus | 14.57 | $1.20 \times 10^{-47}$ | 0.40 | [0.35, 0.45] |
| Left supramarginal gyrus | 15.74 | $4.77 \times 10^{-55}$ | 0.43 | [0.38, 0.49] |
| Left frontal pole | 6.76 | $5.16 \times 10^{-12}$ | 0.19 | [0.13, 0.24] |
| Left temporal pole | 7.29 | $1.18 \times 10^{-13}$ | 0.20 | [0.15, 0.25] |

| | | | | |
|---|---|---|---|---|
| Left transverse temporal gyrus | 9.16 | 2.50x10$^{-20}$ | 0.25 | [0.20, 0.31] |
| Left insula | 13.55 | 1.32x10$^{-41}$ | 0.37 | [0.32, 0.43] |
| Right banks of superior temporal sulcus | 12.12 | 7.72x10$^{-34}$ | 0.33 | [0.28, 0.39] |
| Right caudal anterior cingulate cortex | 5.05 | 1.56x10$^{-7}$ | 0.14 | [0.08, 0.19] |
| Right caudal middle frontal gyrus | 11.91 | 8.93x10$^{-33}$ | 0.33 | [0.27, 0.38] |
| Right cuneus | 7.64 | 8.96x10$^{-15}$ | 0.21 | [0.16, 0.26] |
| Right entorhinal cortex | 4.04 | 1.84x10$^{-5}$ | 0.11 | [0.06, 0.16] |
| Right fusiform gyrus | 15.12 | 4.64x10$^{-51}$ | 0.42 | [0.36, 0.47] |
| Right inferior parietal cortex | 13.03 | 1.02x10$^{-38}$ | 0.36 | [0.30, 0.41] |
| Right inferior temporal gyrus | 13.75 | 9.58x10$^{-43}$ | 0.38 | [0.32, 0.43] |
| Right isthmus cingulate cortex | 7.94 | 8.22x10$^{-16}$ | 0.22 | [0.16, 0.27] |
| Right lateral occipital cortex | 11.39 | 3.71x10$^{-30}$ | 0.31 | [0.26, 0.37] |
| Right lateral orbitofrontal cortex | 10.72 | 5.36x10$^{-27}$ | 0.29 | [0.24, 0.35] |
| Right lingual gyrus | 10.67 | 8.45x10$^{-27}$ | 0.29 | [0.24, 0.35] |
| Right medial orbitofrontal cortex | 7.45 | 3.69x10$^{-14}$ | 0.20 | [0.15, 0.26] |
| Right middle temporal gyrus | 13.9 | 1.16x10$^{-43}$ | 0.38 | [0.33, 0.44] |
| Right parahippocampal gyrus | 6.01 | 6.70x10$^{-10}$ | 0.17 | [0.11, 0.22] |
| Right paracentral lobule | 9.08 | 5.31x10$^{-20}$ | 0.25 | [0.20, 0.30] |
| Right pars opercularis of inferior frontal gyrus | 14.6 | 8.08x10$^{-48}$ | 0.40 | [0.35, 0.46] |
| Right pars orbitalis of inferior frontal gyrus | 10.31 | 3.72x10$^{-25}$ | 0.28 | [0.23, 0.34] |
| Right pars triangularis of inferior frontal gyrus | 12.16 | 4.97x10$^{-34}$ | 0.33 | [0.28, 0.39] |
| Right pericalcarine cortex | 2.32 | 6.87x10$^{-3}$ | 0.06 | [0.01, 0.12] |
| Right postcentral gyrus | 11.13 | 6.59x10$^{-29}$ | 0.31 | [0.25, 0.36] |
| Right posterior cingulate cortex | 9.34 | 4.50x10$^{-21}$ | 0.26 | [0.20, 0.31] |
| Right precentral gyrus | 11.72 | 8.23x10$^{-32}$ | 0.32 | [0.27, 0.38] |
| Right precuneus | 10.6 | 1.91x10$^{-26}$ | 0.29 | [0.24, 0.35] |
| Right rostral anterior cingulate cortex | 4.42 | 3.46x10$^{-6}$ | 0.12 | [0.07, 0.18] |
| Right rostral middle frontal gyrus | 11.53 | 7.03x10$^{-31}$ | 0.32 | [0.26, 0.37] |
| Right superior frontal gyrus | 12.82 | 1.57x10$^{-37}$ | 0.35 | [0.30, 0.41] |
| Right superior parietal cortex | 10.03 | 6.31x10$^{-24}$ | 0.28 | [0.22, 0.33] |
| Right superior temporal gyrus | 15.05 | 1.30x10$^{-50}$ | 0.41 | [0.36, 0.47] |
| Right supramarginal gyrus | 14.29 | 6.06x10$^{-46}$ | 0.39 | [0.34, 0.45] |
| Right frontal pole | 6.55 | 2.18x10$^{-11}$ | 0.18 | [0.13, 0.23] |
| Right temporal pole | 7.19 | 2.45x10$^{-13}$ | 0.20 | [0.14, 0.25] |
| Right transverse temporal gyrus | 9.4 | 2.77x10$^{-21}$ | 0.26 | [0.20, 0.31] |
| Right insula | 12.77 | 2.95x10$^{-37}$ | 0.35 | [0.30, 0.40] |

**Table S6. Subcortical volume differences between schizophrenia and healthy controls**

| Regions | T-values | p values (Bonferroni-corrected) | Cohen's D | Cohen's D 95%CI |
|---|---|---|---|---|
| Laccumb | 1.81 | 1 | 0.05 | [0,0.1] |
| Lamyg | 6.6 | 4.55x10-11 | 0.18 | [0.13,0.24] |
| Lcaud | -1.12 | 1 | -0.03 | [-0.08,0.02] |
| Lhippo | 12.32 | 2.05x10-34 | 0.34 | [0.28,0.39] |
| Lpal | -9.56 | 1 | -0.26 | [-0.32,-0.21] |

| | | | | |
|---|---|---|---|---|
| Lput | -4.41 | 1 | -0.12 | [-0.18,-0.07] |
| Lthal | 9.74 | 2.94x10-22 | 0.27 | [0.21,0.32] |
| Raccumb | 3.62 | 2.95x10-4 | 0.1 | [0.05,0.15] |
| Ramyg | 6.58 | 5.10x10-11 | 0.18 | [0.13,0.23] |
| Rcaud | -1.64 | 1 | -0.04 | [-0.1,0.01] |
| Rhippo | 12.52 | 1.75x10-35 | 0.34 | [0.29,0.4] |
| Rpal | -6.84 | 1 | -0.19 | [-0.24,-0.13] |
| Rput | -4.4 | 1 | -0.12 | [-0.17,-0.07] |
| Rthal | 9.91 | 5.60x10-23 | 0.27 | [0.22,0.33] |

**Table S7. Schizophrenia cortical epicenter ranking (ordered by significance of functional epicenters)**

| Regions | Functional Epicenter R values | Functional Epicenter $p_{spin}$ values (Bonferroni-corrected) | Structural Epicenter R values | Structural Epicenter $p_{spin}$ values (Bonferroni - corrected) |
|---|---|---|---|---|
| Left entorhinal cortex | 0.69 | <.001 | -0.08 | 1 |
| Left banks of superior temporal sulcus | 0.68 | <.001 | 0.38 | 0.163 |
| Left inferior temporal gyrus | 0.67 | <.001 | 0.24 | 1 |
| Right inferior temporal gyrus | 0.66 | <.001 | 0.19 | 1 |
| Right entorhinal cortex | 0.65 | <.001 | 0.01 | 1 |
| Right banks of superior temporal sulcus | 0.64 | <.001 | 0.29 | 1 |
| Left pars triangularis of inferior frontal gyrus | 0.63 | <.001 | 0.35 | 0.197 |
| Left pars opercularis of inferior frontal gyrus | 0.6 | <.001 | 0.45 | 0.007 |
| Right pars triangularis of inferior frontal gyrus | 0.58 | <.001 | 0.37 | 0.15 |
| Right lateral orbitofrontal cortex | 0.57 | <.001 | -0.11 | 1 |
| Left caudal middle frontal gyrus | 0.56 | <.001 | 0.39 | 0.122 |
| Left lateral orbitofrontal cortex | 0.55 | <.001 | -0.07 | 1 |
| Right pars orbitalis of inferior frontal gyrus | 0.54 | <.001 | 0.23 | 1 |
| Left rostral middle frontal gyrus | 0.54 | <.001 | 0.22 | 1 |
| Left supramarginal gyrus | 0.54 | <.001 | 0.36 | 0.435 |
| Left pars orbitalis of inferior frontal gyrus | 0.54 | <.001 | 0.21 | 1 |
| Right caudal middle frontal gyrus | 0.51 | <.001 | 0.33 | 0.653 |
| Left superior temporal gyrus | 0.51 | <.001 | 0.22 | 1 |
| Right superior frontal gyrus | 0.5 | <.001 | -0.12 | 1 |
| Right middle temporal gyrus | 0.5 | <.001 | 0.15 | 1 |
| Left superior frontal gyrus | 0.49 | <.001 | -0.09 | 1 |
| Left middle temporal gyrus | 0.48 | <.001 | 0.32 | 0.313 |
| Right superior temporal gyrus | 0.47 | <.001 | 0.26 | 1 |
| Right pars opercularis of inferior frontal gyrus | 0.46 | .001 | 0.22 | 1 |
| Right inferior parietal cortex | 0.45 | .001 | 0.34 | 0.333 |
| Right rostral middle frontal gyrus | 0.44 | .001 | 0.19 | 1 |
| Left inferior parietal cortex | 0.44 | .001 | 0.39 | 0.143 |
| Left temporal pole | 0.44 | .002 | 0.21 | 1 |
| Left transverse temporal gyrus | 0.43 | .002 | 0.21 | 1 |
| Right supramarginal gyrus | 0.43 | .004 | 0.28 | 1 |

| | | | | |
|---|---|---|---|---|
| Right transverse temporal gyrus | 0.41 | .007 | 0.21 | 1 |
| Left precentral gyrus | 0.39 | .008 | 0.37 | 0.299 |
| Left posterior cingulate cortex | 0.38 | .009 | -0.19 | 1 |
| Left caudal anterior cingulate cortex | 0.38 | .009 | -0.13 | 1 |
| Left insula | 0.37 | .01 | 0.05 | 1 |
| Right precentral gyrus | 0.37 | .012 | 0.31 | 1 |
| Left fusiform gyrus | 0.37 | .012 | -0.01 | 1 |
| Right insula | 0.36 | .014 | 0.01 | 1 |
| Left paracentral lobule | 0.36 | .017 | 0.06 | 1 |
| Right paracentral lobule | 0.35 | .017 | 0.11 | 1 |
| Left superior parietal cortex | 0.35 | .017 | 0.2 | 1 |
| Right postcentral gyrus | 0.34 | .017 | 0.25 | 1 |
| Right caudal anterior cingulate cortex | 0.34 | .019 | -0.14 | 1 |
| Right posterior cingulate cortex | 0.34 | .019 | -0.14 | 1 |
| Right temporal pole | 0.33 | .02 | 0.08 | 1 |
| Right fusiform gyrus | 0.32 | .021 | 0.01 | 1 |
| Right superior parietal cortex | 0.32 | .03 | 0.25 | 1 |
| Left postcentral gyrus | 0.31 | .033 | 0.28 | 1 |
| Left parahippocampal gyrus | 0.29 | .071 | -0.08 | 1 |
| Right parahippocampal gyrus | 0.29 | .075 | -0.11 | 1 |
| Right lateral occipital cortex | 0.24 | .081 | 0.07 | 1 |
| Left lateral occipital cortex | 0.24 | .084 | 0.03 | 1 |
| Right frontal pole | 0.24 | .084 | 0 | 1 |
| Right precuneus | 0.21 | .142 | -0.21 | 1 |
| Left frontal pole | 0.21 | .182 | -0.06 | 1 |
| Left precuneus | 0.18 | .242 | -0.24 | 1 |
| Left isthmus cingulate cortex | 0.13 | .394 | -0.27 | 1 |
| Right isthmus cingulate cortex | 0.11 | .48 | -0.26 | 1 |
| Left pericalcarine cortex | 0.11 | .48 | -0.07 | 1 |
| Right pericalcarine cortex | 0.1 | .538 | -0.14 | 1 |
| Left cuneus | 0.08 | .637 | -0.18 | 1 |
| Left medial orbitofrontal cortex | 0.07 | .651 | -0.09 | 1 |
| Left lingual gyrus | 0.06 | .749 | -0.08 | 1 |
| Right cuneus | 0.05 | .766 | -0.16 | 1 |
| Right lingual gyrus | 0.04 | .773 | -0.2 | 1 |
| Left rostral anterior cingulate cortex | 0.02 | .8 | -0.2 | 1 |
| Right medial orbitofrontal cortex | -0.01 | .903 | -0.17 | 1 |
| Right rostral anterior cingulate cortex | -0.05 | .958 | -0.16 | 1 |

**Table S8. Schizophrenia subcortical epicenter ranking**

| Regions | Functional Epicenter R values | Functional Epicenter $p_{spin}$ values (Bonferroni-corrected) | Structural Epicenter R values | Structural Epicenter $p_{spin}$ values (Bonferroni - corrected) |
|---|---|---|---|---|
| L_Accumbens | -0.07 | 1 | -0.16 | 1 |
| L_Amygdala | 0.53 | <0.001 | 0.03 | 1 |
| L_Caudate | 0.47 | 0.001 | 0.16 | 1 |
| L_Hippocampus | 0.34 | 0.235 | -0.05 | 1 |
| L_Pallidum | 0.42 | 0.014 | 0.21 | 1 |
| L_Putamen | 0.5 | 0.000 | 0.14 | 1 |
| L_Thalamus | 0.38 | 0.088 | -0.02 | 1 |
| R_Accumbens | -0.16 | 3.048 | -0.23 | 1 |
| R_Amygdala | 0.48 | 0.001 | -0.01 | 1 |
| R_Caudate | 0.43 | 0.010 | -0.01 | 1 |
| R_Hippocampus | 0.32 | 0.428 | -0.08 | 1 |
| R_Pallidum | 0.33 | 0.237 | 0.03 | 1 |
| R_Putamen | 0.46 | 0.001 | 0.13 | 1 |
| R_Thalamus | 0.33 | 0.309 | 0.18 | 1 |

**Table S9. Divergent and convergent regions of different stages of SCZ (Note, no unique epicenters were found for chronic SCZ)**

| Divergent Regions | | Convergent Regions |
|---|---|---|
| **First Episode Psychosis (FEP)** | **Early SCZ** | **FEP+Early SCZ + Chronic SCZ** |
| *Functional Connectivity* | *Functional Connectivity* | *Functional Connectivity* |
| Left cuneus | Left banks of superior temporal sulcus | Left inferior parietal cortex |
| Left fusiform gyrus | Left caudal middle frontal gyrus | Left inferior temporal gyrus |
| Left lateral occipital cortex | Left entorhinal cortex | Left lateral orbitofrontal cortex |
| Left lingual gyrus | Left middle temporal gyrus | Left pars opercularis of inferior frontal gyrus |
| Left pericalcarine cortex | Left pars orbitalis of inferior frontal gyrus | Left pars triangularis of inferior frontal gyrus |
| Left superior parietal cortex | Left posterior cingulate cortex | Left rostral middle frontal gyrus |
| Left superior temporal gyrus | Left superior frontal gyrus | Left supramarginal gyrus |
| Right caudal anterior cingulate cortex | Left insula | Right banks of superior temporal sulcus |
| Right lateral occipital cortex | Right caudal middle frontal gyrus | Right lateral orbitofrontal cortex |
| Right lingual gyrus | Right entorhinal cortex | Right pars triangularis of inferior frontal gyrus |
| Right pericalcarine cortex | Right inferior parietal cortex | |
| Right superior parietal cortex | Right inferior temporal gyrus | |
| Right transverse temporal gyrus | Right middle temporal gyrus | *Structural Connectivity* |
| | Right pars opercularis of inferior frontal gyrus | Left caudal middle frontal gyrus |
| *Structural Connectivity* | Right pars orbitalis of inferior frontal gyrus | Left pars opercularis of inferior frontal gyrus |
| Left banks of superior temporal sulcus | Right rostral middle frontal gyrus | Left precentral gyrus |
| Left cuneus | Right superior frontal gyrus | Right caudal middle frontal gyrus |
| Left isthmus cingulate cortex | | Right inferior parietal cortex |
| Left middle temporal gyrus | *Structural Connectivity* | Right pars triangularis of inferior frontal gyrus |
| Left pars orbitalis of inferior frontal gyrus | Right lingual gyrus | |
| Left temporal pole | Right pars opercularis of inferior frontal gyrus | |
| Right banks of superior temporal sulcus | Right pars orbitalis of inferior frontal gyrus | |
| Right superior temporal gyrus | | |
| Left pars orbitalis of inferior frontal gyrus | | |
| Left temporal pole | | |
| Right banks of superior temporal sulcus | | |
| Right superior temporal gyrus | | |

**Mega-analysis Robustness and sensitivity analyses**

*Reproducibility of cortical hub vulnerability across different centrality metrics*

To show that cortical hub vulnerability was reproducible across multiple network metrics of centrality, besides hub strength (degree centrality) we calculated the betweenness, eigenvector and closeness centralities of the 68x68 parcellated functional and structural cortical connectivity matrices of the HCP using R package "*NetworkToolbox*". All centrality metrics show a high degree of intercorrelation, (mean±SD: Rfunc = 0.69±0.31, Rstruc = 0.9±0.08, Table S9). The high intercorrelation is somewhat expected, given that different aspects of the same underlying construct are measured. Similar to our main analysis using hub strength (degree centrality) (see main methods hub vulnerability model section), we examined the correlation between the cortical alteration map of SCZ with the betweenness, eigenvector and closeness centrality of each region. Correlations between SCZ-related cortical alterations and the additional centrality measures revealed very similar results as observed with hub strength (degree centrality) (Table S11).

**Table S10. Correlation matrix of centrality measures**

| **Functional Connectivity** | | | | |
|---|---|---|---|---|
| | Strength | Eigenvector | Betweenness | Closeness |
| Strength | 1 | 1 | 0.41 | 0.98 |
| Eigenvector | 1 | 1 | 0.35 | 0.96 |
| Betweenness | 0.41 | 0.35 | 1 | 0.48 |
| Closeness | 0.98 | 0.96 | 0.48 | 1 |
| **Structural Connectivity** | | | | |
| | Strength | Eigenvector | Betweenness | Closeness |
| Strength | 1 | 0.98 | 0.86 | 0.92 |
| Eigenvector | 0.98 | 1 | 0.83 | 0.95 |
| Betweenness | 0.86 | 0.83 | 1 | 0.79 |
| Closeness | 0.92 | 0.95 | 0.79 | 1 |

**Table S11. Cortical hub vulnerability using different centrality metrics**

| Hub vulnerability | Centrality | Strength | | Eigenvector | | Betweenness | | Closeness | |
|---|---|---|---|---|---|---|---|---|---|
| | | R | $p_{spin}$ | R | $p_{spin}$ | R | $p_{spin}$ | R | $p_{spin}$ |
| Connectivity | Functional | 0.58 | <0.0001 | 0.54 | 0.0001 | 0.43 | <0.0001 | 0.63 | <0.0001 |
| | Structural | 0.32 | 0.02 | 0.26 | 0.04 | 0.36 | 0.01 | 0.27 | 0.04 |

*Reproducibility across HCP age-matched and age-divergent ENIGMA SCZ samples*

To show the robustness of our findings to the mean age group discrepancy of the multisite ENIGMA SCZ sample and the HCP sample, we split the ENIGMA SCZ sample into HCP age-matched groups and age-divergent groups. We first used the HCP mean age plus/minus 1SD to define an HCP age-matched ENIGMA SCZ sample (N = 1222, 495 SCZ) and an age-divergent group (N = 4084, 1944 SCZ). To show that the findings are robust across various definitions of age-matched groups, we also used the HCP mean age plus/minus 2SDs to define the HCP age-matched ENIGMA SCZ sample (N = 2651, 1119 SCZ) and an age-divergent groups (N = 2655, N = 1320 SCZ). Regarding the matching to the HCP sample by age mean ± 1SD, we observed a very high agreement between the resulting t-values ($r = 0.97$, $p < 2.2e^{-16}$), and functional ($r = 0.92$, $p < 2.2e^{-16}$) and structural ($r = 0.98$, $p < 2.2e^{-16}$) epicenters. We are also able to confirm our finding for the hub vulnerability of functional ($r = 0.49$, $p_{spin} = 2e^{-05}$ ) and structural nodes ($r = 0.26$, $p_{spin} = 0.027$). We find similar results when matching the sample by age using the mean ± 2SD, observing an even higher agreement between the resulting t-values ($r = 0.97$, $p < 2.2e^{-16}$), and functional ($r = 0.98$, $p < 2.2e^{-16}$) and structural ($r = 0.98$, $p < 2.2e^{-16}$) epicenters. We are also able to confirm our finding for the preferential vulnerability of functional ($r = 0.63$, $p = 1e-08$) and structural nodes ($r = 0.36$, $p = 0.003$) to cortical thickness reduction. Lastly, we tested whether our results would be replicated in the most age-dissimilar group, i.e., individuals outside of the (mean+-2SD) window of the HCP distribution. Again, we could replicate our original findings robustly, with high agreement of t-values ($r = 0.96$, $p < 2.2e^{-16}$), and functional ($r = 0.97$, $p < 2.2e^{-16}$) and structural ($r = 0.98$, $p < 2.2e^{-16}$) epicenters. We are also able to confirm our finding for the preferential vulnerability of functional ($r = 0.49$, $p_{spin} = 2e^{-05}$) and structural hubs ($r = 0.27$, $p_{spin} = 0.023$).

**Robustness and site-specific confirmation analysis of morphological alterations, hub vulnerability and disease epicenter models in SCZ**

*Cortical and subcortical alterations in SCZ*

To examine the reproducibility of our mega-analytic findings and to make sure that our findings were not outlier-driven, we repeated our analysis within each participating site separately. One of the participating sites (GIPSI) was excluded from this site-specific analysis since this site contains data of patients only. In a first step, we computed the t-value map of the cortical thickness differences between patients and controls within each participating site. Site-specific schizophrenia-related cortical alterations were similar to our multisite mega-analytical findings (Fig S1a). In addition, in 21 out of 25 sites the spatial pattern of the schizophrenia-related cortical t-value map was significant correlated with the mega-analytic (multisite aggregation) cortical t-value map indicating good spatial similarity between the cortical alteration maps (Table S13).

*Hub vulnerability and epicenter mapping*

We next examined within each participating site the reproducibility of our finding that more central cortical nodes tend to display higher values of cortical thickness reductions. The positive correlation of cortical thickness reduction with functional corticocortical degree centrality was more pronounced in functional networks (mean± SD: R= 0.23±0.28, Table S14 ) rather than structural networks (mean± SD: R= 0.15±0.14., Table S14), with 19 and 8 sites out of 25 showing statistical significance of this relationship respectively. In a final step, we examined the reproducibility of our epicenter findings, by identifying functional and structural epicenters of SCZ (see main methods for details) within each participating site. As observed in the multisite findings site-specific epicenters were most often identified in temporo-paralimbic extending to frontal brain regions (Fig S1b), with a high degree of correlation between our original map and the site-specific ones for functional, (median R=0.6; IQR=[0.26,0.8], Table S15) and structural epicenters (median R= 0.42; IQR=[0.25,0.55], Table S15)

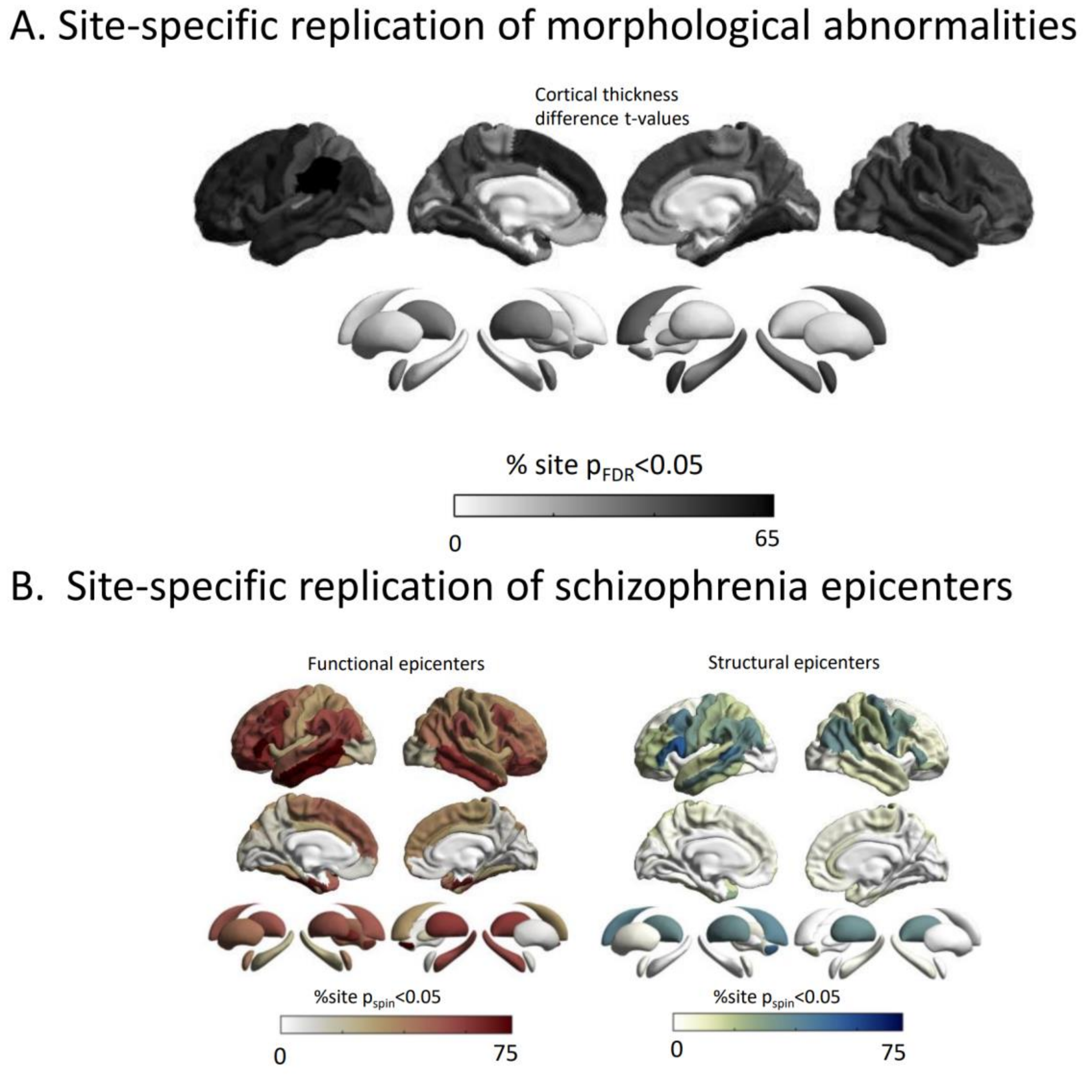


Figure S1. Site-specific replication of (A) morphological abnormalities in SCZ (B) functional and structural epicenters.

Table S13. Correlation between site-specific and mega-analytic cortical alteration maps of schizophrenia

| Sites | R | $p_{spin}$ |
|---|---|---|
| **ASRB** | 0.53 | <0.001 |
| **fidmag** | 0.58 | <0.001 |
| **FSL_Rome** | 0.25 | 0.010 |
| **IGP** | 0.06 | 0.304 |
| **Singapore** | 0.58 | <0.001 |
| **Zurich** | 0.33 | 0.005 |
| **COBRE** | 0.57 | <0.001 |
| **UCISZ** | 0.67 | <0.001 |
| **PAFIP3T** | 0.55 | <0.001 |
| **FOR210Marburg** | 0.68 | <0.001 |
| **FOR210Muenster** | 0.38 | 0.002 |
| **SWIFT** | 0.38 | 0.002 |
| **CAMH** | 0.36 | 0.002 |
| **PAFIP1.5T** | 0.64 | <0.001 |
| **STGO** | 0.76 | <0.001 |
| **SCORE** | 0.03 | 0.411 |
| **UPenn** | 0.48 | <0.001 |
| **PENS** | 0.20 | 0.062 |
| **PHCP** | 0.21 | 0.028 |
| **RSCZ_data** | 0.73 | <0.001 |
| **MCIC** | 0.76 | <0.001 |
| **ESO** | -0.35 | 0.999 |
| **CIAM** | 0.58 | <0.001 |
| **MPRC** | 0.69 | <0.001 |
| **OLIN** | 0.78 | <0.001 |

Table S14. Site-specific hub vulnerability analysis

| Sites | $R_{func}$ | $p_{func}$ | $R_{struc}$ | $p_{struc}$ |
|---|---|---|---|---|
| **ASRB** | -0.13 | 0.644 | 0.13 | 0.194 |
| **fidmag** | 0.39 | 0.006 | 0.04 | 0.367 |
| **FSL_Rome** | 0.60 | 0.010 | 0.40 | 0.001 |
| **IGP** | -0.31 | 0.865 | -0.05 | 0.641 |
| **Singapore** | 0.53 | <0.001 | 0.06 | 0.316 |
| **Zurich** | -0.14 | 0.695 | -0.14 | 0.828 |
| **COBRE** | -0.05 | 0.575 | 0.19 | 0.099 |
| **UCISZ** | 0.21 | 0.081 | 0.22 | 0.045 |
| **PAFIP3T** | 0.64 | 0.001 | 0.30 | 0.017 |
| **FOR210Marburg** | 0.42 | 0.002 | 0.23 | 0.041 |
| **FOR210Muenster** | -0.04 | 0.566 | 0.16 | 0.118 |
| **SWIFT** | 0.24 | 0.059 | 0.21 | 0.066 |
| **CAMH** | 0.34 | 0.073 | 0.17 | 0.124 |
| **PAFIP1.5T** | 0.28 | 0.029 | 0.16 | 0.104 |
| **STGO** | 0.62 | <0.001 | 0.23 | 0.053 |
| **SCORE** | 0.22 | 0.069 | -0.01 | 0.530 |
| **UPenn** | 0.22 | 0.097 | 0.12 | 0.179 |
| **PENS** | 0.13 | 0.191 | 0.22 | 0.052 |
| **PHCP** | 0.46 | 0.031 | -0.18 | 0.915 |
| **RSCZ_data** | 0.40 | 0.011 | 0.45 | 0.001 |
| **MCIC** | 0.34 | 0.012 | 0.20 | 0.066 |
| **ESO** | -0.35 | 0.953 | 0.12 | 0.187 |
| **CIAM** | 0.05 | 0.390 | 0.08 | 0.262 |
| **MPRC** | 0.55 | 0.002 | 0.20 | 0.079 |
| **OLIN** | 0.24 | 0.103 | 0.15 | 0.141 |

Table S15. Site-specific epicenter map agreement with mega-analytical epicenter map

| Sites | $R_{func}$ | $p_{func}$ | $R_{struc}$ | $p_{struc}$ |
|---|---|---|---|---|
| **ASRB** | 0.48 | 0 | 0.48 | 0 |
| **fidmag** | 0.8 | 0 | 0.44 | 0 |
| **FSL_Rome** | 0.17 | 0.132 | 0.1 | 0.124 |
| **IGP** | 0.15 | 0.132 | 0.25 | 0.115 |
| **Singapore** | 0.82 | 0 | 0.55 | 0 |
| **Zurich** | 0.45 | 0 | 0.31 | 0 |
| **COBRE** | 0.63 | 0 | 0.48 | 0 |
| **UCISZ** | 0.87 | 0 | 0.71 | 0 |
| **PAFIP3T** | 0.52 | 0 | 0.33 | 0 |
| **FOR210Marburg** | 0.77 | 0 | 0.54 | 0 |
| **FOR210Muenster** | 0.57 | 0 | 0.32 | 0 |
| **SWIFT** | 0.55 | 0 | 0.47 | 0 |
| **CAMH** | -0.03 | 0.587 | -0.25 | 0.598 |
| **PAFIP1.5T** | 0.83 | 0 | 0.56 | 0 |
| **STGO** | 0.86 | 0 | 0.56 | 0 |
| **SCORE** | -0.12 | 0.832 | -0.07 | 0.845 |
| **UPenn** | 0.88 | 0 | 0.69 | 0 |
| **PENS** | -0.02 | 0.557 | -0.19 | 0.567 |
| **PHCP** | 0.26 | 0.016 | -0.08 | 0.007 |
| **RSCZ_data** | 0.64 | 0 | 0.41 | 0 |
| **MCIC** | 0.92 | 0 | 0.68 | 0 |
| **ESO** | -0.55 | 1 | -0.42 | 1 |
| **CIAM** | 0.77 | 0 | 0.41 | 0 |
| **MPRC** | 0.6 | 0 | 0.42 | 0 |
| **OLIN** | 0.8 | 0 | 0.55 | 0 |

#### Subject-level cortical abnormality modeling

We next sought to examine whether our network-based models can be translated to individual schizophrenia patients' data and how they are influenced by individual clinical factors. Batch-corrected cortical thickness data of patients were first adjusted for age and sex by residualizing the effect of age and sex using a linear model. Subsequently they were z-scored relative to healthy controls to generate individualized morphological abnormality z-score maps. To test the hub vulnerability hypothesis at an individual level we descriptively compared the expected vs observed incidence of statistically significant positive correlations of nodal centrality and patient-specific morphological abnormality maps, adjusting for spatial autocorrelation using spin permutation tests. According to an alpha of 0.05 and the properties of the Gaussian distribution we define the expected incidence of statistically significant positive correlation values as 0.025. We next sought to test the robustness of our epicenter findings at the individual patient level. We identified patient-specific structural and functional epicenter maps by iteratively correlating the connectivity profile of each brain region to each patient's morphological abnormality map as described in the epicenter mapping section. Significance was tested for each patient-specific epicenters using spin permutation test ($p_{spin}<0.05$) as described in the method section in the manuscript. We finally computed the percentage of individuals for which each region was a statistically significant epicenter, correcting for multiple testing with the Bonferroni method.

*Subject-level hub vulnerability modeling*

To assess whether network atrophy models can also explain individual patient data, each patient-specific cortical abnormality map was correlated with the normative degree centrality maps (Fig. S2). We observed similar associations between individual cortical maps and functional (pspin < 0.05 in 18.2% of individuals with SCZ) as well as structural cortico-cortical hubs (pspin < 0.05 in 8.1% of individuals with SCZ) as seen in the group-level analysis. By contrast, a null distribution of p-values (corrected for spatial autocorrelation) would only show a rate of approximately 2,5% statistically significant positive correlations. Thus, we observed a 7.2-fold (18.2% vs. 2.5%; $\chi^2_{func}$ = 2466.4, df = 1, p-value < $2.2e^{-16}$) and 3.2-fold (8.1% vs 2.5%, $\chi^2_{struc}$= 315.58, df = 1, p-value < $2.2e^{-16}$) enrichment of significant associations between individual morphometric maps and cortical centrality maps than would be expected in the null hypothesis.

*Subject-level epicenter modeling*

Using each patient's individual cortical abnormality map, we further identified patient-specific structural and functional epicenters with a marked overlap in the top significant epicenters identified by our mega-analysis. Specifically, 9 out of 10 top epicenters overlap between the individual epicenter models and our group-level mega-analysis including the entorhinal cortices, banks of superior temporal sulci, left inferior temporal gyrus, and frontal gyri (bilateral pars triangularis, left pars opercularis, right pars orbitalis) (Fig. S2). In summary, although the individual subject-level data displayed overall lower sensitivity, due to the increased heterogeneity in cortical abnormality patterns, the results closely mirrored our group-level analysis. Collectively, individual network modeling supported both the hub vulnerability hypothesis and the most significant epicenters as identified by our mega-analysis.

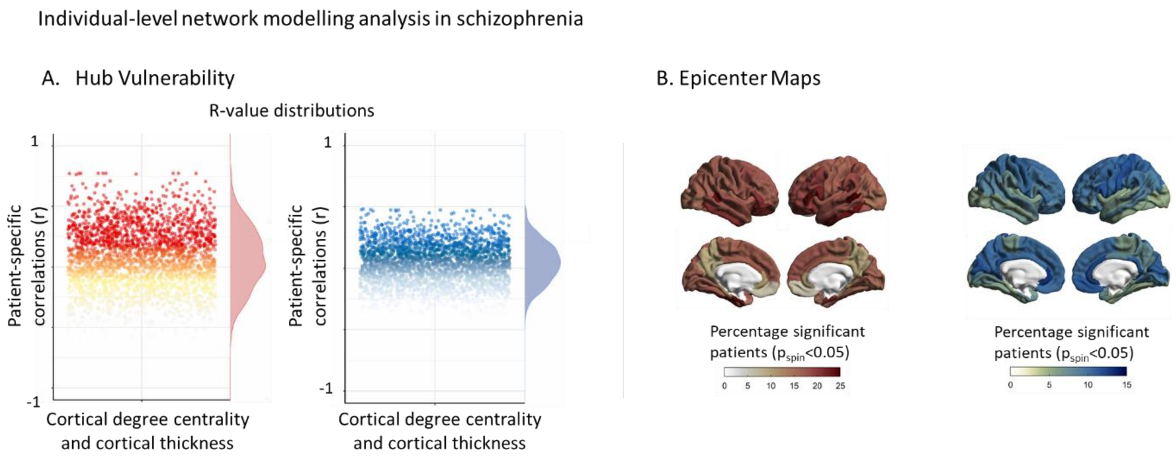


**Figure S2. Individual-level network modeling analysis in schizophrenia**. **(A)** Hub vulnerability. On an individual patient level, we computed patient-specific morphological abnormality maps and tested the hub vulnerability hypothesis for each patient correcting for spatial autocorrelation (pspin<0.05). The resulting R-value distributions are enriched in positive correlations (7.2-fold, 18.2% vs. 2.5%; $\chi^2_{func}$ = 2466.4, df = 1, p-value < 2.2e$^{-16}$ and 3.2-fold, 8.1% vs 2.5%, $\chi^2_{struc}$=315.58, df = 1, p-value < 2.2e$^{-16}$). **(B)** Epicenter Mapping. On an individual patient level, we computed patient-specific disease epicenters by identifying regions with a connectivity profile which significantly correlated ($p_{spin}$<0.05) with each patient's morphological abnormality map. Epicenter map depicts the percentage of individual patients for whom each region is a significant functional (above) or structural (below) epicenter. The ranking of regions most highly enriched for statistical significance in individuals correlates highly with the original epicenter map.

### Subject-level correlation of clinical variables to hub vulnerability and epicenters

Having established both network-based models in subject-level data (see results above), we next examined the association between individual clinical factors and patient's hub vulnerability and epicenters respectively. To this end, we correlated the subject-level hub vulnerability scores with antipsychotic medication, duration of illness, PANSS total, PANSS positive, negative and general score (Table S16). To examine the relationship between individual subject-level epicenters and clinical factors, we correlated each patient's epicenters (based on significance) with the above-mentioned clinical factors. To control for multiple comparisons p-values were adjusted within each clinical variable analysis using the Bonferroni method. (Tables S17-18).

**Table S16. Correlations between subject-level functional and structural hub vulnerability and clinical scores**

| **Individual Hub Vulnerability** | | | | |
|---|---|---|---|---|
| | **Functional Connectivity** | | **Structural Connectivity** | |
| Clinical Variables | R | pval (Bonferroni) | R | pval (Bonferroni) |
| PANSS Positive | 0.06 | 0.027 | 0.06 | 0.025 |
| PANSS Negative | 0.03 | 0.241 | 0.05 | 0.018 |
| PANSS General | 0.21 | <0.0001 | 0.13 | 0.01 |
| PANSS Total | 0.1 | 0.001 | 0.09 | 0.004 |
| Chlorpromazine | -0.02 | 0.403 | -0.01 | 0.832 |
| Duration of Illness | -0.04 | 0.112 | 0.02 | 0.309 |
| All p-values are corrected for multiple comparison using the Bonferroni method | | | | |

**Table S17. Correlations between individual subject-level functional epicenters and clinical scores**

| **Functional connectivity** | | | | | | | | | | | | |
|---|---|---|---|---|---|---|---|---|---|---|---|---|
| | PANSS Positive | | PANSS Negative | | PANSS General | | PANSS Total | | Chlorpromazine equivalents | | Duration of Illness | |
| **Brain Region** | R | pval | R | pval | R | pval | R | pval | R | pval | R | pval |
| Left banks of superior temporal sulcus | 0.05 | 1 | 0.03 | 1 | 0.15 | 0.296 | 0.09 | 0.296 | 0.02 | 1 | -0.04 | 1 |
| Left caudal anterior cingulate cortex | 0.05 | 1 | 0.04 | 1 | 0.24 | <0.001 | 0.1 | 0.088 | -0.03 | 1 | -0.03 | 1 |
| Left caudal middle frontal gyrus | 0.04 | 1 | 0.02 | 1 | 0.06 | 1 | 0.06 | 1 | 0.01 | 1 | -0.02 | 1 |
| Left cuneus | 0.04 | 1 | 0.01 | 1 | 0.22 | 0.001 | 0.08 | 0.704 | -0.03 | 1 | -0.03 | 1 |
| Left entorhinal cortex | 0.04 | 1 | 0.03 | 1 | 0.07 | 1 | 0.08 | 0.981 | 0 | 1 | -0.01 | 1 |
| Left fusiform gyrus | 0.05 | 1 | 0.01 | 1 | 0.17 | 0.052 | 0.08 | 0.763 | -0.03 | 1 | -0.02 | 1 |
| Left inferior parietal cortex | 0.04 | 1 | 0 | 1 | 0.06 | 1 | 0.05 | 1 | 0 | 1 | -0.04 | 1 |
| Left inferior temporal gyrus | 0.06 | 0.929 | 0.02 | 1 | 0.18 | 0.019 | 0.1 | 0.121 | 0.01 | 1 | -0.02 | 1 |
| Left isthmus cingulate cortex | 0.04 | 1 | 0.01 | 1 | 0.11 | 1 | 0.05 | 1 | -0.03 | 1 | -0.01 | 1 |
| Left lateral occipital cortex | 0.02 | 1 | -0.01 | 1 | 0.15 | 0.249 | 0.05 | 1 | -0.04 | 1 | -0.03 | 1 |
| Left lateral orbitofrontal cortex | 0.03 | 1 | 0.02 | 1 | 0.11 | 1 | 0.06 | 1 | 0.02 | 1 | -0.03 | 1 |
| Left lingual gyrus | 0.04 | 1 | 0.02 | 1 | 0.23 | <0.001 | 0.09 | 0.299 | -0.05 | 1 | -0.02 | 1 |
| Left medial orbitofrontal cortex | 0.01 | 1 | 0.01 | 1 | -0.01 | 1 | 0.02 | 1 | 0 | 1 | 0.02 | 1 |

| | | | | | | | | | | | | |
|---|---|---|---|---|---|---|---|---|---|---|---|---|
| Left middle temporal gyrus | 0.05 | 1 | 0.02 | 1 | 0.02 | 1 | 0.05 | 1 | 0.04 | 1 | -0.01 | 1 |
| Left parahippocampal gyrus | 0.04 | 1 | 0.03 | 1 | 0.13 | 0.957 | 0.07 | 1 | -0.03 | 1 | 0 | 1 |
| Left paracentral lobule | 0.08 | 0.088 | 0.05 | 1 | 0.24 | <0.001 | 0.13 | 0.002 | -0.02 | 1 | -0.04 | 1 |
| Left pars opercularis of inferior frontal gyrus | 0.04 | 1 | 0.01 | 1 | 0.15 | 0.284 | 0.07 | 1 | 0.01 | 1 | -0.05 | 0.979 |
| Left pars orbitalis of inferior frontal gyrus | -0.01 | 1 | 0 | 1 | -0.05 | 1 | 0 | 1 | 0.03 | 1 | -0.04 | 1 |
| Left pars triangularis of inferior frontal gyrus | 0.04 | 1 | 0.02 | 1 | 0.09 | 1 | 0.06 | 1 | 0.06 | 1 | -0.03 | 1 |
| Left pericalcarine cortex | 0.03 | 1 | 0.02 | 1 | 0.22 | 0.001 | 0.07 | 1 | -0.06 | 1 | -0.01 | 1 |
| Left postcentral gyrus | 0.09 | 0.022 | 0.05 | 1 | 0.26 | <0.001 | 0.14 | <0.001 | -0.01 | 1 | -0.03 | 1 |
| Left posterior cingulate cortex | 0.04 | 1 | 0.03 | 1 | 0.2 | 0.005 | 0.09 | 0.308 | -0.03 | 1 | -0.03 | 1 |
| Left precentral gyrus | 0.08 | 0.143 | 0.05 | 1 | 0.24 | <0.001 | 0.13 | 0.002 | -0.01 | 1 | -0.04 | 1 |
| Left precuneus | 0.05 | 1 | 0.02 | 1 | 0.18 | 0.025 | 0.09 | 0.204 | -0.05 | 1 | -0.01 | 1 |
| Left rostral anterior cingulate cortex | 0.03 | 1 | 0.04 | 1 | 0.11 | 1 | 0.06 | 1 | -0.04 | 1 | -0.01 | 1 |
| Left rostral middle frontal gyrus | 0.04 | 1 | 0.02 | 1 | 0.16 | 0.126 | 0.08 | 0.691 | 0 | 1 | -0.02 | 1 |
| Left superior frontal gyrus | 0.05 | 1 | 0.02 | 1 | 0.14 | 0.342 | 0.08 | 0.791 | 0 | 1 | -0.03 | 1 |
| Left superior parietal cortex | 0.05 | 1 | 0.03 | 1 | 0.23 | 0.001 | 0.1 | 0.073 | -0.04 | 1 | -0.02 | 1 |

| | | | | | | | | | | | | |
|---|---|---|---|---|---|---|---|---|---|---|---|---|
| Left superior temporal gyrus | 0.05 | 1 | 0.03 | 1 | 0.18 | 0.035 | 0.1 | 0.103 | -0.01 | 1 | -0.05 | 1 |
| Left supramarginal gyrus | 0.05 | 1 | 0.02 | 1 | 0.13 | 0.68 | 0.07 | 1 | -0.01 | 1 | -0.04 | 1 |
| Left frontal pole | 0 | 1 | 0.01 | 1 | -0.08 | 1 | 0 | 1 | 0.04 | 1 | 0 | 1 |
| Left temporal pole | -0.01 | 1 | -0.01 | 1 | -0.14 | 0.442 | -0.03 | 1 | 0.07 | 1 | -0.02 | 1 |
| Left transverse temporal gyrus | 0.07 | 0.56 | 0.04 | 1 | 0.24 | <0.001 | 0.11 | 0.014 | -0.01 | 1 | -0.02 | 1 |
| Left insula | 0.05 | 1 | 0.03 | 1 | 0.23 | <0.001 | 0.1 | 0.059 | -0.02 | 1 | -0.04 | 1 |
| Right banks of superior temporal sulcus | 0.04 | 1 | 0.03 | 1 | 0.16 | 0.081 | 0.09 | 0.265 | 0.01 | 1 | -0.05 | 1 |
| Right caudal anterior cingulate cortex | 0.05 | 1 | 0.03 | 1 | 0.21 | 0.003 | 0.1 | 0.071 | -0.03 | 1 | -0.03 | 1 |
| Right caudal middle frontal gyrus | 0.01 | 1 | -0.02 | 1 | 0.05 | 1 | 0.03 | 1 | -0.02 | 1 | -0.05 | 1 |
| Right cuneus | 0.05 | 1 | 0.03 | 1 | 0.24 | <0.001 | 0.1 | 0.057 | -0.06 | 1 | -0.02 | 1 |
| Right entorhinal cortex | 0.04 | 1 | 0.01 | 1 | 0.03 | 1 | 0.05 | 1 | 0.02 | 1 | -0.02 | 1 |
| Right fusiform gyrus | 0.04 | 1 | 0.02 | 1 | 0.15 | 0.191 | 0.07 | 1 | -0.04 | 1 | -0.01 | 1 |
| Right inferior parietal cortex | 0.04 | 1 | 0.01 | 1 | 0.13 | 0.895 | 0.07 | 1 | -0.03 | 1 | -0.03 | 1 |
| Right inferior temporal gyrus | 0.03 | 1 | 0 | 1 | 0.08 | 1 | 0.05 | 1 | -0.01 | 1 | -0.04 | 1 |
| Right isthmus cingulate cortex | 0.03 | 1 | 0.01 | 1 | 0.13 | 0.682 | 0.06 | 1 | -0.07 | 1 | -0.02 | 1 |
| Right lateral occipital cortex | 0.04 | 1 | 0.03 | 1 | 0.18 | 0.025 | 0.08 | 0.753 | -0.03 | 1 | -0.01 | 1 |
| Right lateral orbitofrontal cortex | 0.02 | 1 | -0.01 | 1 | 0.1 | 1 | 0.03 | 1 | 0.01 | 1 | -0.05 | 1 |
| Right lingual gyrus | 0.04 | 1 | 0.01 | 1 | 0.21 | 0.002 | 0.07 | 1 | -0.05 | 1 | -0.02 | 1 |

| | | | | | | | | | | | | |
|---|---|---|---|---|---|---|---|---|---|---|---|---|
| Right medial orbitofrontal cortex | -0.01 | 1 | 0.01 | 1 | -0.06 | 1 | 0 | 1 | 0 | 1 | 0.01 | 1 |
| Right middle temporal gyrus | 0.03 | 1 | 0 | 1 | 0.06 | 1 | 0.04 | 1 | 0.02 | 1 | -0.01 | 1 |
| Right parahippocampal gyrus | 0.03 | 1 | 0.02 | 1 | 0.14 | 0.42 | 0.08 | 0.832 | -0.04 | 1 | -0.01 | 1 |
| Right paracentral lobule | 0.06 | 0.724 | 0.04 | 1 | 0.24 | <0.001 | 0.12 | 0.01 | -0.02 | 1 | -0.03 | 1 |
| Right pars opercularis of inferior frontal gyrus | 0.04 | 1 | 0.03 | 1 | 0.17 | 0.044 | 0.09 | 0.378 | -0.01 | 1 | -0.03 | 1 |
| Right pars orbitalis of inferior frontal gyrus | 0.01 | 1 | 0.01 | 1 | -0.04 | 1 | 0.01 | 1 | 0.04 | 1 | -0.01 | 1 |
| Right pars triangularis of inferior frontal gyrus | 0.02 | 1 | 0 | 1 | 0.12 | 1 | 0.04 | 1 | 0.01 | 1 | -0.06 | 0.443 |
| Right pericalcarine cortex | 0.04 | 1 | 0 | 1 | 0.22 | 0.001 | 0.07 | 1 | -0.05 | 1 | -0.03 | 1 |
| Right postcentral gyrus | 0.09 | 0.041 | 0.06 | 0.877 | 0.24 | <0.001 | 0.14 | <0.001 | -0.01 | 1 | -0.03 | 1 |
| Right posterior cingulate cortex | 0.04 | 1 | 0.03 | 1 | 0.21 | 0.002 | 0.09 | 0.306 | -0.03 | 1 | -0.03 | 1 |
| Right precentral gyrus | 0.06 | 1 | 0.03 | 1 | 0.21 | 0.001 | 0.1 | 0.064 | -0.01 | 1 | -0.03 | 1 |
| Right precuneus | 0.04 | 1 | 0.03 | 1 | 0.19 | 0.011 | 0.09 | 0.325 | -0.06 | 1 | 0 | 1 |
| Right rostral anterior cingulate cortex | -0.03 | 1 | 0.02 | 1 | 0.06 | 1 | 0.01 | 1 | -0.03 | 1 | 0.01 | 1 |
| Right rostral middle frontal gyrus | 0.03 | 1 | 0.01 | 1 | 0.14 | 0.303 | 0.05 | 1 | -0.02 | 1 | -0.04 | 1 |

| | | | | | | | | | | | | |
|---|---|---|---|---|---|---|---|---|---|---|---|---|
| Right superior frontal gyrus | 0.04 | 1 | 0.02 | 1 | 0.16 | 0.088 | 0.08 | 0.455 | -0.02 | 1 | -0.05 | 1 |
| Right superior parietal cortex | 0.05 | 1 | 0.02 | 1 | 0.18 | 0.02 | 0.08 | 0.762 | -0.03 | 1 | -0.02 | 1 |
| Right superior temporal gyrus | 0.05 | 1 | 0.03 | 1 | 0.17 | 0.057 | 0.09 | 0.34 | -0.01 | 1 | -0.04 | 1 |
| Right supramarginal gyrus | 0.07 | 0.57 | 0.03 | 1 | 0.23 | <0.001 | 0.11 | 0.015 | -0.01 | 1 | -0.03 | 1 |
| Right frontal pole | -0.01 | 1 | 0.01 | 1 | -0.08 | 1 | -0.02 | 1 | 0.02 | 1 | -0.01 | 1 |
| Right temporal pole | -0.01 | 1 | 0.01 | 1 | -0.13 | 0.83 | -0.01 | 1 | 0.03 | 1 | -0.01 | 1 |
| Right transverse temporal gyrus | 0.06 | 0.82 | 0.05 | 1 | 0.26 | <0.001 | 0.12 | 0.008 | -0.02 | 1 | -0.04 | 1 |
| Right insula | 0.05 | 1 | 0.04 | 1 | 0.24 | <0.001 | 0.11 | 0.033 | -0.02 | 1 | -0.04 | 1 |
| All p-values are corrected for multiple comparison using the Bonferroni method | | | | | | | | | | | | |

**Table S18. Correlations between individual subject-level structural epicenters and clinical scores**

| **Structural Connectivity** | | | | | | | | | | | | |
|---|---|---|---|---|---|---|---|---|---|---|---|---|
| | PANSS Positive | | PANSS Negative | | PANSS General | | PANSS Total | | Chlorpromazine equivalents | | Duration of Illness | |
| | R | pval | R | pval | R | pval | R | pval | R | pval | R | pval |
| Left banks of superior temporal sulcus | -0.01 | 1 | 0.01 | 1 | -0.09 | 1 | 0 | 1 | 0.02 | 1 | 0.01 | 1 |
| Left caudal anterior cingulate cortex | -0.01 | 1 | 0.02 | 1 | 0.04 | 1 | -0.01 | 1 | -0.01 | 1 | 0.02 | 1 |
| Left caudal middle frontal gyrus | 0.08 | 0.228 | 0.07 | 0.009 | 0.1 | 0.265 | 0.11 | 0.001 | 0.02 | 1 | 0 | |
| Left cuneus | 0.01 | 1 | -0.01 | 1 | 0.11 | 1 | 0.03 | 1 | -0.06 | 1 | 0.01 | 1 |
| Left entorhinal cortex | -0.01 | 1 | 0.01 | 1 | -0.12 | 1 | -0.01 | 1 | -0.04 | 1 | 0.03 | 1 |
| Left fusiform gyrus | 0 | 1 | -0.02 | 1 | -0.08 | 1 | -0.02 | 1 | -0.04 | 1 | 0.03 | 0.415 |
| Left inferior parietal cortex | 0.03 | 1 | 0.03 | 1 | -0.04 | 1 | 0.04 | 1 | 0.02 | 1 | 0.01 | 1 |
| Left inferior temporal gyrus | 0.01 | 1 | 0.01 | 1 | -0.03 | 1 | 0.02 | 1 | -0.01 | 1 | 0.02 | 1 |
| Left isthmus cingulate cortex | 0.02 | 1 | -0.02 | 1 | 0.11 | 1 | 0.02 | 1 | -0.05 | 1 | 0.03 | 1 |
| Left lateral occipital cortex | 0.03 | 0.065 | -0.02 | 1 | -0.11 | 1 | -0.03 | 0.143 | -0.05 | 1 | 0.03 | 1 |
| Left lateral orbitofrontal cortex | 0.02 | 1 | 0.03 | 1 | -0.04 | 1 | -0.02 | 1 | 0.04 | 1 | 0.04 | 1 |
| Left lingual gyrus | 0.02 | 1 | -0.02 | 1 | -0.04 | 1 | -0.03 | 1 | -0.05 | 1 | 0.01 | 1 |
| Left medial orbitofrontal cortex | 0.05 | 1 | 0 | 1 | -0.13 | 1 | -0.06 | 1 | 0.03 | 1 | 0.04 | 1 |
| Left middle temporal gyrus | 0.02 | 1 | 0.01 | 0.508 | -0.07 | 1 | 0.01 | 1 | 0.04 | 1 | 0.03 | 1 |
| Left parahippocampal gyrus | 0.02 | 1 | -0.01 | 1 | -0.08 | 1 | -0.03 | 1 | -0.05 | 1 | 0.02 | 1 |
| Left paracentral lobule | 0.12 | 1 | 0.07 | 1 | 0.24 | 1 | 0.16 | 1 | -0.02 | 1 | 0 | 1 |
| Left pars opercularis of inferior frontal gyrus | 0.04 | 1 | 0.03 | 1 | 0.04 | 1 | 0.05 | 1 | 0.05 | 1 | 0 | 1 |
| Left pars orbitalis of inferior frontal gyrus | 0.01 | 1 | 0.01 | 0.386 | -0.01 | 1 | 0 | 1 | 0.05 | 1 | -0.01 | 1 |

| | | | | | | | | | | | | |
|---|---|---|---|---|---|---|---|---|---|---|---|---|
| Left pars triangularis of inferior frontal gyrus | 0.02 | 1 | 0.03 | 1 | -0.04 | 1 | 0.02 | 1 | 0.08 | 1 | 0.01 | 1 |
| Left pericalcarine cortex | 0.01 | 1 | -0.01 | 1 | 0.01 | 1 | -0.01 | 1 | -0.06 | 1 | 0.02 | 0.134 |
| Left postcentral gyrus | 0.11 | 1 | 0.06 | 1 | 0.21 | 1 | 0.15 | 1 | 0 | 1 | -0.02 | 0.154 |
| Left posterior cingulate cortex | 0.03 | 1 | 0.04 | 1 | 0.11 | 1 | 0.06 | 1 | 0.01 | 1 | 0.05 | 1 |
| Left precentral gyrus | 0.15 | 1 | 0.08 | 1 | 0.23 | 0.064 | 0.17 | 0.357 | 0.04 | 1 | -0.01 | 1 |
| Left precuneus | 0.06 | 1 | 0.02 | 1 | 0.17 | 1 | 0.08 | 1 | -0.04 | 1 | 0.01 | 1 |
| Left rostral anterior cingulate cortex | 0.01 | 1 | 0.03 | 1 | -0.01 | 1 | -0.01 | 1 | 0.01 | 0.533 | 0.02 | 1 |
| Left rostral middle frontal gyrus | 0.01 | 1 | 0.02 | 1 | -0.01 | 1 | 0.01 | 1 | 0.04 | 1 | 0.01 | 1 |
| Left superior frontal gyrus | 0.06 | 1 | 0.04 | 1 | 0.13 | 0.659 | 0.06 | 1 | 0.01 | 1 | 0.03 | 1 |
| Left superior parietal cortex | 0.05 | 1 | 0.04 | 1 | 0.23 | 1 | 0.11 | 1 | -0.07 | 1 | 0 | 1 |
| Left superior temporal gyrus | 0.02 | 1 | 0.01 | 1 | -0.06 | 1 | -0.01 | 1 | 0.02 | 1 | 0 | 1 |
| Left supramarginal gyrus | 0.02 | 1 | 0.01 | 1 | -0.01 | 1 | 0.03 | 1 | 0.04 | 1 | -0.03 | 1 |
| Left frontal pole | 0.01 | 1 | 0.02 | 1 | -0.01 | 1 | -0.01 | 1 | 0.02 | 1 | 0.01 | 1 |
| Left temporal pole | 0.08 | 0.118 | -0.02 | 1 | -0.29 | 0.065 | -0.12 | 0.346 | 0.05 | 1 | 0.02 | 1 |
| Left transverse temporal gyrus | 0.04 | 1 | 0.04 | 1 | 0.05 | 1 | 0.06 | 1 | 0.03 | 1 | 0 | 1 |
| Left insula | 0 | 1 | 0.03 | 1 | -0.04 | 1 | 0 | 1 | 0 | 1 | 0.02 | 1 |
| Right banks of superior temporal sulcus | 0.04 | 1 | -0.01 | 1 | -0.15 | 1 | -0.05 | 1 | 0.01 | 1 | -0.03 | 1 |
| Right caudal anterior cingulate cortex | 0.01 | 1 | 0.02 | 1 | 0.02 | 1 | 0.01 | 1 | 0.01 | 1 | 0.02 | 1 |
| Right caudal middle frontal gyrus | 0.07 | 1 | 0.05 | 1 | 0.19 | 0.166 | 0.11 | 1 | 0.01 | 1 | -0.02 | 0.185 |
| Right cuneus | 0.01 | 1 | -0.01 | 1 | 0.09 | 1 | 0.03 | 1 | -0.06 | 1 | 0.01 | 1 |
| Right entorhinal cortex | 0 | 1 | -0.01 | 1 | -0.08 | 1 | -0.02 | 1 | 0.03 | 1 | 0.03 | 1 |
| Right fusiform gyrus | 0.01 | 1 | -0.02 | 1 | -0.04 | 1 | -0.02 | 1 | -0.01 | 1 | 0.02 | 1 |
| Right inferior parietal cortex | 0.02 | 1 | -0.01 | 1 | 0.02 | 1 | 0.01 | 1 | 0.01 | 1 | -0.04 | 1 |

| | | | | | | | | | | | | |
|---|---|---|---|---|---|---|---|---|---|---|---|---|
| Right inferior temporal gyrus | 0.04 | 1 | -0.03 | 1 | -0.16 | 1 | -0.08 | 1 | 0.03 | 1 | 0 | 1 |
| Right isthmus cingulate cortex | 0 | 1 | 0 | 1 | 0.1 | 1 | 0.02 | 1 | -0.08 | 1 | 0.03 | 1 |
| Right lateral occipital cortex | 0.02 | 1 | -0.02 | 1 | -0.01 | 1 | -0.02 | 1 | -0.01 | 1 | 0.01 | 1 |
| Right lateral orbitofrontal cortex | 0.05 | 1 | -0.02 | 1 | -0.08 | 1 | -0.07 | 1 | 0.03 | 1 | -0.01 | 1 |
| Right lingual gyrus | 0.02 | 1 | -0.02 | 1 | -0.01 | 0.558 | -0.02 | 1 | -0.04 | 1 | 0.03 | 1 |
| Right medial orbitofrontal cortex | 0.03 | 1 | 0 | 1 | -0.05 | 1 | -0.04 | 1 | 0.02 | 0.164 | 0.02 | 1 |
| Right middle temporal gyrus | 0.03 | 1 | -0.01 | 1 | -0.03 | 1 | -0.03 | 1 | 0.02 | 1 | 0 | 1 |
| Right parahippocampal gyrus | 0.04 | 1 | -0.01 | 1 | -0.04 | 1 | -0.03 | 1 | -0.01 | 1 | 0.03 | 0.759 |
| Right paracentral lobule | 0.1 | 1 | 0.07 | 1 | 0.26 | 1 | 0.16 | 1 | -0.03 | 1 | 0 | 1 |
| Right pars opercularis of inferior frontal gyrus | 0.05 | 0.865 | 0.04 | 1 | 0.17 | 1 | 0.08 | 0.958 | 0 | 1 | -0.03 | 1 |
| Right pars orbitalis of inferior frontal gyrus | 0.05 | 1 | -0.02 | 1 | -0.13 | 1 | -0.06 | 1 | -0.01 | 1 | -0.02 | 1 |
| Right pars triangularis of inferior frontal gyrus | 0.01 | 1 | 0.01 | 1 | 0.04 | 1 | 0.02 | 1 | 0.01 | 1 | -0.06 | 1 |
| Right pericalcarine cortex | 0.01 | 1 | -0.01 | 1 | 0.1 | 1 | 0.01 | 1 | -0.06 | 1 | 0 | 1 |
| Right postcentral gyrus | 0.09 | 0.502 | 0.05 | 1 | 0.16 | 0.735 | 0.11 | 1 | -0.01 | 1 | -0.01 | 1 |
| Right posterior cingulate cortex | 0.08 | 1 | 0.06 | 1 | 0.23 | 1 | 0.12 | 1 | -0.01 | 1 | 0.02 | 1 |
| Right precentral gyrus | 0.1 | 0.504 | 0.05 | 1 | 0.22 | 0.066 | 0.13 | 1 | 0 | 1 | -0.03 | 0.574 |
| Right precuneus | 0.04 | 1 | 0.03 | 1 | 0.16 | 1 | 0.08 | 1 | -0.04 | 1 | 0.03 | 1 |
| Right rostral anterior cingulate cortex | 0.05 | 1 | 0 | 1 | -0.04 | 1 | -0.05 | 1 | 0.02 | 1 | 0.02 | 1 |
| Right rostral middle frontal gyrus | 0 | 1 | 0 | 1 | 0.05 | 1 | -0.01 | 1 | 0.02 | 1 | -0.03 | 1 |
| Right superior frontal gyrus | 0.06 | 1 | 0.04 | 1 | 0.14 | 1 | 0.07 | 1 | 0.01 | 1 | 0.02 | 1 |
| Right superior parietal cortex | 0.03 | 1 | 0.01 | 1 | 0.13 | 1 | 0.06 | 1 | -0.05 | 1 | -0.02 | 1 |

| | | | | | | | | | | | | |
|---|---|---|---|---|---|---|---|---|---|---|---|---|
| Right superior temporal gyrus | 0.08 | 1 | -0.05 | 1 | -0.22 | 1 | -0.11 | 1 | 0.02 | 1 | -0.02 | 1 |
| Right supramarginal gyrus | 0 | 1 | -0.02 | 1 | -0.02 | 1 | 0 | 1 | 0.01 | 1 | -0.05 | 1 |
| Right frontal pole | 0.01 | 1 | 0.02 | 1 | 0.03 | 1 | 0 | 1 | 0.03 | 1 | 0 | 1 |
| Right temporal pole | 0.04 | 1 | 0 | 1 | -0.17 | 1 | -0.06 | 1 | 0.01 | 1 | 0.02 | 1 |
| Right transverse temporal gyrus | 0 | 1 | 0.01 | 1 | -0.04 | 1 | 0 | 1 | 0.02 | 1 | -0.01 | 1 |
| Right insula | 0.03 | 1 | 0.01 | 1 | 0.01 | 1 | -0.01 | 1 | 0.01 | 1 | -0.02 | 1 |
| All p-values are corrected for multiple comparison using the Bonferroni method | | | | | | | | | | | | |

*Subject-level robustness and sensitivity analysis*

To show the robustness of the correlation values between clinical variables and our individual-level hub vulnerability and epicenter findings we performed a sensitivity permutation analysis in our sample. Specifically, we generated 100 different permutations of 80% of our sample- i.e., an 80-20 split without resampling. We accordingly repeated the correlation analysis with the clinical variables 100 times. Our results show that the original findings are highly robust to perturbation of our sample (Fig S3-9).

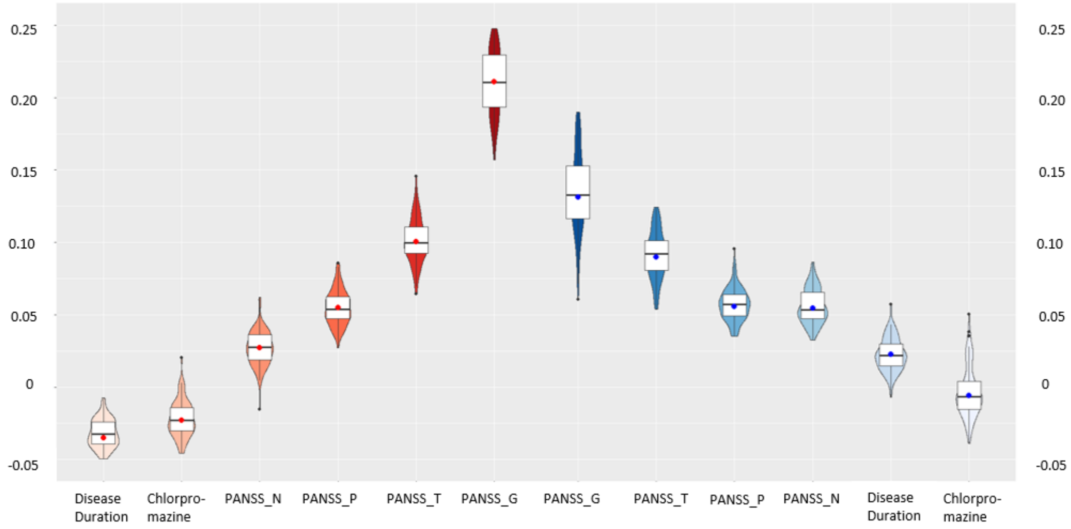


Figure S3. Robustness-Permutation analysis of stability of correlation of hub vulnerability at the individual level to individual clinical symptoms. Violin plots represent the distribution of values of the 100 permutations using a different 80% of the sample. Central dot represents the value obtained using 100% of our sample.

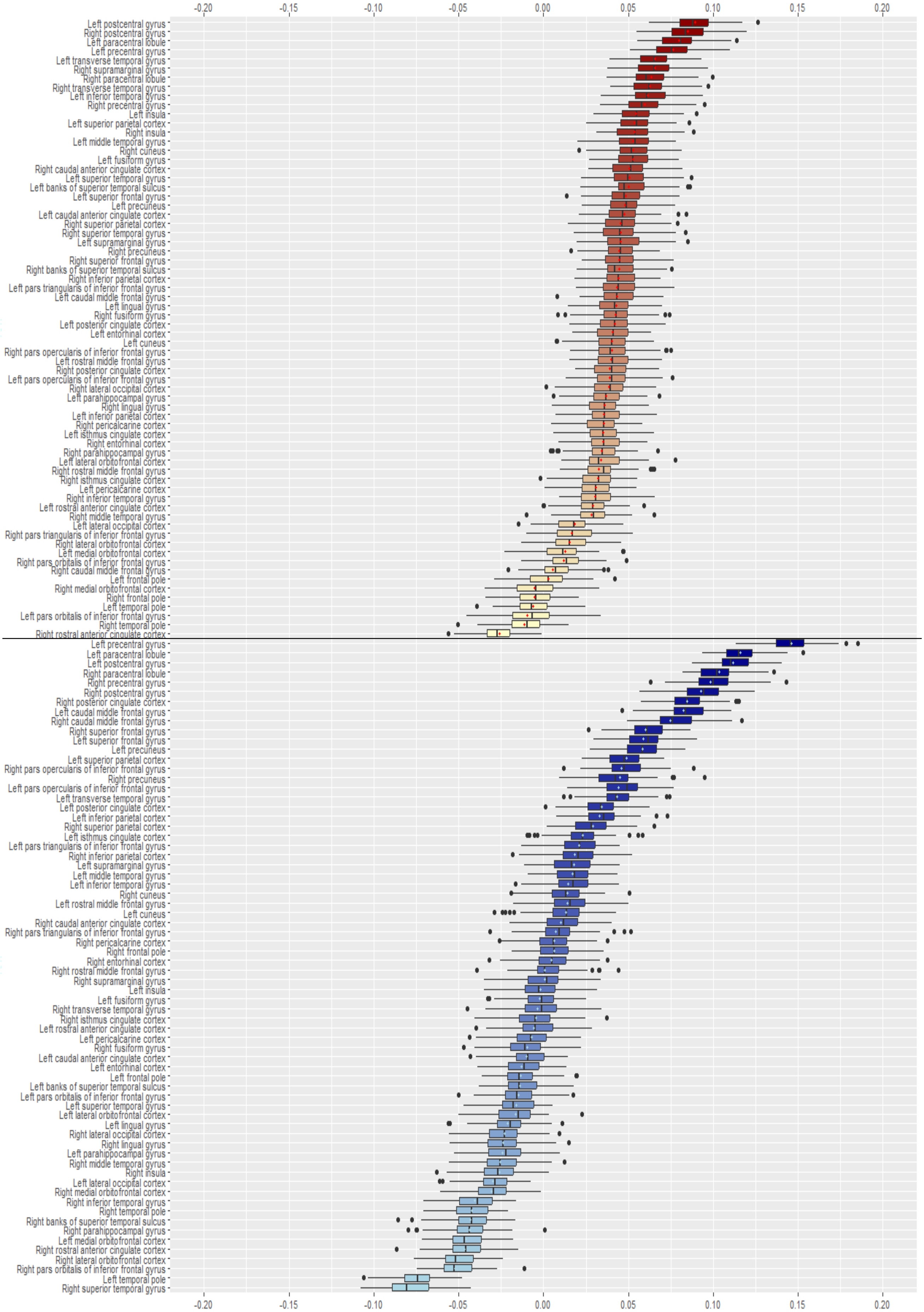


Figure S4. Robustness-Permutation analysis of stability of correlation of epicenter values at the individual level to individual PANSS Positive symptoms. Boxplots represent the distribution of values of the 100 permutations using a different 80% of the sample. Central dot represents the value obtained using 100% of our sample.

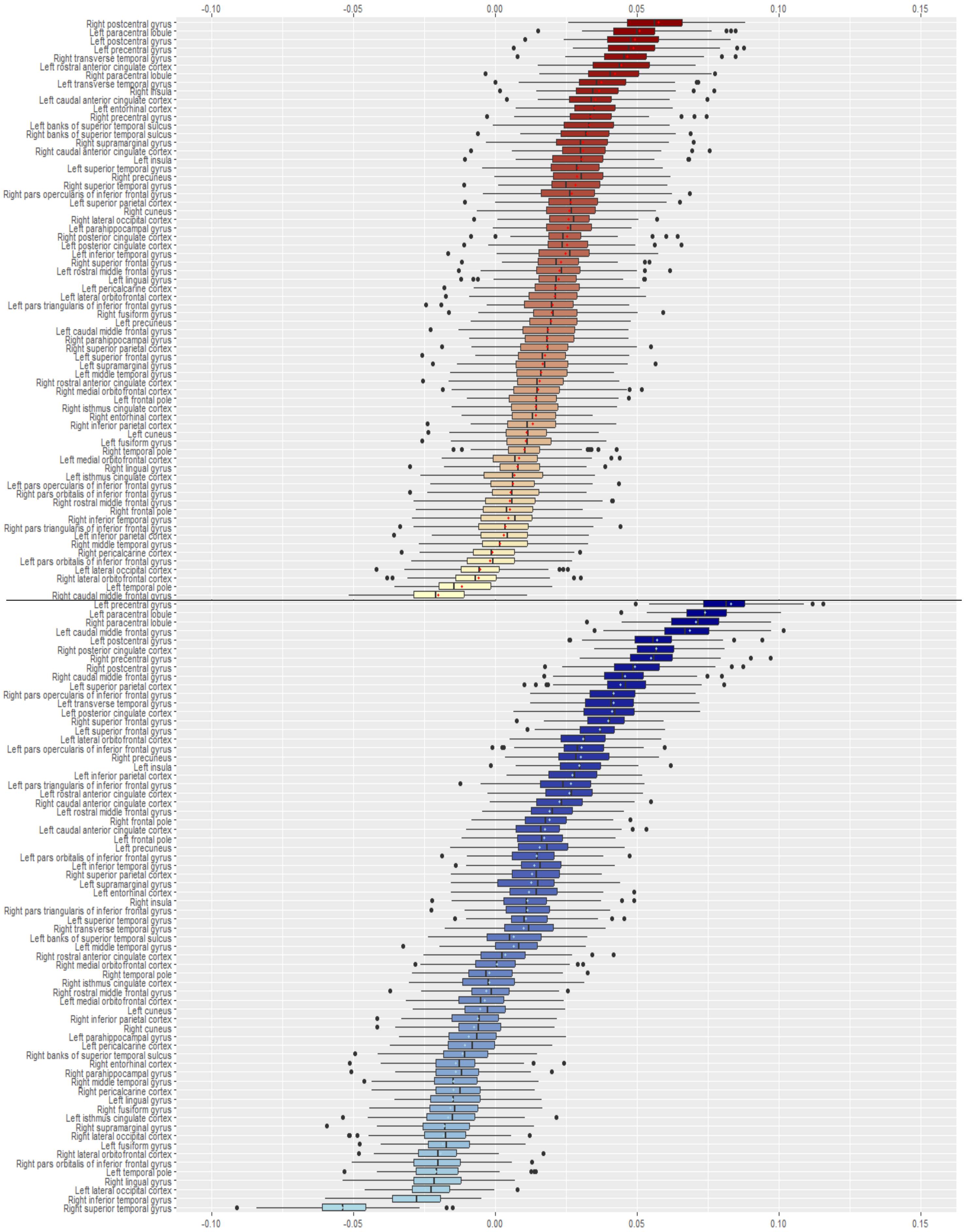


Figure S5. Robustness-Permutation analysis of stability of correlation of epicenter values at the individual level to individual PANSS Negative symptoms. Boxplots represent the distribution of values of the 100 permutations using a different 80% of the sample. Central dot represents the value obtained using 100% of our sample.

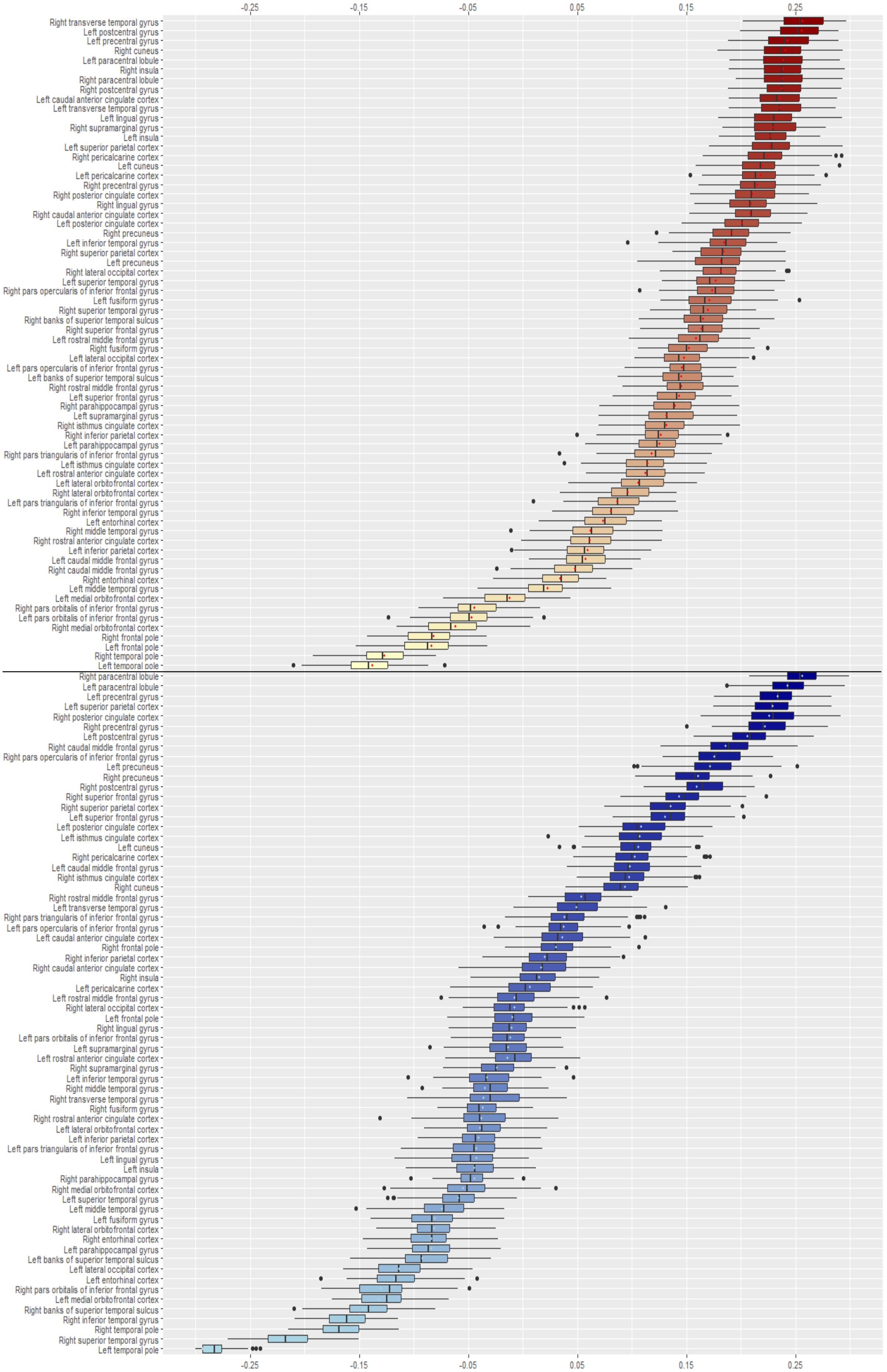


Figure S6. Robustness-Permutation analysis of stability of correlation of epicenter values at the individual level to individual PANSS General symptoms. Boxplots represent the distribution of values of the 100 permutations using a different 80% of the sample. Central dot represents the value obtained using 100% of our sample.

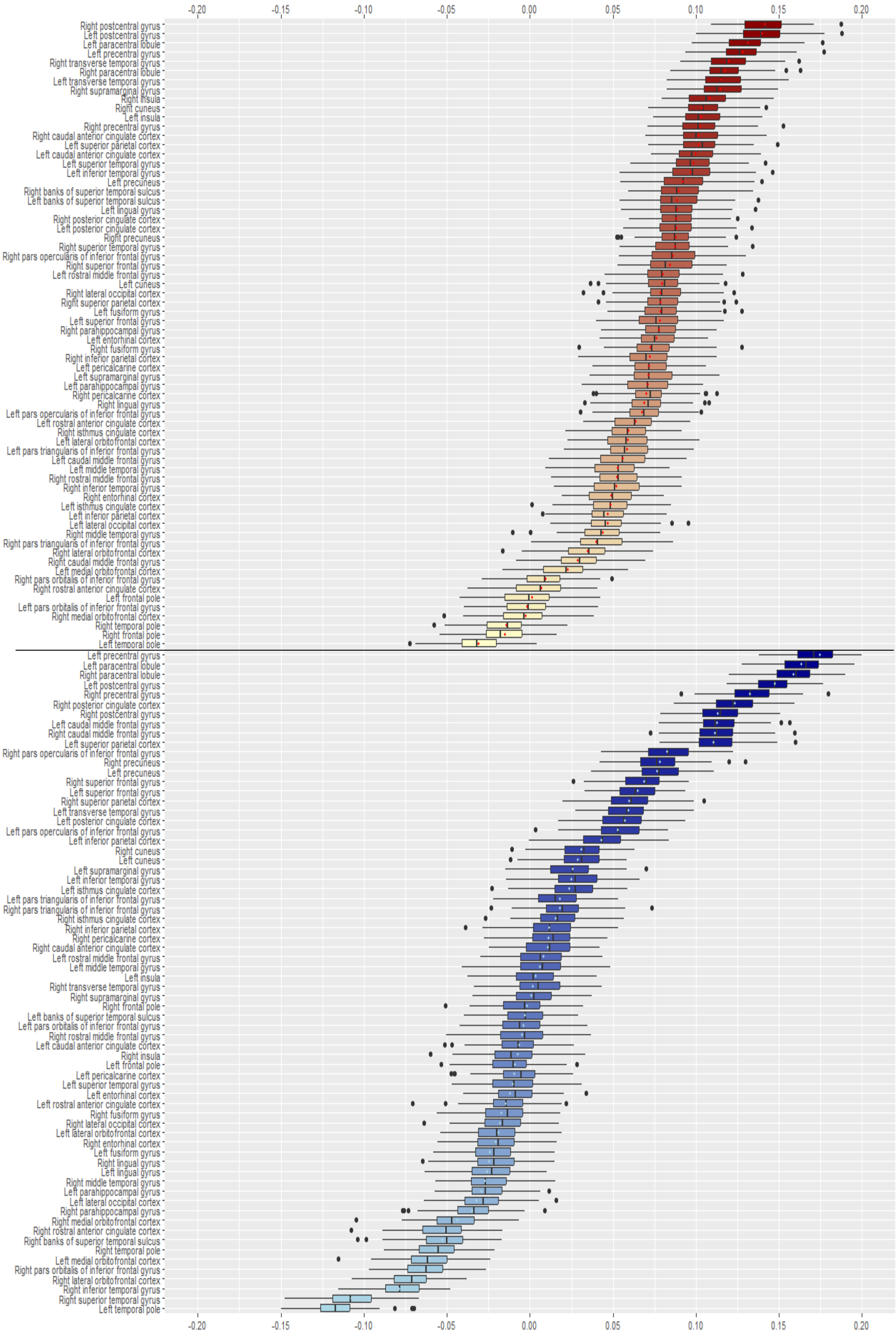


Figure S7. Robustness-Permutation analysis of stability of correlation of epicenter values at the individual level to individual PANSS Total Score. Boxplots represent the distribution of values of the 100 permutations using a different 80% of the sample. Central dot represents the value obtained using 100% of our sample.

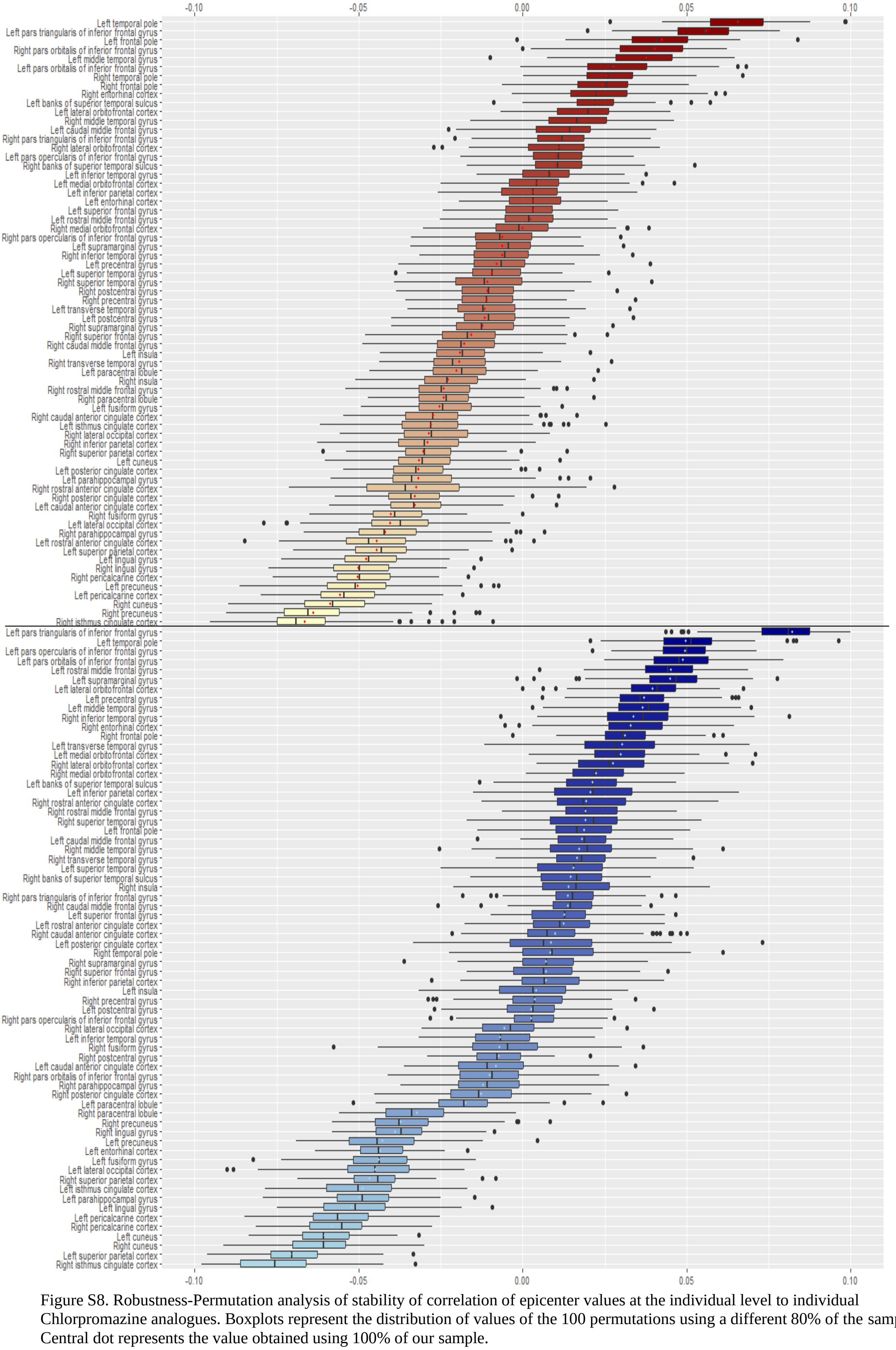

Figure S8. Robustness-Permutation analysis of stability of correlation of epicenter values at the individual level to individual Chlorpromazine analogues. Boxplots represent the distribution of values of the 100 permutations using a different 80% of the sample. Central dot represents the value obtained using 100% of our sample.

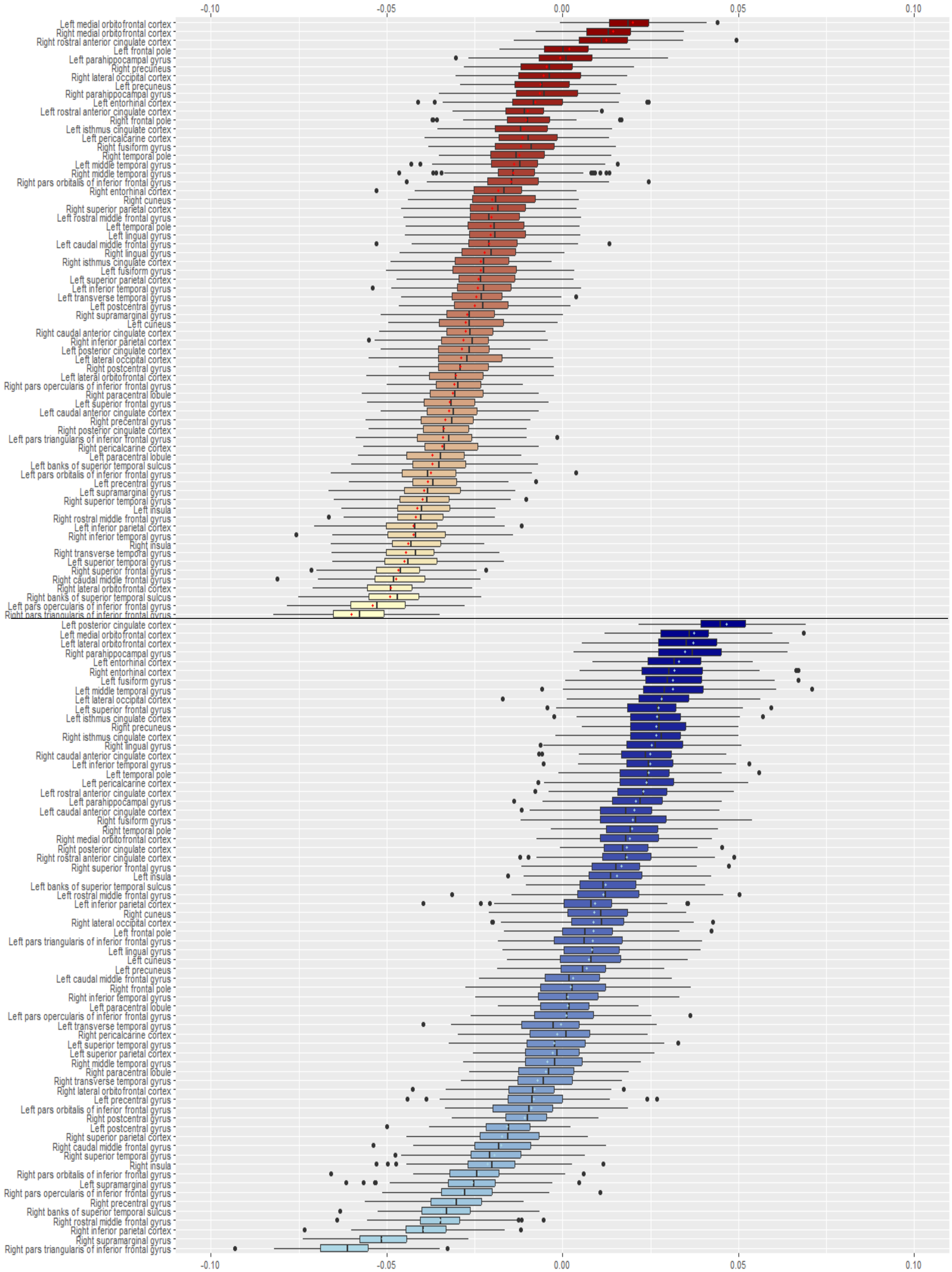


Figure S9. Robustness-Permutation analysis of stability of correlation of epicenter values at the individual level to individual Duration of Illness. Boxplots represent the distribution of values of the 100 permutations using a different 80% of the sample. Central dot represents the value obtained using 100% of our sample.

### Acknowledgments per dataset

Acknowledgments per dataset are as follows:

**ASRB:** The Australian Schizophrenia Research Bank (ASRB), was supported by the National Health and Medical Research Council of Australia (NHMRC) (Enabling Grant, ID 386500), the Pratt Foundation, Ramsay Health Care, the Viertel Charitable Foundation and the Schizophrenia Research Institute. Chief Investigators for ASRB were Carr, V., Schall, U., Scott, R., Jablensky, A., Mowry, B., Michie, P., Catts, S., Henskens, F., Pantelis, C. We thank Loughland, C., the ASRB Manager, and acknowledge the help of Jason Bridge with ASRB database queries. CP was supported by NHMRC Senior Principal Research Fellowships (IDs: 628386 & 1105825); GC was supported by the Schizophrenia Research Institute utilizing infrastructure funding from the New South Wales Ministry of Health and New South Wales Ministry of Trade and Investment (Australia); JF was supported by NHMRC project grant (1063960); MG was supported by NHMRC as an R.D. Wright Biomedical Career Development Fellow (1061875). MJC was supported by NHMRC Senior Research Fellowship (1121474). CSW is funded by the NSW Ministry of Health, Office of Health and Medical Research. CSW is a recipient of a National Health and Medical Research Council (Australia) Principal Research Fellowship (PRF) (#1117079).

**CAMH:** The CAMH datasets were collected and shared with support from the CAMH Foundation and the Canadian Institutes of Health Research.

**CIAM:** The CIAM study is supported by the South African Medical Research Council and National Research Foundation of South Africa.

**COBRE:** The COBRE dataset and investigators were supported by NIH grants R01EB006841 & P20GM103472, as well as NSF grant 1539067. JT (senior author) and VDC are supported by 5R01MH094524. JMS is supported by R01 AA021771 and P50 AA022534.

**ESO:** The ESO study was funded by NPU I – LO1611 and Ministry of Health, Czech Republic – Conceptual Development of Research Organization 00023001 (IKEM).

**FIDMAG:** Supported by Instituto de Salud Carlos III (Co-funded by European Regional Development Fund/European Social Fund) "Investing in your future"): Miguel Servet Research Contract (CPII16/00018 to E. Pomarol-Clotet and CP14/00041 to J. Radua.).

**FOR2107 Marburg:** This work was funded by the German Research Foundation (DFG), Tilo Kircher (speaker FOR2107; DFG grant numbers KI 588/14-1, KI 588/14-2), Axel Krug (KR 3822/5-1, KR 3822/7-2), Igor Nenadic (NE 2254/1-2), Carsten Konrad (KO 4291/3-1).

**FOR2107 Muenster:** The FOR2107 Muenster study was funded by the German Research Foundation (DFG, grant FOR2107 DA1151/5-1 and DA1151/5-2 to UD) and the Interdisciplinary Center for Clinical Research (IZKF) of the medical faculty of Münster (grant Dan3/012/17 to UD).

**FSL_Rome:** This study was supported by grants (RC10-11-12-13-14-15/A) from the Italian Ministry of Health and by the ERANET NEURON from the European Commission.

**MPRC:** Support was received from NIH grants U01MH108148, 2R01EB015611, R01MH112180, R01DA027680, R01MH085646, P50MH103222 and T32MH067533, a State of Maryland contract (M00B6400091) and NSF grant (1620457).

**OLIN:** The Olin study was supported by NIH grants R37MH43375 and R01MH074797.

**PAFIP:** The PAFIP study was supported by Instituto de Salud Carlos III, MINECOSAF2013-46292-R, PSYSCAN (Exp.: HEALTH.2013.2.2.1-2_Grant agreement no. 603196), FIS PI14/00639. We want to particularly acknowledge the patients and the BioBankValdecilla

(PT13/0010/0024) integrated in the Spanish National Biobanks Network for its collaboration. We thank IDIVAL Neuroimaging Unit for its help in the technical execution of this work.

**PENS:** This work was supported by the awards by the Department of Veterans Affairs Clinical Science Research and Development Service (Grant No. I01CX000227 [to SRS]) and the National Institute of Mental Health of the National Institutes of Health (NIH) (Grant No. R01MH112583 [to SRS]). Thiswork is also supported by the National Institute of Neurological Disorders and Stroke of the NIH (Grant No. P30 NS076408), the National Eye Institute of the NIH (Grant No. P30 EY011374), the National Institute of Biomedical Imaging and Bioengineering of the NIH (Grant No. P41EB015894), and the NIH (Grant No. 1S10OD017974-01). The content is solely the responsibility of the authors and does not necessarily represent the official views of the Department of Veterans Affairs or the NIH.

**PHCP**: This work was supported by the National Institutes of Health (U01MH108150).

**SCORE:** This study was supported in part by grant 3232BO_119382 from the Swiss National Science Foundation. We thank the FePsy (Frueherkennung von Psychosen; early detection of psychosis) Study Group from the University of Basel, Department of Psychiatry, Switzerland, for the recruitment of the study participants. The FePsy Study was supported in part by grant No. SNF 3200$^{-05}$7216/1, ext./2, ext./3.

**Singapore**: This work was supported by research grants from the National Healthcare Group, Singapore, and the Singapore Bioimaging Consortium research grants awarded to K.S.

**SWIFT:** This research was supported in part by the Swiss National Science Foundation (grant no. 320030_146789).

**UCISZ:** The UCISZ study was supported by the National Institutes of Mental Health grant number R21MH097196 to TGMvE. UCISZ data was processed by the UCI High Performance Computing cluster supported by Joseph Farran, Harry Mangalam, and Adam Brenner and the National Center for Research Resources and the National Center for Advancing Translational Sciences, National Institutes of Health, through Grant UL1 TR000153. AB was also supported by NIH grants 5R01 MH61603, and 2R01MH058251; JF by NIMH (R01 MH-58262).

**UPENN:** The UPENN study was supported by National Institute of Mental Health grants MH064045, MH60722, MH019112, MH085096 (DHW), and R01MH10770 (TDS).
